# Gene functions determine stochastic or adaptive futile transcription in cancers through deregulated start site deployments

**DOI:** 10.64898/2025.12.22.696121

**Authors:** Aditi Lakshmi Satish, Praveen Kumar, Ishani Morbia, Umashankar Singh

**Author notes:** Corresponding author, Umashankar Singh –.

## Abstract

**Background:** Cancers display near–universal hallmarks, including energy addiction, aerobic glycolysis and biosynthesis. In addition to glucose (and glutamine) addiction, cancers also display futile transcription. Discretely measurable futile transcription occurs between transcription start sites (TSSs) and start codons of protein–coding genes. Multiple TSSs for each gene offer various combinations of futile transcription with no effect on the encoded proteins. The relationship between proximal versus distal TSS deployment, futile transcription and energy addiction of cancers remain unclear.

**Methods:** By analyzing FANTOM CAGE data we show that TSS deployment dysregulation in cancers increases the energy cost coefficients of cancer transcriptomes. We define the bases of the altered energy cost coefficients of cancer transcriptomes by comparing TSS deployment frequencies, associated entropies and futile transcription distances across heterogeneous pools of cancers and non–cancers.

**Results:** We show that TSS deployment entropies differ between cancers and non–cancers. It leads to an overall higher distal TSS deployment in cancers but selectively favors an energy– economical proximal TSS deployment for cancer hallmark genes involved in cell proliferation and biosynthesis. We show that the frequency of TSS deployment is linked to the futile transcription distance and gene function differently in cancers and non–cancers.

**Conclusions:** This work lays out a theoretical framework describing stochasticity of TSS deployment in the context of cancerogenesis and energetics of transcription. It also suggests that the current human TSS landscape has evolved to minimize futile transcription, an outcome favored by stochastic TSS deployment. Under normal conditions of energy metabolism they form a threshold which incipient cancer cells breach.

**Highlights:**

⍰ Stochastic TSS deployment favors transcription from proximal TSSs.
⍰ Proximal or distal TSS deployment depends on gene function.
⍰ Cancers generally deploy distal TSSs at a high resource cost.
⍰ Key cancer hallmark supporting genes shift to proximal TSSs even in cancers.

## 1. Introduction

Cellular transformation is at the molecular core of cancerogenesis. A concerted deregulation of multiple checkpoints over a long period of time leads to the manifestation of cancer (1). The outcome of multiple cell–transforming perturbations depends on various factors such as cellular epigenotype, genotype, stemness, macro– and micro–environmental influences. Remarkably, over and above these heterogeneities, the path to cellular transformation and its eventual manifestation as a disease is common across cancers to the extent that near– universal hallmarks for cancer have been prescribed (2). These hallmarks of cancer famously include loss of tumor suppressor checkpoints, oncogenic drive, immortalization and escaping of programmed cell death as core features for which distinct sets of genes have been identified (2–5). It can be argued that the universal nature of cancer hallmarks is due to the primordial and highly conserved nature of molecular pathways which guide some of these fundamental normal cellular processes. Some of the first set of universal hallmarks of cancers to be described were perturbations in well–conserved processes, including cell proliferation (cell cycle), contact inhibition (cell migration and cell–cell communication) and Warburg effect (energy metabolism and biosynthesis). These features of cancer which are deviations from nature’s law of “optimization and least action” can be justified only if they present evolutionary advantages to an incipient cancer cell leading to their selection during tumorigenesis. With the advent of modern techniques deeper insights into cancerogenesis have been obtained (5,6). We now understand the molecular nature of genetic and epigenetic changes, their heterogeneities in a tumor cell mass and their influence on gene expression (5–8). Altered gene expression is the ultimate bottleneck through which mutations and epimutations alter the cellular fates, contributing to the manifestation of hallmark features and the multi–step process of tumor evolution.

Altered gene expression is one of the most well–studied phenomena in cancers. A lingering question in such studies has been the choice of appropriate controls for comparative gene expression analyses. Despite such limitations, several reproducible and predictable patterns of gene expression deregulation have been discovered in cancers that transcend the heterogeneities of cancer and normal sample types (9–15). The molecular outcomes of common oncogenic, tumor–suppressor and restriction checkpoint mutations/epimutations are often traceable to a downstream transcription regulator, often transcription factors, and associated gene expression. A common feature of gene expression deregulation in cancers is the transcriptional addiction of cancer cells (6,16,17), evidenced by glutamine addiction (18), high RNA Pol2 activity (6,19) and biosynthetic bias in energy metabolism (20). Additional layers of transcription deregulation involve post–transcriptional gene silencing in the form of altered lengths of untranslated regions and microRNAs (21–25). As such, transcription, the first step in gene expression, seems to integrate metabolism and energy addiction, mutations, epimutations and cellular environments.

Resistance against gene expression deregulation seems to be the key barrier to be breached for any event to exert its phenotypic effects stably. For an evolving tumor, the mutations, epimutations, and PTGS can be viewed as different strategies to achieve the same outcome, that is, gene deregulation. Regulation of transcription is a primary mechanism to control RNA synthesis, the first step in gene expression. The regulatory element landscape of the human genome and its tight epigenetic modulation plays a key role in determining the transcriptional outcome (26). Evidently, specific DNA sequence features exist in promoter regions. Also, mutations and epimutations in promoter regions are well established to contribute to cellular transformation and cancerogenesis (8,13,26,27). The identity of what constitutes a promoter and levies a regulation on genes is fundamentally determined by the transcription start site (TSS). The location of the TSS gives rise to totally different outcomes for non–coding transcripts or protein–coding mRNA. Given the vastness of our genome as compared to the small number of protein–coding genes, there are immense possibilities for multiple TSSs for each gene. It is established that there are multiple TSSs for most genes. In some cases, this leads to alterations in splicing and reading frames (28–35). However, for most protein–coding genes, the choice of TSSs does not impact the final protein product they code for. However, it is not fully understood how TSSs are selected or deployed. While different sequence features are expected to bestow different transcription starting capacities on different TSSs, it remains unknown how exactly a TSS selection is made during the process of transcription (36–39). The current understanding is that TSS deployment is largely stochastic, as supported by experimental and computational studies (32,40–47).

However, for a “no difference” in the encoded protein, deployments of proximal or >1 kb TSSs entail very different energy costs. The understanding that such a crucial cell fate– determining energy–intensive process is outsourced plainly to stochasticity pits itself against the evolutionary selection of >1 kb TSSs. Moreover, the incentive underlying the choice of one TSS over the other for the same gene and encoded protein remains unaddressed, especially when a distal TSS is preferred over a proximal one. The functional implications of the overall stochasticity of transcription can be better understood by separately addressing its two major dimensions: the frequency of transcription and TSS selection. The latter may not be significant if the TSSs are clustered tightly such that local DNA sequence, as well as chromatin neighborhoods, do not distinguish the various TSSs in a cluster, and stochasticity prevails in selecting TSSs in a cluster with no preference (48). However, when alternative TSSs are located kilobases apart, their different chromatin neighborhoods add a regulatory layer to their otherwise stochastic deployment; co–compartmentalization is established only for different co–expressed genes (49–52), not for different TSSs of the same gene. It is implicit that transcription from >1 kb TSSs would be resource intensive, which shall be justified by the regulatory advantages its deployment provides to a healthy cell.

Long–distance upstream transcription beyond an unavoidable length (minimal length of DNA required for Pol2 positioning and subsequent firing) is futile unless the regulatory benefits offset the overall resource and energy cost (hitherto referred to as “energy cost”). Each base transcribed is an avoidable drain on cellular energy resources. The net energy cost of upstream transcription can be assumed to be a function of upstream region length (distance between TSS and start codon) and transcriptional activity (frequency of deployment of the TSS). The energy costs of futile transcription must pose a challenge to the usability of long– distance upstream TSSs, which gets compounded by alternative translation initiation possibilities due to long 5′UTRs (53–56). In addition, the regulatory features of the TSS–start codon region are likely to suffer a higher mutational load proportional to their lengths (21,57,58). This mutational load disadvantage is only likely to resist dependence on long– distance upstream TSSs over long periods of time. In contrast, the energy costs of transcription from long–distance TSSs are likely to act as an immediate impediment. Thus, the energy costs of transcription would act as a barrier to the usage of long–distance TSSs in cells which are normally sensitive to the fundamental premise of energy conservation.

We use this energy conservation argument to hypothesize that regulatory mechanisms in non–cancer cells, regardless of their cell types, developmental lineages and other variabilities, resist long–distance futile transcription. Transformed cells, derived from cancers or transformed *in vitro*, have deregulated energy conservation mechanisms due to which this resistance against futile transcription gets overridden. We analyze curated FANTOM CAGE data (32,59,60) to test this hypothesis using a simple energy parameter based on the frequency of TSS usage and the distance of futile transcription. We report that disruption of transcriptional energy cost minimization is a feature of cancers in general. Our results demonstrate that the (i) genome presents an energy–conserving design of TSS localization relative to the start codons, (ii) through predominantly stochastic mechanisms the normal (non–cancer) cells utilize this property of the TSS landscape to minimize futile transcription, (iii) cancers exhibit a bipolar deviation from this scheme resulting in an increased transcription–associated energy–cost equivalent. This bipolar deviation from the normal energy–conserving design of TSS deployment is apparently to meet the adaptive needs of rapidly proliferating cells. At one end, the genes linked with cancer hallmarks of enhanced biosynthesis and cell proliferation undergo a high rate of deployment from very proximal TSSs. This tends to outdo the energy–conserving nature of TSS deployment observed even in non–cancers. On the other extreme, an increased transcription from the highly distal TSSs adds to the transcriptional energy cost equivalent through futile transcription. As a sum total the cancers maintain a promoterome more expensive than that of the non–cancers through altered TSS selection. The association between gene function and preference for proximal or distal TSSs shows the adaptive advantage cells derive by switching TSS. We argue that transcription cost coefficient minimization is a property of non–cancers lost in cancers near– universally akin to being a cancer hallmark. Transcriptional energy cost optimization appears to be a force behind the evolution of TSSs, their deployment patterns and is constantly under feedback from biosynthetic and proliferative needs of the cells.

## 2. Materials and methods

### 2.1. Curation of FANTOM CAGE data

FANTOM5 hg38 promoterome CAGE data containing TSS coordinates for 1824 human samples as TPM, were downloaded from UCSC (https://hgdownload.soe.ucsc.edu/gbdb/hg38/fantom5/) in BigWig format. Each sample had two BigWig files to represent the TSSs mapped to the forward strand (fwd) and reverse (rev) separately, both of which were downloaded and processed similarly. The CAGE TPM track (read counts normalized to tags per million) was used as the representative of the TSS activity (CAGE raw score - gamma). The BigWig files were first converted to Wig files and then to BED files. Sample names that contained any of the following keywords: “cancer”, “tumor”, “oma”, “leukemia” were labelled “Cancers (*C*)”. To avoid any confounding effects, we have excluded all embryonic stem cells and embryoid body cells from the analysis. The remaining samples were regarded as “Non–cancers (*N*)”. This keyword search based classification was further manually curated to prevent false classifications. The CAGE data classification provided 1255 non-cancers and 476 cancers with corresponding TSS coordinates in BED format. The BED files from the two strands for the same sample were then concatenated for the analyses.

### 2.2. Gene and TSS annotation of CAGE data

The generated BED files had the TSS data as 2–base window coordinates on hg38 along with the TSS ID, TPM scores and the strand. However, the TSS IDs were labelled *id_n*, where *n* refers to the row number and devoid of gene identity. The original TSS IDs (denoted as “*pn@GENE*” where *n* is a natural number and gives each TSS a numerical within-gene identity and GENE is the official gene symbol (NCBI Gene)) was available for the consolidated CAGE data in three different databases. The hg38 CAGE robust peaks data from UCSC (https://hgdownload.soe.ucsc.edu/gbdb/hg38/fantom5/hg38.cage_peak.bb) was used to map the TSS IDs. However, some of the TSS IDs contained a unique label that were retrieved from the Riken FANTOM5 hg19 robust peaks (https://fantom.gsc.riken.jp/zenbu/gLyphs/#config=FANTOM5_promoterome_hg19). We also used Riken FANTOM5 hg38 (https://fantom.gsc.riken.jp/zenbu/gLyphs/#config=FANTOM5_promoterome_hg38) to create a custom liftover across the genome assemblies. The TSS IDs retrieved from the custom liftover were incorporated into each sample data.

Some of the TSSs (with the same TSS ID) had multiple entries within a sample with overlapping coordinates. To ensure non-redundancy and complete utilisation of the available data from CAGE, the coordinates of the overlapping 2–base TSS windows were merged, and their scores were summed up to generate a single aggregate score. Samples were then concatenated as columns by their sample identity *N* or *C* (see section 3.1 for sample classification) to produce the CAGE raw score matrix. Only those TSSs common to both *N* and *C*, represented by at least one sample in each, were included in the CAGE matrix. The final matrices had the dimensions 55048 × 1255 and 55048 × 476 in *N* and *C* respectively.

### 2.3. Estimating TSS distance (d) and its derivatives

For each annotated TSS of a protein–coding gene in the CAGE profiles, the corresponding most upstream start codon was identified using GENCODE v45 (https://ftp.ebi.ac.uk/pub/databases/gencode/Gencode_human/release_45/gencode.v45.annotation.gtf.gz) keeping in consideration the strand identity. Genes annotated as antisense, non– coding and those from the pseudoautosomal regions were excluded from the analysis. The distance between the TSS and the start codon was calculated independently for protein– coding genes on either strand. The distance for the forward strand TSSs was measured as the absolute difference of the start coordinate of the TSS from the start codon start site and on the reverse strand as the absolute difference of the start codon end coordinate from the TSS end coordinate wherein the coordinates were all from the plus strand.

For the 5’ UTR benchmarking, we used hg38 *knownGene* GTF file from UCSC (https://hgdownload.soe.ucsc.edu/goldenPath/hg38/bigZips/genes/hg38.knownGene.gtf.gz). The GTF file was filtered to get only the 5’ UTR coordinates. The length of the region was used as the distance of the 5’ UTR.

#### 2.3.1. Gene-level distance (*HMd*)

The gene–level distance calculated from all TSSs of a gene is a common value in both *N* and *C* since it is derived from the genome.

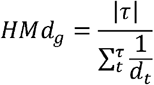

Where,

*HMd*_*g*_ is the harmonic mean of *d* for a gene *g*.

*d*_*t*_ is the distance of TSS *t* from the ATG of its gene *g*.

*t* is a TSS in the gene-space *g*.

*τ* is the set of all TSSs in the gene-space *g*.

| *τ* | is the number of TSSs in the gene-space *g*.

#### 2.3.2 Incremental *d*

Incremental *d* was calculated for each gene as a ratio of the *d* difference between two consecutive TSSs of a gene and the lower *d* in the pair.

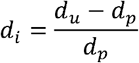

Where,

*d*_*i*_ is the incremental *d* for the pair.

*d*_*u*_ is the distance of the longer TSS of the pair.

*d*_*p*_ is the distance of the TSS immediately downstream to *d*_*u*_.

#### 2.3.3. Fractional *d*

The share of a TSS’ *d* among all the TSS’ of that gene.

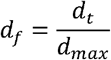

Where,

*d*_*f*_ is the fractional *d* for a TSS *t* in its gene–space.

*d*_*t*_ is the actual *d* for the TSS *t*.

*d*_*max*_ is the *d* of the most distal TSS in the same gene–space.

The *d* and its direct derivatives - incremental *d*, fractional *d* and harmonic mean of *d* (*HMd*) are inherent features of the genome and hence common to all the samples independent of their identities.

### 2.4. Normalizing and scaling of CAGE scores

The raw CAGE matrices were either normalised or scaled according to the analyses. Normalization of a TSS in a sample was done as the share of that score in the entire promoterome independently for *N* and *C*.

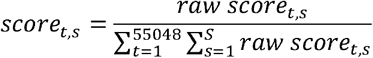

Where,

*score* _*t,s*_ is the normalized score of TSS *t* in sample *s*.

*raw score* _*t,s*_ is the raw score of the TSS *t* in sample *s*.

*t* is the TSS.

*s* is a sample in the sample set *S*.

*S* is the cancer or non–cancer sample set.

Scaling, however, was performed commonly for both *N* and *C*. A scaling factor was estimated for each sample such that the sum of scores for each sample would be equal after scaling.

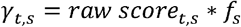

Where,

*γ*_*t,s*_ is the scaled score of TSS *t* in sample *s*.

*raw score*_*t,s*_ is the raw score of TSS *t* in sample *s*.

*f*_*s*_ is the scaling factor for sample *s*.

*t* is the TSS.

*s* is the sample in the sample set *S*.

*S* is the cancer or non–cancer sample set.

Note: All parameters derived from the raw score, normalized score or scaled scores are dependent on the sample set of the parameter. All the parameters described below would therefore contain a value in *N* and one in *C*. The sample (*s*) or sample set (*S*) identities, although not mentioned for brevity, are implied unless mentioned otherwise.

The TSS identities and CAGE scores are summarised in table S5 and the full data is available upon request.

### 2.5. Transcript Count Index (TCI) of a TSS

The *TCI* is a population parameter that is calculated as the share of TSS’ score across all the samples in the promoterome of that sample set, scaled by a billion. Since the scores are normalized, the *TCI* is simply calculated as the sum of normalized scores for a TSS across all samples.

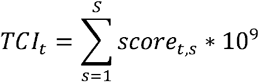

Where,

*TCI*_*t*_ is the transcript count index of TSS *t* in the sample set *S*.

*score*_*t,s*_ is the normalized score of the TSS *t* in sample *s*.

*s* is a sample in the sample set *S*.

*S* is the cancer or non-cancer sample set.

A total of 55048 *TCI*s (one for each TSS) was obtained for *N* and *C* each.

The maximum and minimum *TCI* at every *d* were calculated as *TCI*_*max*_ and *TCI*_*min*_ respectively.

### 2.6. Energy cost coefficient of transcription (E)

We summarise the cost of transcription of a TSS or its parent gene in the following ways:

#### TSS-level E *(E*_*t*_)

The TSS–level *E* is a population parameter for each TSS from all its samples in a population. It is calculated as the number of times the TSS’ d is transcribed.

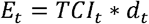

Where,

*E* is the cost of transcription from the TSS *t*.

*TCI* is the transcript count index of the TSS *t*.

*d*_*t*_ is the distance of the TSS *t* from the ATG of its corresponding gene.

#### Gene-level E *(E*_*g*_)

The gene–level *E* is a population parameter for each gene from all its samples in a population. It is calculated as the total energy cost of transcribing a gene from each of its TSSs.

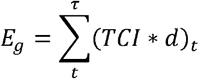

Where,

*E*_*g*_ is the total cost of transcribing the gene *g* from all its TSSs.

*t* is a TSS in the gene–space *g*.

*τ* is the set of all TSSs in the gene–space *g*.

*TCI* is the transcript count index for the TSS *t* in the gene–space *g*.

*d*_*t*_ is the distance of the TSS *t* from the ATG of its corresponding gene *g*.

### 2.7. Base Count Index (BCI)

The population parameter *BCI* for each TSS was calculated as the share of *E* of a TSS across all samples within the sample set, scaled by a billion.

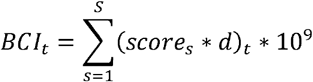

Where,

*BCI* _*t,S*_ is the base count index of TSS *t* in population *S*.

*score*_*t,S*_ is the normalized score of the TSS *t* in sample *s*.

*d*_*t*_ is the distance of the TSS *t* from the ATG of its corresponding gene *g*.

*s* is a sample in the sample set *S*.

*S* is the cancer or non–cancer sample set.

### 2.8. Effective d (*v*d)

The effective *d* can be both a sample–level and population level value for each gene. The effective *d* at population level is calculated as the *TCI*–weighted arithmetic mean of *d* while at the sample level, it is the score–weighted arithmetic mean of *d*.

#### 2.8.1. Population–level effective *d*

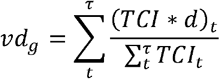

Where,

*vd*_*g,S*_ is the effective distance of gene *g* in population *S*.

*TCI* _*t,S*_ is the transcript count index of TSS *t* in population *S*.

*d*_*t*_ is the distance of TSS *t* from the ATG of its gene *g*.

*t* is the TSS in the gene–space *g*.

*τ* is the set of all TSSs in the gene–space *g*.

*S* is the cancer or non–cancer sample set.

#### 2.8.2. Sample–level effective *d*

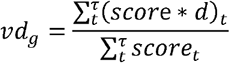

Where,

*vd*_*g*_ is the effective distance of gene *g* in sample *s*.

*score*_*t*_ is the normalized score of TSS *t* in sample *s*.

*d*_*t*_ is the distance of TSS *t* from the ATG of its gene *g*.

*t* is the TSS in the gene–space *g*.

*τ* is the set of all TSSs in the gene–space *g*.

*s* is the sample in sample set *s*.

*S* is the cancer or non–cancer sample set.

Note: Effective *d* is always assumed at the population–level in this work except when performing the PCAs.

### 2.9. Gene–space entropy (gH)

The gene–space entropy was calculated only for those genes which had at least two TSSs. We define the gene–space through the probability of a TSS’ activity across all samples of the population with respect to the activities of all the TSSs of that gene.

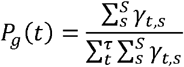

Where,

*P*_*g*_ (*t*) is the probability of occurrence of the TSS *t* in its gene–space *g*.

*γ*_*t,s*_ is the scaled score of the TSS *t* in sample *s*.

*t* is a TSS in the gene–space *g*.

*τ* is the set of all TSSs in the gene–space *g*.

*s* is a sample in the sample set *S*.

*S* is the cancer or non–cancer sample set.

We then derive the gene–space entropy as the Shannon entropy of the probabilities of all the TSSs of the gene. Shannon entropy measures the information in bits or binary transfer of information; The formula for entropy thus uses a base 2 in the logarithm. The gene–space, however, allows only one of the *τ* set of TSSs to be deployed at any point of time. Thus, given that the gene is transcribed, it can be transcribed in as many ways as the number of TSSs of that gene (also represented as |*τ* |). The gene–space entropy thus uses a logarithm base equal to the number of TSSs contributing to that gene–space.

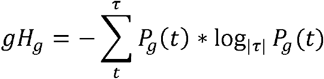

Where,

*gH*_*g*_ is the gene–space entropy for a given gene *g* in sample set *S*.

*P*_*g*_ (*t*) is the probability of deployment of TSS *t* within its gene–space *g*.

*t* is the TSS in the gene–space *g*.

*τ* is the set of all TSSs of the gene–space *g*.

### 2.10. Sample–space entropy (sH)

Stochastic deployment of TSSs in the *N* and *C* populations across CAGE profiles was calculated using Shannon entropy for the sample–space of the TSS. Since entropy is a measure of the probability of occurrence, and the CAGE scores for a TSS is a direct measure of the number of times the TSS was deployed, we used the ratio of the scaled scores and number of samples in which the TSS was deployed as the probability equivalent in the sample–space entropy calculation. Since some TSSs could be represented only by a small proportion of the samples and there is an unequal sample count in the *N* and *C* sample sets, we considered only those TSSs that had a CAGE score in at least 100 non–cancer and 100 cancer samples.

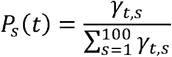

Where,

*P*_*s*_ (*t*) is the probability of occurrence of the TSS *t* in its sample–space.

*γ*_*t,s*_ is the scaled score for the TSS *t* in sample *s*.

*t* is a TSS.

*s* is a random sample in the sample set *S*.

*S* is the cancer or non–cancer sample set.

The sample–space was then calculated as simply the Shannon entropy with a log–base 2. Since the sampling involves a randomization, we estimate the sample–space entropy as the arithmetic mean of the sample–space entropy calculated through 10 independent iterations of random sampling.

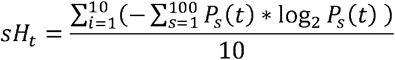

Where,

*sH*_*t*_ is the sample–space entropy for TSS t in the sample set *S*.

*P*_*s*_ (*t*) is the probability of occurrence of the TSS *t* in its sample–space.

*t* is a TSS.

*s* is a randomly derived sample in the sample set *S*.

*i* is the iteration for the random draw.

This mean sample–space Shannon entropy from the 10 iterations shall henceforth be referred to as the sample–space entropy of the TSS. Some TSSs have such poor deployment frequencies across samples that the random draw could produce all samples with a 0 score. We termed these TSSs as “non–frequently deployed” and the remaining as “frequently deployed”. The frequently deployed TSSs were further used to generate a new set of entropy values as described earlier except with only the non–zero scoring samples in *N* and *C* instead of all samples for the 100 randomizations. This filtering of samples was done independently for each TSS in *N* and in *C*.

### 2.11. Compound entropy (cH)

We use the product of the TSS probabilities in the sample–space and the gene–space to define the compound probability for that TSS. The gene-space probabilities are calculated as described in the gene–space entropy, however, the sample–space probabilities were calculated using all samples within *N* and *C* instead of just 100 random samples.

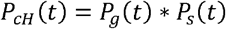

Where,

*P*_*cH*_ (*t*) is the compound probability of TSS *t*.

*P*_*g*_ (t) is the gene–space probability of TSS *t*.

*P*_*s*_ (*t*) is the sample–space probability of TSS *t*.

*t* is a frequently deployed TSS.

The compound entropy was then calculated for each gene similar to the gene–space entropy – with a log–base equal to the number of TSSs in that gene.

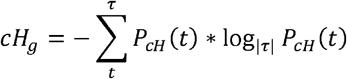

Where,

*cH*_*g*_ is the compound entropy for a gene *g* in sample set *S*.

*P*_*cH*_ (*t*) is the probability of deployment of TSS *t* within its gene–space *g*.

*t* is the TSS in gene–space *g*.

*τ* is the set of all TSSs of the gene–space *g*.

|*τ*| is the number of TSSs in the gene.

*S* is the cancer or non–cancer sample set.

### 2.12. TSS equalization simulation

This simulation was done to equalize TSSs activity (*TCI*) per gene iteratively for *N* and *C*. For each gene, the TSSs were first sorted in ascending order of their *d*. Iteratively, consecutive TSS activities were replaced by their means, thus equalizing their corresponding *TCI* s independently in *N* and *C*. The total number of iterations of *TCI* equalizations were governed by the maximum TSS count for a gene. In our data, it was the *SORBS2* gene with 39 TSSs and thus 38 iterations of equalizations (plus one non–equalizing zeroth iteration) were done to exhaust all TSSs. Those genes with lesser number of TSSs exhaust all TSSs much before this iteration and do not change their values. After every iteration, the gene– space entropy and the gene–level *E* were calculated and compared between *N* and *C* through a gene–paired Wilcoxon test. The simulation was done twice: first in the ascending order of *d* per gene (most proximal TSS to the most distal TSS) and second in the descending order of *d* per gene (most distal TSS to the most proximal TSS).

### 2.13. Promoterome entropy maximisation simulation

A simulated promoterome was generated with an equal number of random samples from *N* and *C*, and completely entropized samples. The promoterome starts as a matrix of random samples of *N* and *C* scaled scores only. With every iteration, one sample each of *N* and *C* was replaced by completely entropized samples. Over 50 iterations, the simulated promoterome shifted from completely *N* and *C* to completely entropized. A completely entropized sample was generated using the mean of scaled scores of any sample. All TSSs of this sample thus have the same score. In every iteration, the simulated promoterome was used to calculate the energy cost, *gH, sH* and *cH*. Since the sample selection involved randomization, 10 sub– iterations were performed for each iteration and the mean of each of the parameters *E*_*t*_, *gH,sH* and *cH* was calculated as a single value per iteration. The TSSs were then segregated by the *d*–bins: 0–1 kb, 1–5 kb, 5–10 kb, 10–50 kb and >50 kb. For each *d*–bin, an exponential curve was fit for the 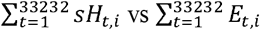 data for every iteration. The *sH*– *E*_*t*_ relationship was best described by the exponential growth function with rate constant parameter 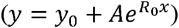 for all bins except the first where a correlation could not be interpreted (Table S15). The actual mean *sH* for each *d*–bin obtained from the *N* and *C* data was used to interpolate the corresponding *E*_*t*_. The curve fitting and interpolation was done using Origin 2017.

### 2.14. Promoterome substitution simulation

A simulated promoterome was constructed with 100 randomly selected cancer samples. With every iteration the cancer samples were replaced with an iteration count number of non– cancer samples. Each iteration involved 10 sub–iterations where the sample–space entropy and energy cost was recorded per TSS and the mean of the sub–iterations was used as the representative value per iteration. The TSSs were split by the distance binned by 0.1kb or 1kb and plotted to compare *sH* and *E*_*t*_ for every iteration.

### 2.15. Functional category enrichment analyses

All functional category enrichment analyses were performed on GOrilla. The terminal quartiles (Q1 and Q4) of the *C*/*N* ratios for each of the parameters (*vd,gH,sH* and *vd/cH*) sorted were used to produce two gene sets. Gene Ontology (GO) biological process (BP) and molecular function (MF) term enrichment was performed on these two sets in two different directions of comparison using GOrilla (61,62) at default settings for analyses of non–ranked lists. Both gene sets were analyzed as paired gene lists where one gene set was used as a foreground against the other as the background for the enrichment analysis (foreground/background: Q1/Q4 and Q4/Q1). GO terms that satisfied the p–value criteria were manually assigned a broader functional category (Biosynthesis and metabolism, Cell proliferation, DNA and chromatin organization, Signalling and homeostasis, Development and differentiation, and Others).

### 2.16. Principal Component Analysis (PCA)

We calculated a sample–level effective *d* for each of the 1731 CAGE profiles and 15861 genes (as described in Methods, 2.8.2; *TCI* was replaced by scaled scores for the calculation). PCA was performed on the same data using the R package *PCAtools* and the first three principal components were plotted. The metadata for the PCA annotations were curated manually (see Methods, 2.1) and available in Table S5. The variances associated with the principal components were compared through a Levene’s test from the *cars* package.

### 2.17. Statistics and data visualization

All statistical analyses were performed on R version 4.4.1 (2024-06-14). All randomizations and operations involving randomizations were carried out using the R *base* package of the aforesaid version. All paired tests were performed using the *wilcox*.*test()* function of *base* R package with the argument *paired* set to *T* if the test was paired Wilcoxon signed–rank test or *F* if it were a Wilcoxon rank–sum test. Statistical details of the data presented in the manuscript are summarised in supplementary table S16. The statistics table includes false discoveries calculated as a fraction of bootstrap iterations that a test for a variable gave the same result. For paired data, we swapped the N and C identities for 50% of the data as a measure false positives while for unpaired data, we pooled both groups and randomly sampled a pseudo identity. Plots were generated using the *ggplot2* package of *tidyverse*. The plots corresponding to gene ontology and the substitution simulation were made with Graphpad Prism 10.4.1. The interpolation was done using Origin 2017. All schematics were custom made on Apple Keynote.

## 3. Results

### 3.1. Curation of CAGE and GENCODE data reveals an energy–conserving design of TSS deployment and its deregulation in cancers

The human FANTOM CAGE data (Noguchi et al. 2017) provides TSS coordinates, their CAGE scores as well as their gene affiliations. We combined that with the GENCODE annotations which provide start codon coordinates for each gene. The most–upstream start codon for any gene was designated as the only one to which all TSSs of the same gene would be mapped. This implied that some TSSs which were present downstream of the designated start codon were excluded from our analyses (Table S1). The distance between TSSs (start of the CAGE coordinates) and the start codon was abbreviated as *d* (Fig 1, A and B). The *d* is essentially a count of nucleotides in GRCh38 obtained as a difference in the positions of TSSs and the associated start codons (Table S2). All references to *d* (and its derivatives) henceforth imply the base–count as units. This yielded a total of 57461 TSSs from 16302 protein–coding genes for our analysis with approximately 3 TSSs per gene on average, and at least 76% of the genes had more than one TSS. For a large population of TSSs (60%) the *d* values were 1 kb or less (Fig 1C). The <1 kb *d* tendency was prevalent for the TSSs even when the *d* profiles were calculated per gene as a harmonic mean of *d* (*HMd*, see Methods 2.3.1; Table S3) of the constituent TSSs (Fig 1C). Indeed, an incremental *d* profile (a ratio of the *d* difference between two consecutive TSSs of a gene and the lower *d* in the pair) obtained by a gene–level analysis of TSSs recapitulated the pattern of *d* restriction (Table S2 and Fig S1). Referring to a particular TSS located at *d* for each gene, the consecutive >1 kb TSSs for the majority of genes showed a *d* increment of about 50% with a large population exhibiting a *d* increment restriction between 4% and 0.1% (Fig S1). It follows from these observations that the <1 kb TSS deployment presents an advantage which has been accorded to most if not all genes through the process of evolution.

**Fig 1.**
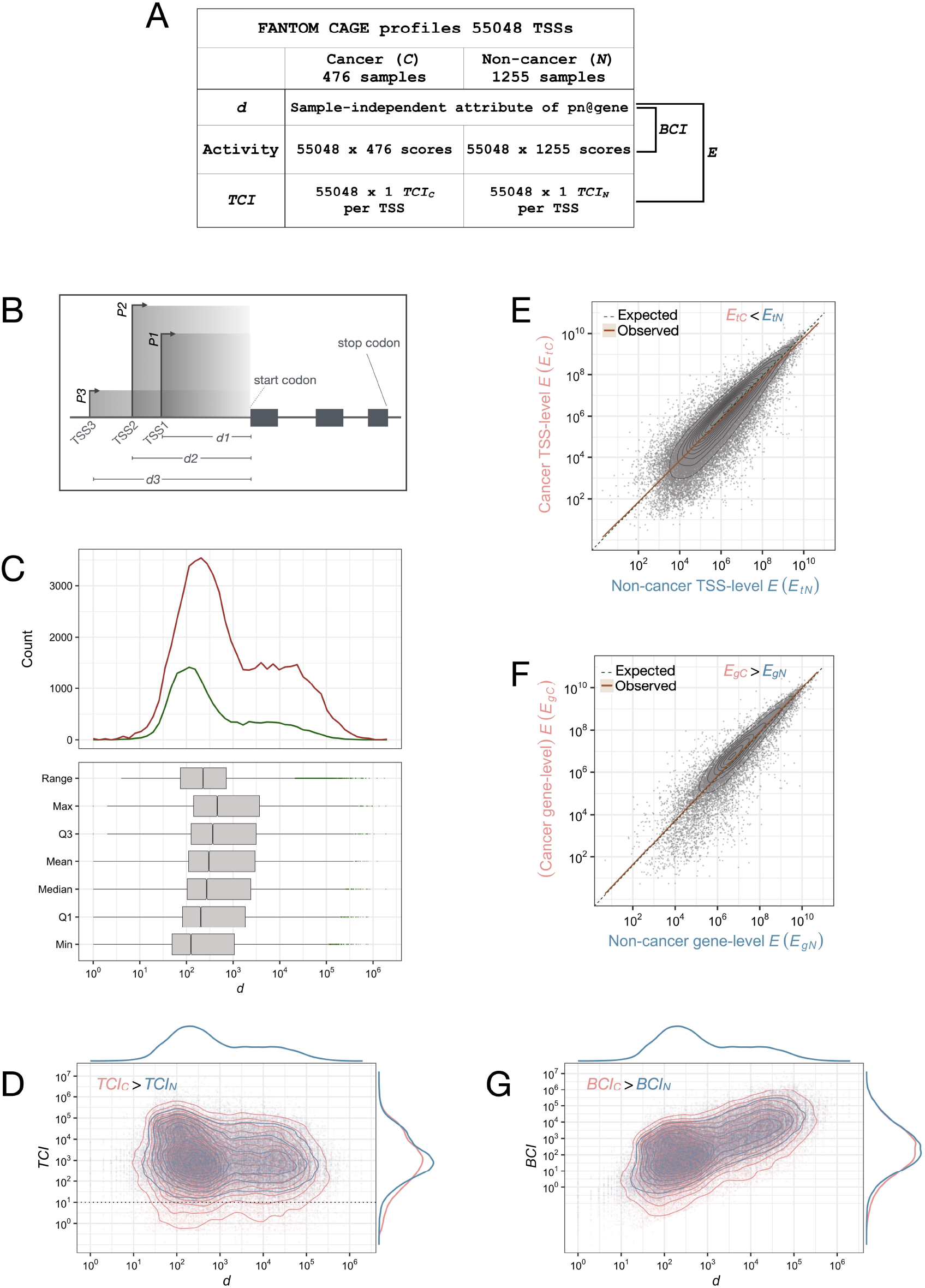
Minimization of the energy cost coefficient of transcription (*E*) by proximation of length transcribed upstream of the start codon (*d*) is an evolved feature of the human promoterome utilized by non–cancers and disrupted in cancers. A: The schematic summarizes CAGE data curation based on GENCODE annotations of start codons for derivation of the parameters *d, TCI, BCI* and *E* and CAGE metadata–based classification of samples into cancers (*C*) or non–cancers (*N*). B: A diagrammatic representation of a hypothetical gene with an ORF marked by start and stop codons and three TSSs with corresponding *d* (*d*1,*d*2,*d*3) and activities *P* (*P1, P2* and *P3*) respectively. The area shaded in grey shows the product *d* * *P* and represents the energy cost coefficient *E* equivalent associated with each TSS. C: The distribution of *d* values for 56412 TSSs (red) shows a proximal concentration of TSSs with majority concentrated in the range 100–1000 bps (X– axis). A gene–level *d* (calculated as a harmonic mean of *d* of all TSSs for a gene, *HMd*) shows that this TSS–level *d* profile is distributed across 16093 genes as well. D: *TCI*, the share of a TSS in the aggregated promoterome from a set of samples classified as cancer (*C*) or non–cancer (*N*), is significantly higher in cancer (Wilcoxon signed–rank test paired by 55048 TSSs, p–value = 7.846e-3). Despite the difference, throughout the spectrum of *d* (X– axis) leaky low–level transcription is visible in *C* (demarcated by the dotted line at *TCI* value 10 on the Y–axis). E and F: TSS level *E* (*E*_*t*_ ) in *C* and *N*, derived from the respective *TCI* s and *d*, are highly correlated but lower in *C* (Wilcoxon signed–rank test paired by 55.048 TSSs, p–value < 2.2e-16). However, the gene–wise bundling of *E* (summing up *E*_*t*_ for a gene into *E*_*g*_) inverts the *E* difference (*E*_*g*_ > *E*_*g*_, Wilcoxon signed–rank test paired by 15861 genes, p–value < 2.2e-16) indicating a relationship between gene function, cancer and energy cost coefficient of transcription. G: A population size–independent parameter base count index (*BCI*), calculated as a share of CAGE *score* * *d* for any TSS in the aggregated promoterome recapitulates the excess futile transcription in *C* over all ranges of *d* (*BCI*_*C*_ > *BCI*_*N*_, Wilcoxon signed–rank test paired by 55048 TSSs, p–value < 2.2e-16).

The *d* values for all the TSSs were benchmarked against the known 5′ UTR lengths (UCSC) (Fig S2). As expected, the *d* values were higher than the UTR lengths due to the non– exclusion of introns in *d* calculation even though for the majority of TSSs the *d* values showed a distribution similar to that of the 5′ UTR lengths (Fig S2). Unlike 5′ UTR lengths however, a substantial number of TSSs showed much higher *d* values, ranging over two orders of magnitude (Fig S2). Such a disconnect between *d* and 5′ UTR lengths showed that despite concentration of TSSs <1kb, the transcriptional regulatory constraints built into the genome enforce transcription start from some start sites located so far from the start codons that they vastly exceed the 5′ UTR length. Thus, the transcription regulation through TSS deployment entails futile transcription not necessarily required for downstream translation regulation.

We hypothesized that if alternative TSS deployment does not affect the downstream translation process then the normal cells would tend to minimize futile transcription and in a “one gene–multiple TSS” system the high– *d* TSSs must offer distinct regulatory advantages to get deployed. Altered energy metabolism distinguishes transformed cells from normal ones and as such the normal energy conserving nature of TSS deployment would decompose in cancers leading to an overall more futile transcription through a dysregulated TSS deployment. To test this hypothesis, we compared CAGE data from non–cancer and cancer samples in multiple different ways.

CAGE samples were classified as non–cancer (*N*) or cancer (*C*) through a curated process (Fig 1A and Table S4) based on the CAGE metadata and the associated scores for all the TSSs were collated (see Methods, 2.2). The representations of TSSs were expectedly non– uniform within, as well as unequal between, the *N* and *C* sample groups. A major fraction of TSSs were consistently detected similarly across >95% of *N* as well as *C* sample groups (Fig S3, A and B). However, a significant subset of TSSs had a low intra–group representation of <5%. The number of TSSs represented in <5% samples was remarkably higher in *C* (11144) than that for *N* (8095). Of these TSSs with low representation, 5741 TSSs were common to both *N* and *C* leaving only 2354 and 5403 TSSs unique to *N* and *C* respectively. Overall, a total of 55048 TSSs were commonly listed in N and C sample groups whereas 1346 and 18 TSSs were exclusive to the *N* and *C* populations respectively. Of the entire spectrum of TSS representation frequency in *N* or *C* only the TSSs with <5% representation showed a significantly higher *d* in *C* than that in *N* (Fig S3C; Wilcoxon rank–sum test, p–value = 8.755e-4). Removing the *N*–exclusive and *C*–exclusive TSSs from the comparison did not affect this finding (not shown). The TSSs deployed differently in *C* and *N* (TSS detected in <5% of either *C* or N) have a sample–type preference for deployment. Stochastic TSS deployment is unlikely to give rise to such a *C–* or *N*–biased TSS deployment profile. Thus, if the sample type–specific low representation frequency of TSSs is a representation of non– randomness with which TSSs get deployed then these data indicated that TSSs with less stochastic deployment in *C* have a relatively higher *d*.

To analyze the overall TSS deployment rates in *N* or *C* (Table S5 and Extended data table: https://usegalaxy.org/api/datasets/f9cad7b01a472135b85e9cab771dc355/display?to_ext=tabular) without any effect of the differences in sample counts, we worked out a population size– independent parameter to represent the relative TSS deployment frequency. The source CAGE data originally from FANTOM5 was processed, as described in methods, is available as extended data. The share of each TSS in the total CAGE score count was separately derived for *C* or *N* sample pools and used as a Transcript Count Index (*TCI*), a TSS–specific parameter representing its net deployment frequency in a sample group regardless of the intrinsic inter–sample variations (Table S6). Overall *TCI*_*C*_ was significantly higher than *TCI*_*N*_ (Fig S4) even when 54.32% TSSs had *TCI*_*C*_ <*TCI*_*N*_ (Wilcoxon signed–rank test paired by 55048 TSSs, p–value = 7.846e-3). The *TCI*_*N*_ and *TCI*_*C*_ showed different relationships (Pearson and Filon’s z, p–value < 2.2e-16) with *d* (Spearman’s correlation coefficient *r* for *TCI*_*N*_ vs *d* = -0.1468 and p–value < 2.2e-16, *r* for *TCI*_*C*_ vs *d* = -0.1369 and p–value < 2.2e-16) (Fig 1D). While low *TCI* was abundant in *C* at all *d* ranges (Fig 1D, dotted line threshold *TCI* of 10), the higher *TCI* in *C* was restricted primarily to a subset of proximal *d* TSSs. A comparison of *TCI*_*max*_ or *TCI*_*min*_ between *N* and *C*, over all the possible *d* values showed two distinct patterns for <1 kb and >1kb TSSs (Fig S5, A and B). The <1 kb TSSs in *C* and *N* both showed a wide range of relatively discrete *TCI*_*max*_ and *TCI*_*min*_ at the extremes with missing intermediate range *TCI* values. At >1 kb TSSs, however the *TCI*_*max*_ and *TCI*_*min*_ showed a continuous distribution. This suggested that a range of transcriptional output is achieved, presumably through a wide range of regulated deployment limited to >1 kb *d* TSSs (Fig S5, A and B). Interestingly, at >1 kb TSSs leaky transcription was detected in *C* as low levels of *TCI*_*max*_ were pervasive over all *d* ranges. Supporting this, across all the *d* values only *TCI*_*min*_ were lower in *C* than in *N* (Wilcoxon signed–rank test paired by 55048 TSSs, p– value < 2.2e-16) while *TCI*_*max*_ did not exhibit any difference. Thus, clearly there are *C* and *N* specific *d*–dependence patterns of TSS deployment. The concentration of *d* proximal to start codons concomitant with a difference in *d*–dependent TSS deployment indicated that the futile transcription could be different between *C* and *N*.

### 3.2. Energy conserving nature of TSS deployment depends on minimization of d

The energy cost of transcription from a TSS can be estimated by combining two parameters it directly depends on: the transcriptional activity for a TSS (represented by *TCI*) and the transcriptional distance from the TSS to the start codon (*d*). The product of a *TCI* and the respective *d* generates an energy coefficient *E*_*t*_ for each TSS. The actual energy costs of transcription from that TSS would be a combination of constitutive energy costs of transcription (TSS–invariant parameters, such as the energy cost of ribonucleotide biosynthesis and RNA polymerase activity, innate to the mechanism of transcription) and *E*_*t*_ (a TSS–variant parameter including the frequency of transcriptional firing represented by *TCI* and *d*; Fig 1B). The net energy cost coefficient of transcription from a TSS could be expressed as a product of *TCI* and *d. E*_*t*_ for *C* and *N* were compared for 55048 TSSs. *E*_*tC*_ was higher than *E*_*tN*_ for 45.68% TSSs and lower than *E*_*tN*_ for 54.32% TSSs with *E*_*tC*_ having a significantly lower value than *E*_*tN*_ (Wilcoxon signed–rank test paired by 55048 TSSs, p– value < 2.2e-16; Fig 1E and S6A). This was unexpected as *TCI* was significantly higher than *TCI*_*C*_ (Wilcoxon signed–rank test paired by 55048 TSSs, p–value < 0.05; Fig S4). Interestingly however, gene–level *E* (*E*_*g*_ ) derived by summation of *E*_*t*_ of all TSSs separately for *N* (*E*_*gN*_ ) and *C* (*E*_*gC*_) for a gene was significantly higher for *C* as compared to *N* (Wilcoxon signed–rank test paired by 15861 genes, p–value < 2.2e-16; Fig 1F and S6B). *E*_*gC*_ was higher than *E*_*gN*_ for 52.23% genes suggesting that the *E*_*t*_ differences between *N* and *C* are not random TSS–specific attributes. Rather, the TSS–level *E* contributes to gene–level *E* thereby connecting transcription energy cost coefficient with gene function.

The results so far showed that widespread alterations in TSS deployment is a feature of transcriptional dysregulation in cancers. A net effect of this altered TSS deployment is an increase in transcriptional resource cost. This would only be possible if the altered TSS deployment in cancers was a net use of larger *d*. Assuming that the *N* and *C* samples from which we analyzed the CAGE data represent the generic TSS deployment properties of non– cancer and cancer cells respectively, we sought to quantitatively test the possibility of increased *d* deployment in cancers. These results were supported by TSS–level paired analysis of consecutive distal–to–proximal deployment ratios using *TCI*. This analysis demonstrated an increased distal TSS utilisation in *C* for every relatively proximal TSS of a gene (Fig S7, A and B). We generated a Base Count Index (B*C*I) from the *N* and *C* sample pools (Table S6) that represents the share of deployment of each TSS in the transcription of *d* out of the total *d* deployment transcription of a sample group. In *E*_*t*_, *TCI* represented the TSS activity as relative to the sample group but *d* remained absolute. In B*C*I, however, the TSS– activity as well as *d* both were combined relative to the sample group. In TSS–paired comparisons *BCI*_*C*_ > *BCI*_*N*_ (Fig S8; Wilcoxon signed–rank test paired by 55048 TSSs, p– value < 2.2e-16) with an overall high variance in cancers (Fig 1G) unlike *E*_*t*_ which was *E*_*tC*_ < *E*_*tN*_ . *E*_*t*_ and *BCI* both have two components, TSS–activity and TSS–*d*.

Such a decomposed comparison of transcriptional resource cost for TSSs independent of their gene affiliations reinforced the importance of futile *d* transcriptional cost, TSS localization with respect to the start codons and their deployment. Through a direct proportionality between B*C*I on one hand, and *TCI* and *d* on the other, the following deductions could be worked out:

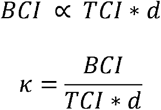

The constant *κ* is a factor which represents systemic *d*–functionalisation. Since the population aggregates of the terms *BCI, TCI* and *d* are identical for both *N* and *C* (see Methods, 2.7, 2.5 and 2.3), the factor *κ* is bound to be different between *N* and *C*. We could calculate *κ* separately for *N* and *C* as follows: *κ*_*C*_ = 2.71991e-4 and *κ*_*N*_ = 2.48027e-4. Thus, the *κ*_*C*_ was 9.6615% higher than *κ*_*N*_ . It followed from these findings that each TSS in cancers is likely to conduct a higher *d*–dependent futile transcription by this relative magnitude.

However, *d* is a real TSS–level parameter encoded in the genome. For each gene a fixed mean *d* could be represented by a harmonic mean of the *d* values of its TSSs (*HMd*) (Fig 1C, see Methods, 2.3.1). TSS deployment (represented by CAGE scores) functionalizes the *HMd* for each gene in a sample–specific manner giving rise to a virtual effective *d* for each gene. The effective *d* for the genes were calculated as a *TCI*–weighted mean *d* (*vd*) of all contributing TSSs (refer to schematic, Fig 2A; Table S3). *HMd* represents an expected effective *d* for a gene if all the TSSs were equally deployed. In contrast, *vd* represents the actual effective *d* which factors in the inter–TSS deployment differences with high *vd* indicating an overall higher preference for distal TSSs for a gene. *vd*, which was higher than *HMd* in *N* as well as *C* (not shown), was also overall higher in cancers (Wilcoxon signed– rank test paired by 15861 genes, p–value = 2.21e-35; Fig S9A). *vd* showed the power to classify individual *C* samples from *N* samples (Fig 2, B–E). Thus, *vd* functioned as a parameter representing the differences between *E*_*gC*_ and *E*_*gN*_ in the context of *d* at the gene– level. If these properties of *vd* are functionally relevant in cancer, we expected functional category enrichment in genes ranked by *vd*. Genes were ranked by *vd*_*C*_ /*vd*_*N*_ ratios and terminal quartiles (Fig S9B) were compared for functional category enrichment (Table S7). The low *vd* quartile, with a decreased *vd* in cancers, was rich in GO BP terms pertaining to biosynthesis, cell proliferation, cell cycle and GO MF terms associated with DNA metabolism (Fig 2, F and G). The upper quartile genes were rich in GO BP terms pertaining to cellular signalling and GO MF terms linked to signal transduction mechanisms (Fig 2, F and G).

**Fig 2.**
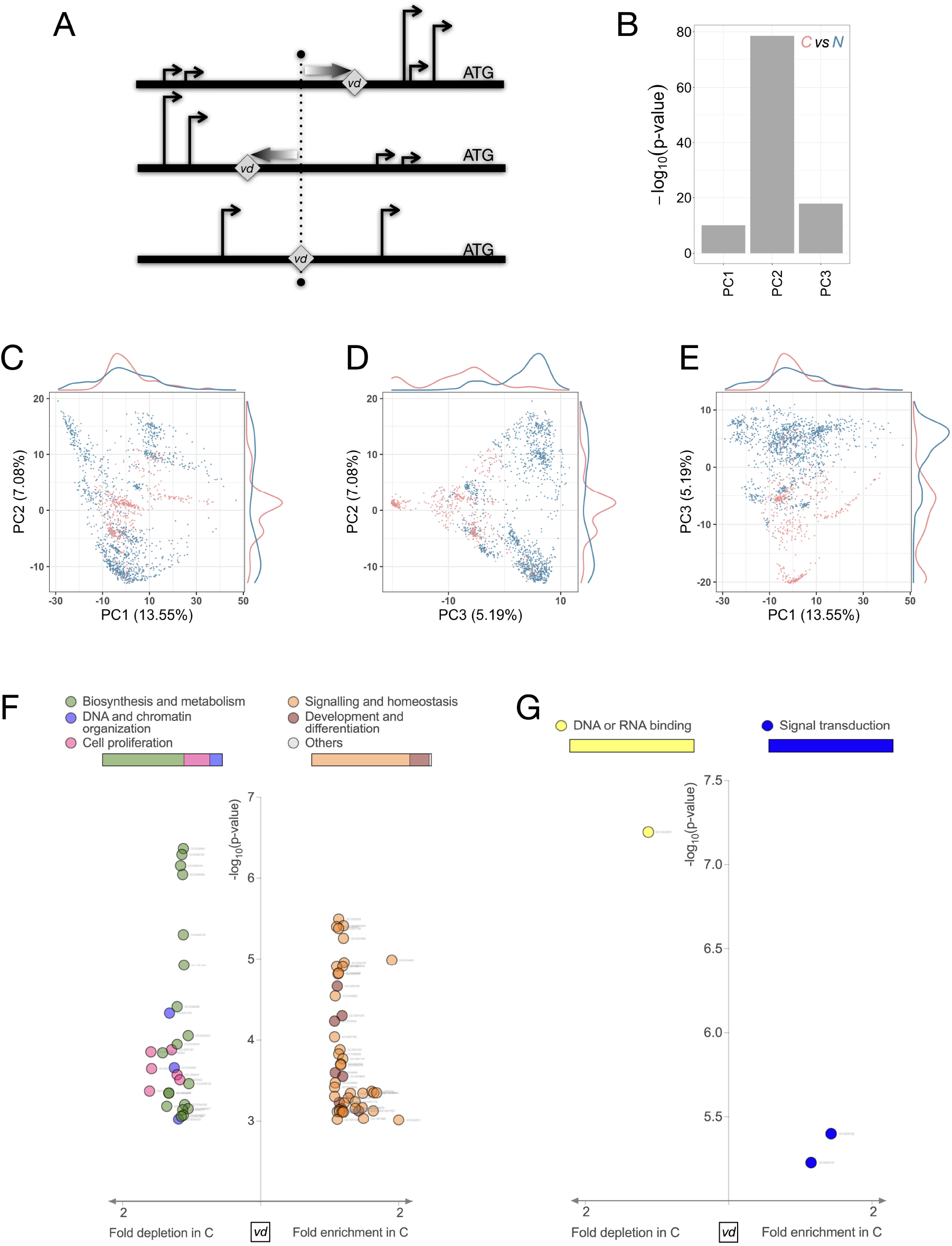
Transcriptional activity (*TCI*)–weighted *d* (*vd*) distinguishes cancers from non– cancers and affects genes according to their cancer–relevant functions. A: A schematic representation of effective *d* (*vd*) using three hypothetical genes with different profiles of TSS locations (vertical arrow heads at different distances upstream of ATG) and activities (heights of the arrowheads). Since *vd* (grey diamonds) is calculated as an activity–weighted *d* from all TSSs for every gene, a *d*–dependent difference in activity would affect the magnitude of *vd*. The lowermost gene exemplifies a simple case where *vd* is equidistant from two equally active TSSs (dotted marker) whereas the other two genes have their *vd* located more proximally (topmost) or distally (middle) due to a combination of *d*, activity and number of TSSs. B–E: Principal component analyses of *vd* value shows clear separation between *C* and *N* despite a large amount of intra–sample group heterogeneity in *N*. Levene’s test of PCA values for the first three components shows a highly significant difference between *C* and *N* (B). Marginal densities along the top and right of the PCA scatter plots highlight the differences between *C* (pink) and *N* (blue). F and G: *vd* difference between *C* and *N* is linked to gene functions. *vd*, although overall increased in *C* over *N*, is maintained lower in *C* for cancer hallmark functions, of which biological processes biosynthesis and cell proliferation (F), and molecular functions DNA or RNA–binding are noteworthy (G). Y–axes show p–value of Fisher exact test for significant enrichment using GOrilla Gene Ontology term enrichment and X–axes show the fold change of *vd* in *C* over *N*. Specific Gene Ontology terms (marked in light grey fonts for every data point and presented in table S7) are merged into color–coded assigned terms presented as legends on the top of the scatter plots.

These results suggested that TSSs are deployed through different *d*–dependent mechanisms in *N* and *C* resulting in different TSS deployment frequencies. This gives rise to a net higher energy cost coefficient of transcription in *C* at the gene level. This deregulation apparently manifests as a pan–cancer feature even if the *N* and *C* sample groups have internal heterogeneities. Stochasticity being a key mechanism of TSS deployment, we next analyzed the stochasticity of TSS deployment in *N* and *C* deducible from the CAGE data and tested if stochastic TSS deployment relates with *d* differently in *N* and *C*.

### 3.3 Gene–space entropies show that mitigation of futile d transcription through stochastic TSS deployment is bypassed in cancers

The genome offers a proximal concentration of TSSs in the gene–space and a stochastic deployment of all TSSs would tend to deploy <1 kb TSSs more than the >1 kb ones for every gene. For any given gene, an alteration in stochastic TSS deployment would affect proximal– versus–distal TSS preferences and a subsequent alteration in *E*. We estimated the stochasticity associated with TSS deployment of a gene using Shannon entropy in the gene– space (*gH*) for the *N* and *C* populations (Table S8, see Methods, 2.9). For any two genes with the same number of TSSs, the gene that deploys all its TSSs more uniformly (strong stochastic deployment) would therefore have a higher gene–space entropy than the gene which deploys TSSs preferentially (strong deterministic deployment) (Fig 3A). We found that the gene–space entropies are similarly distributed in *N* and *C* but the overall *gH*_*C*_ was lower than *gH*_*N*_ (Wilcoxon signed–rank test paired by 11928 genes, p–value = 1.364e-09; Fig 3B) showing that stochastic deployment of TSSs in the gene–space is the dominant mechanism which is resisted in cancers.

**Fig 3.**
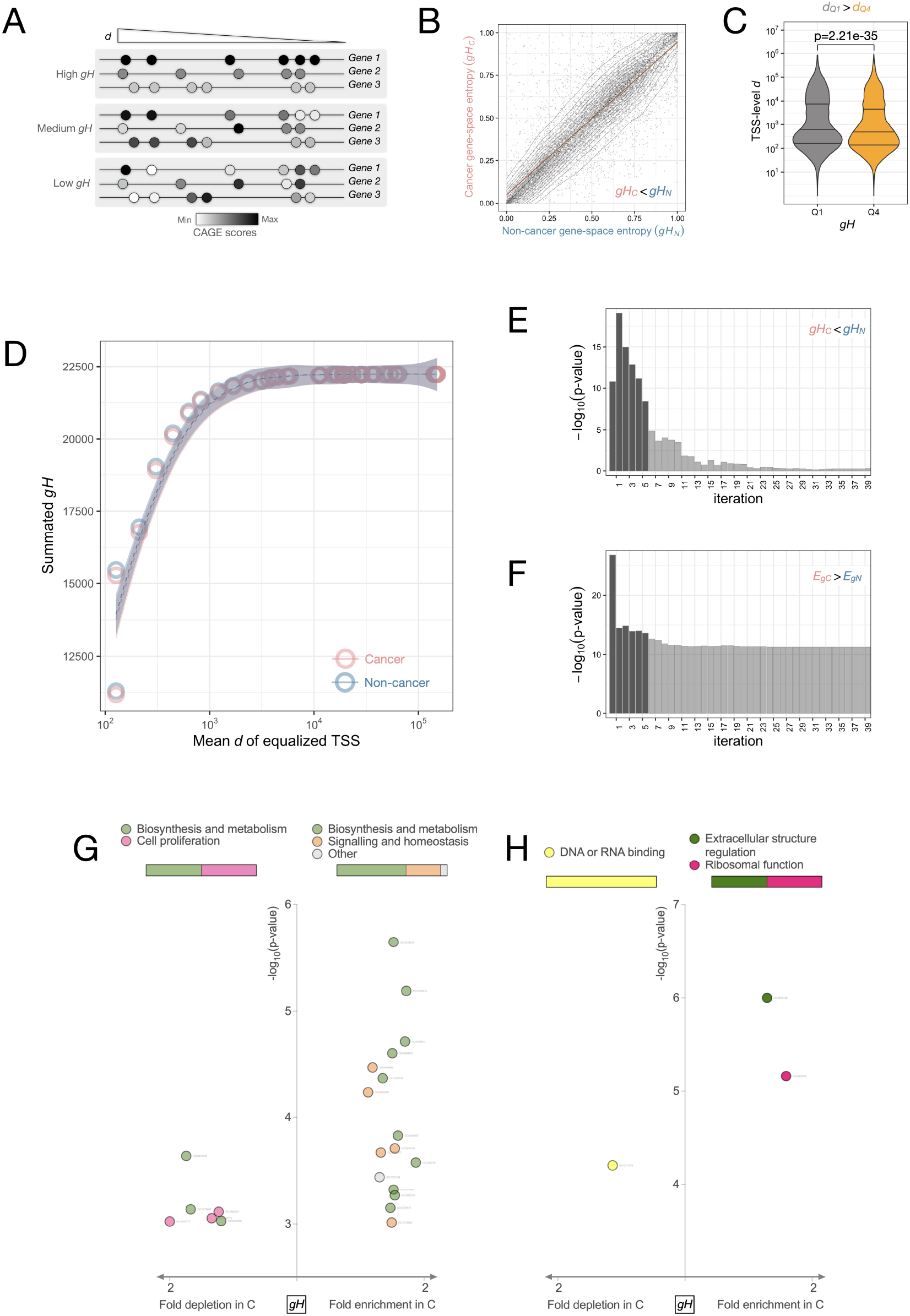
An *E* -conserving preference for proximal TSSs through gene–level stochasticity of TSS deployment (*gH*) is systemically lost in cancers but retained for genes supporting biosynthesis and cell proliferation. A: A schematic showing stochasticity of TSS deployment using a hypothetical set of three genes under conditions of high, medium or low gene–space entropy. The gradient of *d* indicated at the top shows the location of TSSs (circles in shades of grey representing TSS deployment frequencies calculated from CAGE expression scores) with the ends marked “Gene” representing the minimal *d* at the start codon. Similar deployment frequencies of all TSSs of a gene generates high gene–space entropy (topmost). Regulated and restricted deployment of some TSSs generates low gene– space entropy (bottom). B: Overall *gH* is highly similar between *C* and *N* but significantly lower in *C* (Wilcoxon signed–rank test paired by 11928 genes, p–value = 1.364e-09). C: Genes with low *gH* in cancers have high *d* TSSs and vice versa (Wilcoxon rank–sum test, p– value < 2.2e-16) indicating an association between TSS deployment and *d*. D: A forced increase in *gH* by equalization of TSS scores starting from the most proximal two TSSs and proceeding towards consecutive distal TSSs shows that *C* retains overall higher *gH* as compared to *N* (Y–axis) due to the stochastic deployment of TSSs with *d* up to 1 kb (X–axis). E and F: The loss of *gH* (E) with a concomitant increase in *E*_*g*_ (F) in *C* as compared to *N* is highly significant (Y–axes) for the first five TSS score–merging iterations (X–axes). G and H: The genes with low *gH* in *C* are significantly associated with biological processes of biosynthesis and cell proliferation (G) and molecular function of DNA/RNA–binding (H). Y– axes show p–value of Fisher exact text for significant enrichment and X–axes show the fold change of *gH* in *C*, over *N*; more details in (table S9).

Next, the two terminal quartiles of genes sorted by the (*gH*_*C*_ / *gH*_*N*_) ratios (Table S8) were compared. The subset of genes with a higher entropic TSS deployment in cancers had a significantly lower *d* and *HMd* profile of TSSs (Fig 3C Wilcoxon rank–sum test, p–value = 6.32e-18 and Fig S10 Wilcoxon rank-sum test, p-value = 9.19e-05). Supporting these results, we observed that although *gH* does not show a strong correlation with *E*_*g*_ or *d* (not shown), it exhibited a bimodal distribution such that two groups of genes could be segregated by using a *gH* threshold of 0.4 (Fig S11A). The subset of genes with *gH* > 0.4 had higher TSS *d* profiles as well (Wilcoxon rank–sum test, p–value = 6.32e-18, Fig S11B) indicating a more regulated TSS deployment. The genes qualifying into *gH* > 0.4 category independently in *C* and *N* showed no significant differences in their TSS *d* profiles (not shown). Different sets of genes fell into the *gH* <0.4 category independently in *C* and *N*. The genes belonging to this low *gH* cluster in cancers had significantly higher *d* TSSs compared to those in non–cancers (Wilcoxon rank–sum test, p–value = 0.016; Fig S11C). Although these results did not reveal if within a gene an entropic versus deterministic TSS deployment is linked to proximal or distal TSS shift, they did establish that stochastic TSS deployment is sensitive to the overall TSS *d* profile of a gene. Most importantly, the low *gH* preference in cancers for genes with high *d* TSSs raised interesting possibilities which we investigated further.

To understand how the dependence of gene–space *H* on *d* differs between *N* and *C*, we calculated the gene–space *H* and associated *E*_*g*_ for all the genes by iteratively equalizing the activities of consecutive TSSs in a sorted order of *d* (Fig 3D, see Methods, 2.12). Through this mechanism we increased the gene–space *H* by generating the maximum possible *H* for the first (*n*+ 1) number of TSSs in every *n*^th^ iteration of TSS activity equalization. The effects of <1 kb or >1 kb TSSs of any gene on its gene–space *H* and *E*_*g*_ could now be established by observing how TSS equalization affected these parameters. We performed this analysis on TSSs sorted by *d* in ascending (from the most proximal towards the most distal TSS) or descending orders (from the most distal towards the most proximal TSS). Over the iterations, *gH*_*C*_ remained less than *gH*_*N*_ and eventually equalized past 10 iterations (Fig 3E; Wilcoxon signed–rank test paired by 11928 genes per iteration, p–value < 1e-3). Similarly, *E*_*gC*_ remained higher than *E*_*gN*_ even if the change stabilized after 10 iterations (Fig 3F; Wilcoxon signed–rank test paired by genes per iteration, p–value < 1e-3). The majority of *gH* and *E*_*g*_ change was restricted to the equalization of the first five TSSs promoterome–wide (highlighted in Fig 3, E and F). As >90% of genes have five TSSs or less (Table S2 and not shown), a similar finding was obtained through equalization of the last five TSSs as well (Fig S12, A and B). These findings suggested that in a TSS landscape spanning over a wide range of *d*, stochasticity favours proximal TSS deployment in accordance with the *d* distribution of TSSs. The gene–space *H* excess in *N* over *C* is due to the preferred utilization of the first five TSSs and this confers a low *E*_*g*_ on *N* as compared to *C*.

*gH* being tightly linked to proximal TSS deployment and *E* differences between *C* and *N* raised the possibility that it would affect genes in a function–dependent manner. Similar to *vd*, we observed that Q1 of genes ranked by (*gH*_*C*_ / *gH*_*N*_) ratios were enriched in GO BP terms pertaining to cell proliferation and biosynthesis and MF terms linked to DNA/RNA binding (Fig 3, G and H; Table S9). The Q4 set was enriched in GO BP terms associated with signal transduction, similar to that observed for *vd* (Fig 3G and Table S9) and MF terms associated with extracellular structure regulation and ribosomal function. Gene–space entropy is focused on looking at inter–TSS variability within genes for a population. However, it does not account for any sample–space heterogeneity of a population. To understand the role of inter–sample heterogeneities, we compared the TSS deployment probabilities in the sample– space between *N* and *C* sample groups.

### 3.4. Sample–space entropy reveals that selective deployment of high d TSSs alongside a predominantly stochastic deployment of low d TSSs is widespread in cancers

We hypothesized that (1) that non–cancers would manifest tissue type heterogeneity through selective TSS deployment, which would lead to high inter–sample variability in TSS activities, (2) a relative–loss of tissue–specific gene expression patterns is common in cancers and that would reduce this inter–sample variability in TSS activities between samples with an effect on *E*_*t*_ . This could be tested by a cancer versus non–cancer comparison of entropies for individual TSSs across the sample–space. These sample–space entropies (*sH*) would be derived from the probabilities of deployment of each TSS across distinct samples (Table S10). An entirely stochastic deployment of TSSs would escape regulation and become similarly deployed in various samples (Fig 4A). Based on our findings so far, we argued that stochastic deployment of a TSS across samples would perturb the *E* of the system. To begin with we needed to understand the relationship between sample–space entropy and *E*_*t*_ in general independent of the *N* or *C* sample identities. Similar to the TSS equalization simulation on *gH* (see 2.3), we performed an alternative simulation focused on the sample– space with equal contributions from *N* and *C*. We iteratively equalized the scores of a TSS across all samples to steadily maximize the sample–space entropy and tested the corresponding effect on *E*. The sample–space is gene–blind and therefore all TSSs of a sample were equalized unlike *gH* where all TSSs of a gene were equalised (see Methods, 2.13). This controlled entropy increment of artificially generated promoteromes showed a strong increase in *sH* and *E*_*t*_ as compared to the naturally observed *sH* and *E*_*t*_ in *C* as well as *N*. Interestingly, there was a positive association between sample–space entropy and *E*_*t*_ (Fig S13) for all TSSs except those with *d* < 1kb. The restrained range of actual sample–space entropy and associated *E*_*t*_ as observed in *C* and *N*, (Fig S13) indicated that TSS deployment is tightly regulated in the sample–space with only a limited contribution from stochasticity. This regulated TSS deployment minimizes *E*_*t*_ and restrains entropy in the sample–space. Next, we analyzed entropies in the sample–space derived from TSS–specific CAGE scores (Table S10) in different ways.

**Fig 4.**
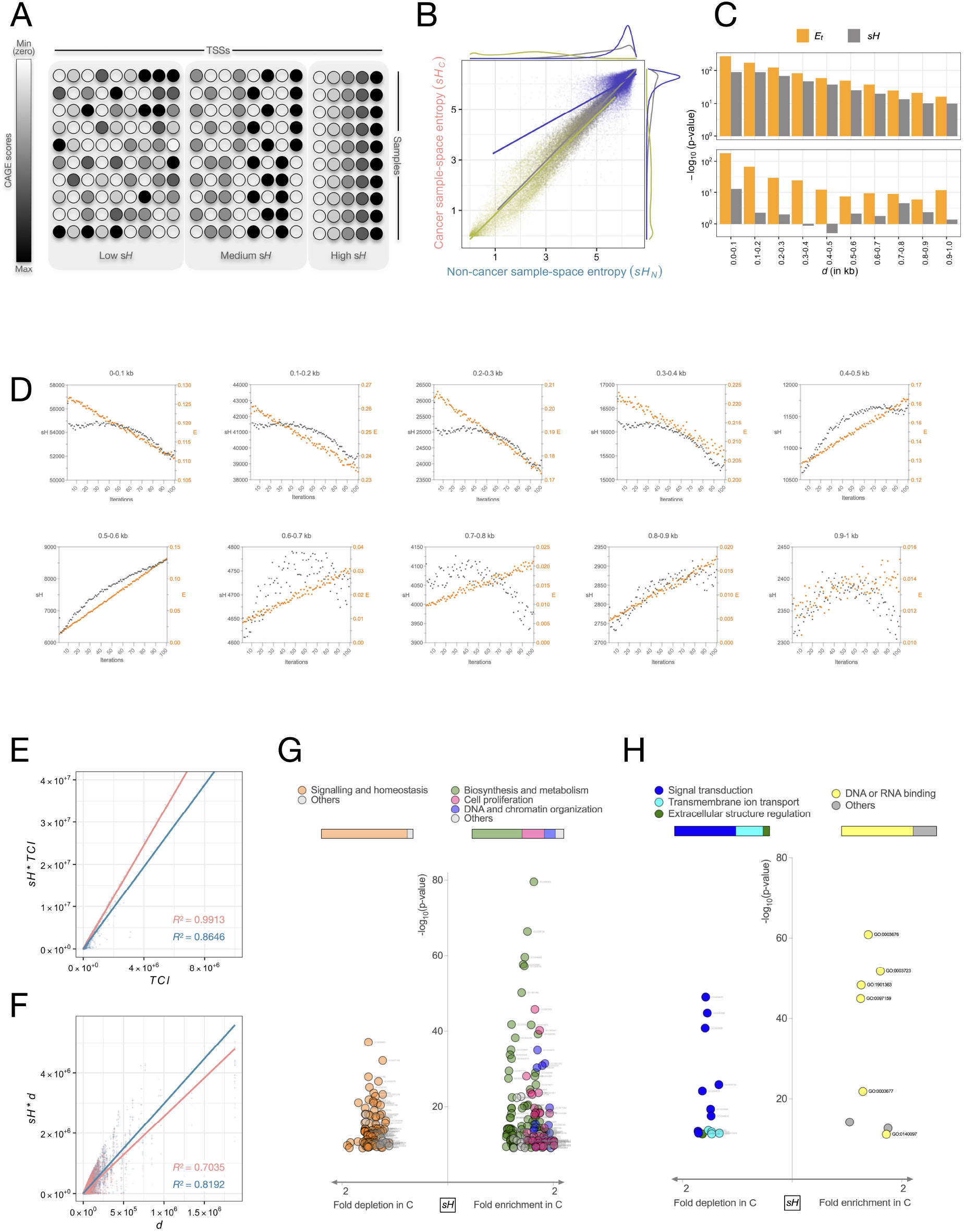
Frequent and stochastic deployment of highly proximal *d* TSSs across samples (*sH*) increases *E* in cancers through high transcriptional activity of genes involved in biosynthesis and cell proliferation. A: A schematic representation of stochasticity of TSS deployment, deduced from the entropy of CAGE scores of each TSS in the sample–space (*sH*). The inter–sample variation of deployment probabilities for a TSS (along the horizontal axis) can be high (low *sH*, left panel), moderate (medium *sH*, middle panel) or very low (high *sH*). B: *sH* is tightly correlated between *C* and *N* for all TSSs (grey dataset). When split into TSS with non–zero representation in minimum 100 samples (*frequently deployed* TSSs, blue) or *non–frequently deployed* TSSs (olive green dataset), distinct patterns of *sH*_*C*_ > *sH*_*N*_ for *frequently deployed* and *sH*_*C*_ < *sH*_*N*_ for *non–frequently deployed* TSSs emerge. C: *sH*– linked *E*_*t*_ differences, calculated as log_2_ ratios of products of *sH* and *E*_*t*_, between *C* and *N* display an association with *d*. The *frequently deployed* TSSs showed an enhanced *sH*–linked *E*_*t*_ excess in *C* over *N* (Y–axis) at proximal TSSs *d*< 0.4 kb (X–axis). D: Substitution of *C* CAGE scores with those from *N* decreases *E*_*t*_ as well as *sH* in a *d*–dependent manner for TSSs with *d*<0.4 kb (*sH* on left Y–axis (black) and *E*_*t*_ on the right Y–axis (orange), 50 iterations of CAGE score substitutions from *C* to *N* on the X–axis). These data show that the unexpected correlation between energy cost coefficient of transcription and consistency of a TSS deployment in cancers or non–cancers is tightly linked to the proximity of TSS to the start codon. E and F: *sH* -linked *TCI* is increased in *C* (E) whereas *sH*–linked *d* is decreased in *C* (F) showing that the *E*_*t*_ increase in *C* is due to transcriptional activity–excess which overrides the *d* conservation in cancers. The units of *sH* * *TCI* and *sH* * *d* and *TCI* are arbitrary. G and H: High *sH* in cancers is significantly associated with genes involved in biological processes biosynthesis and cell proliferation (G) and molecular functions DNA/RNA–binding (H), recapitulating the gene functional category enrichments observed for *vd* and *gH* (details in table S12).

First we calculated *sH* (a schematic in Fig 4A) from *N* and *C* samples for a restricted set of TSSs with non–zero CAGE scores from at least 100 *N* and 100 *C* samples. As CAGE is biased to report only TSS activity, not inactivity, and since entropy values and associated information cannot be deduced from inactive TSS, such constraints allowed sample–space entropy comparisons between *N* and *C* just from active TSSs without any detection bias. Of the 55048 TSSs reported in CAGE data, 33232 TSSs satisfied these constraints (Table S10). For these *frequently deployed* 33232 TSSs *sH* was strongly increased in cancers (Wilcoxon signed–rank test paired by 33232 TSSs, p–value < 2.2e-16; blue–label in Fig 4B). Thus, the *frequently deployed* TSSs are deployed with low inter–sample variability in cancers, whereas the inter–sample variability for these TSSs is higher in non–cancers. On the other hand, the *non–frequently deployed* 21816 TSSs (olive green label in Fig 4B) gave rise to an *N*–biased distribution of *sH* (Wilcoxon signed–rank test paired by 21816 TSSs, p–value < 2.2e-16). The overall sample–space variabilities in TSS deployments between *N* and *C* (grey–label in Fig 4B) could be classified into two different pools: the *frequently deployed* ones which have higher *sH* in *C* and the *non–frequently deployed* ones with lower overall *sH* in *C*.

To understand the cooperation between *sH* and *E*_*t*_, we compared the products or ratios of *E*_*t*_ and *sH* for each TSS between *C* and *N* (Table S11 and Fig S14A) over the entire range of *d*. The *non–frequently deployed* TSSs which have lower *sH* and *E*_*t*_ in *C*, also had lower *sH* \**E*_*t*_ in *C* (Fig S14, B and C; *C<N*, Wilcoxon signed–rank test paired by 21816 TSSs, p–value < 2.2e-16). The *frequently deployed* TSSs, however, showed a significant excess of *sH* * *E*_*t*_ in *C* owing to their higher *sH* and cooperating *E*_*t*_ (Fig S14, D and E; Wilcoxon signed–rank test paired by 33232 TSSs, p–value < 2.2e-16). The *non–frequently deployed* TSSs by definition have high variability across samples with deployment in less than 100 samples. Thus, independent of their TSS activity, their sample–space entropy would remain lower than that of the *frequently deployed* TSSs. Such low deployment frequencies across samples would lower their overall *TCI* and consequently the *E*_*t*_ . Expectedly, the *non–frequently deployed* TSSs, which have lower *sH* and *E*_*t*_ in *C*, as well as a lower *sH* * *E*_*t*_ in *C* (Fig S14A–C, Wilcoxon signed–rank test paired by 21816 TSSs, p–value < 2.2e-16). The *frequently deployed* TSSs, however, showed the opposite behaviour with *sH* and *E*_*t*_ exhibiting a *C*–bias. As a result, the *sH* * *E*_*t*_ was also higher in cancers for these TSSs (Fig S14A and D–E, Wilcoxon signed–rank test paired by 33232 TSS, p–value < 2.2e-16). It was interesting to note that the opposing behaviour of *sH* * *E*_*t*_ between the two categories of TSSs was driven by a cooperation of *sH* and *E*_*t*_ . Such concomitant differences in sample–space variability as well as *E*_*t*_ between *N* and *C* could be explained by the frequency of deployment and the distance. We thenceforth analyzed the influence of *d*–dependent TSS deployment on *sH*.

The *E*_*t*_ /*sH* profiles were analyzed for various *d* bins (Fig S15). We observed that for the *non–frequently deployed* TSSs, the nature of *sH, E* _*t*_ and *E*_*t*_ /*sH* remained *C* < *N* at all *d* values with the first 0–1kb *d* bin being the most significant (p–values tends to 0, Fig S15A). The *frequently deployed* TSSs on the other hand showed an opposite difference (*C* > *N*) for *sH, E*_*t*_ and *E* _*t*_/*sH* (Fig S15B; Wilcoxon signed–rank test paired by *d* bins, p–value < 0.05) except the last bin. However, the most significant difference between *C* and *N* for all the three parameters (*sH, E*_*t*_ and *E*_*t*_ /*sH*) were from the first *d* bin (0–1 kb) (Fig S15A). This suggested that the two categories of TSSs, *frequently deployed* and *non–frequently deployed*, have contrasting properties to balance the energy cost coefficient of transcription across the sample–space in a *d*–dependent manner. To elucidate these differences further, we performed incremental substitution of *C* sample–space with *N* sample–space and calculated *sH* as well as *E*_*t*_ . These simulations using *sH* and associated *E*_*t*_ differences between *N* and *C* for all TSSs reinforced that the lowest *d* bin of 0–1kb accounts for the majority of differences (Fig S16). A finer resolution analysis of these TSSs split into 0.1 kb bins showed that the *sH* and *E*_*t*_ differences are concentrated in a *d* of about 0.4–0.5 kb specifically for the *frequently deployed* TSSs (Fig 4D). We could conclude that in *C*, high *sH* is associated with low *d* as well as high *TCI* (deployment) of TSSs leading to an overall *E*_*t*_ excess in most TSSs. Further, by comparing *sH*–factored *TCI* and *d* (Table S11), we found that *sH* drives high *TCI* and low *d* in cancers. With increasing *sH, TCI* increased more sharply in *C* as compared to *N* (Fig 4E; Fisher’s z test for 55048 TSSs, p–value = 0.0053). To the contrary, with increasing *sH, d* decreased more sharply in *C* as compared to *N* (Fig 4F; Pearson and Filon’s z for 55048 TSSs, p–value = 0). Reinforcing these findings, the same relationship between *sH* and *TCI* or *d* are obtained when using ratios (*d* or *TCI*/*sH*) instead of the product (*d* or *TCI* * *sH*) (not shown). These findings implied that cancers in general deploy low *d* TSSs at higher *TCI* with less heterogeneity across the sample–space. By corollary, these results implied that high *d* TSSs are deployed with low *sH* and low *TCI*. This mechanism apparently leads to an *E*_*t*_ saving outcome in *C* which is limited only to TSSs with *d* less than 0.5kb. These results showed that (i) high *d* TSSs undergo more selective deployment in *C*, (ii) stochastic deployment of TSSs conserves *E*_*t*_ by retorting to low *d* in *N*, and, (iii) even in *C* the higher stochastic deployment targets a small subset of genes with low *d* profile.

Similar to *vd* and *gH, sH* also showed an association with gene functions relevant in cancers. Genes ranked by (*sH*_*C*_ /*sH* _*N*_) ratios were subjected to functional category enrichment (Table S12). The Q4 genes, representing an increased entropy in cancers, were enriched in genes involved in cell proliferation and biosynthesis (Fig 4G) as GO BP terms and DNA/RNA binding as GO MF terms (Fig 4H) and the Q1 genes primarily enriched in signalling (Fig 4, G and H). Combined with the *gH* findings, these results showed that a low *gH* and high *sH* cooperate in cancers to achieve a regulated deployment of proximal TSSs for genes involved in core cancer hallmarks. We combined the gene–space and sample–space to estimate the overall combined entropy affecting TSS deployment and its functional implications in cancer.

### 3.5 Compound entropy represents a system–level stochasticity by combining gene– level TSS selection with the sample–level heterogeneity

The effective stochasticity associated with the deployment of any TSS deducible from the CAGE scores would actually be a combination of the stochasticity in TSS deployment for any gene (estimated as gene–space entropy *gH*) and stochasticity in TSS deployment across samples (estimated as sample–space entropy *sH*). This combined stochasticity can be estimated by a compound entropy (*cH*) calculated from independent non–exclusive combinations of sample–space and gene–space probabilities for each TSS (schematic in Fig 5A). From the compound probabilities, one *cH* per gene was calculated in the gene–space (Table S13).

**Fig 5.**
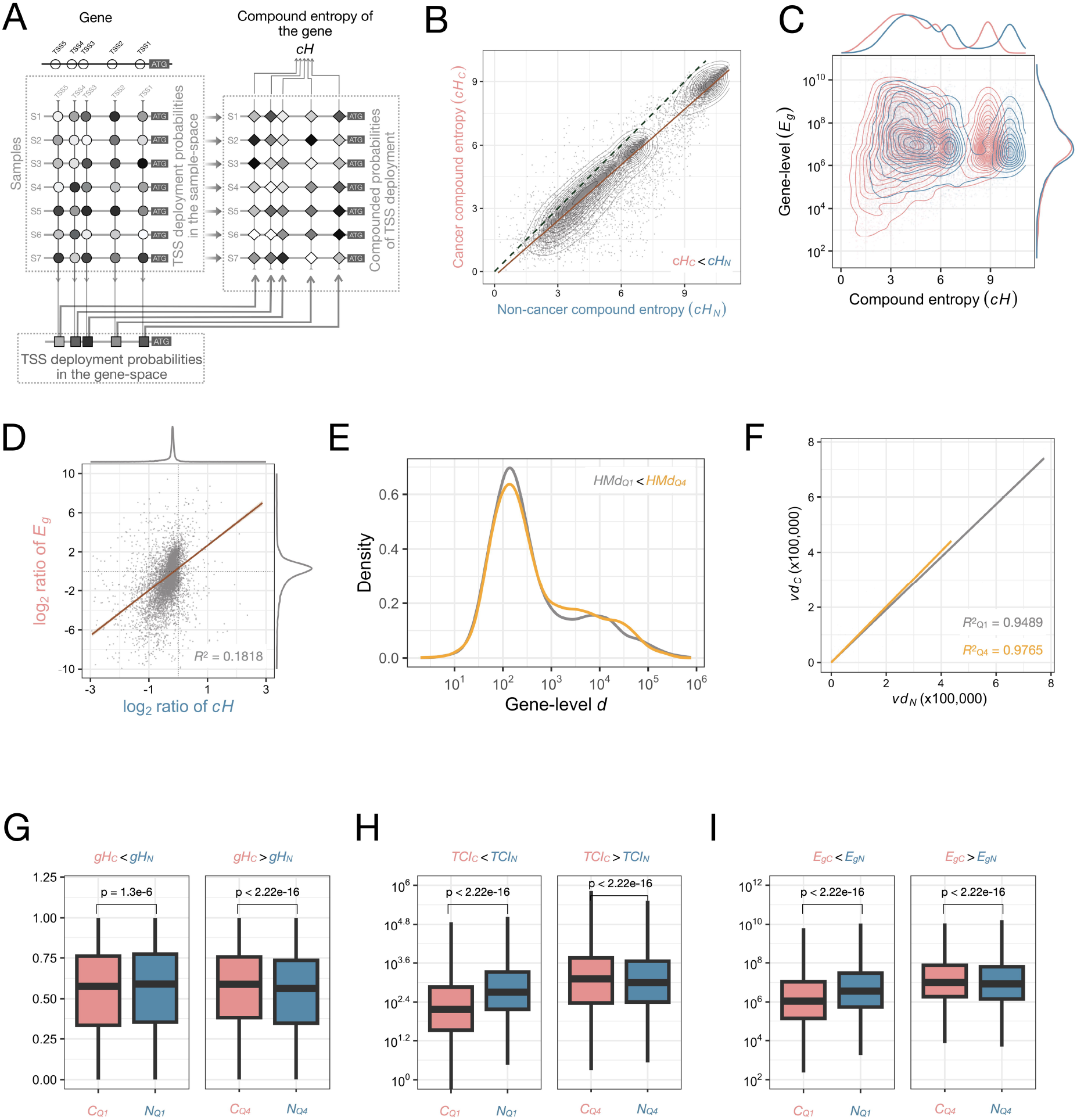
The overall stochasticity of TSS deployment (*cH*) favors a proximal transcription start and minimization of *E* at the gene–level which is disrupted in cancers. A: A schematic representation of the calculation of compound entropy (*cH*) for a hypothetical system of a 5 TSS–gene (TSSs labeled TSS1 through to TSS5) and seven samples (labeled S1 through to S7). ATG demarcates the start codon of the gene, the various TSSs are drawn at varying *d* as unfilled circles. Circles with grey shade depict sample–space probabilities, squares with grey shades depict gene–space probabilities and diamond with grey shades depict compound probabilities derived as products of the sample–space and gene–space probabilities of each TSS. The compound–space probabilities are used to calculate one *cH* value per gene. Different shades of grey represent different probabilities derived from differing CAGE scores. B: *cH* is tightly correlated between *C* and *N* but significantly lower in *C* (Wilcoxon signed–rank test paired by 11928 genes, p–value < 2.2e-16). C: *cH* shows a bidirectional relationship with *E*_*g*_ . *C* has a larger range of *E*_*g*_ than *N* at lower *cH* with a conspicuous set of genes having low *cH* and low *E*_*g*_ in *C* (*cH*<6; red arrow). At *cH*>6 the *E*_*g*_ in *C* exceeds *E*_*g*_ in *N* with a set of genes having high *cH* and high *E*_*g*_ in *C*. D: Products of *cH* and *E*_*g*_ show a tight correlation with a *C–*biased distribution showing that low *cH* and low *E*_*g*_ co–occur more frequently in *C* than *N*. E: *HMd* of genes belonging to terminal quartiles Q1 (green) and Q4 (red) of *cH* (log_2_ *cH*_*C*_ /*cH*_*N*_) show that *HMd* of Q1 is significantly lower than *HMd* Q4 (Wilcoxon rank–sum test, p–value = 1.729e-2) suggesting that genes with a high TSS *d* profile are more prone to stochastic TSS deployment in *C*. F: *vd* scatterplot between *C* and *N* with regression line for the same Q1 (*R*^*2*^=0.9489; grey–line) or Q4 (*R*^*2*^=0.9765; orange–line) genes, show a significant difference between the Spearman’s correlation coefficients (Fisher’s z test for 2982 genes, p–value < 2.2e-16) with Q4 genes set having a *C*– biased distribution. Results in B–F show that higher *cH* in *C* is associated with higher *d* which increases *E*_*g*_. At the same time the lower *cH* in *C* is associated with higher *E*_*g*_ but lower *d* suggesting an activity–driven *E*_*g*_ excess in *C* from low *d* TSSs. G–I: Consistent with the findings described above, we observed that *gH* (G), *TCI* (H) and *E*_*g*_ (I) follow a *C<N* pattern for *cH* (log_2_ *cH*_*C*_ /*cH*_*N*_) Q1 genes (Wilcoxon signed–rank test paired by 2982 genes, p–values ≤ 1.3e-6) whereas these parameters follow a *C>N* pattern for Q4 genes (Wilcoxon signed– rank test paired by 2982 genes, p–values < 2.2e-16).

Before applying compound entropy calculations for differences between *C* and *N* we sought to understand its relationship with *E*_*g*_ in simulated promoteromes (see Methods, 2.14). Using the artificial promoteromes with controlled increase in the entropy, we established that *cH* and *E*_*g*_ covary directly (Fig S17). The *cH*–*E*_*g*_ relationships from these simulations also suggested that the overall actual entropy of TSS deployments, although different between *C* and *N*, exists in a state of very low *E*_*g*_ due to gene–wise bundling TSSs. This low *E*_*g*_ state is not achievable by disregarding a regulated gene–wise TSS deployment sensitive to gene function and TSS *d* profiles.

Comparison of *cH* between *C* and *N* showed an *N*–biased distribution with *cH*_*C*_ < *cH*_*N*_ (Fig 5B; Wilcoxon signed–rank test paired by 11928 genes, p–value < 2.2e-16). *cH* and *E*_*g*_ comparisons showed different relationships between *cH* and *E*_*g*_ in *C* and *N* (Fig 5C). At the gene–level, log_2_ *c*/*N* ratios of *cH* and *E*_*g*_ showed a highly significant positive correlation even if *cH* was tightly restricted to *C*<*N* (Fig 5D; Linear regression coefficient *R*^2^ = 0.1818, p–value < 2.2e-16). *C* showed a higher *E*_*g*_ /*cH* than *N* reinforcing that for every unit increment in *cH*, the increment in *C* is higher than that in *N* (Fig S18; Wilcoxon signed–rank test paired by 11928 genes, p–value < 2.2e-16). Next, we ranked genes by *cH* ratios from *C* and *N* and compared the terminal quartiles Q1 and Q4 (Fig S19). The Q1 genes had a significantly lower *HMd* profile (Fig 5E; Wilcoxon rank–sum test, p–value = 0.017). Unlike *HMd* (the expected d representative of a gene if all TSSs were equally weighted, identical for *C* and N), the *TCI*–weighted *d* (*vd*) was higher (observed effective *d* for a gene) in cancers for the Q1 as well as Q4 genes (Fig 5F; Wilcoxon signed–rank test paired by 2982 genes each, p–value for Q1 = 2.959e-12 and p–value for Q4 = 1.035e-13). Interestingly, three parameters, not factored in calculation of *cH* (gene–level *E, TCI* and *gH*) were lower in *C* than *N* in Q1 genes as compared to the Q4 genes (Fig 5, G–I; Wilcoxon signed–rank test with p–values indicated above the bar plots). Ranked by the *C/N* ratios, the Q1 genes thus stood out as a set of genes for which low *cH* was tightly associated with low *Eg*. The Q4 genes associated with a higher compound entropy in cancers also showed an enrichment of cell proliferation and biosynthesis genes like the parameters described earlier (Fig S20 and Table S14).

Calculations of all the entropies, including *cH*, were dependent on the CAGE scores but independent of the *d* profile of the TSSs. To uncover how entropies in TSS deployment still affected the TSSs in a *d*–dependent manner, and consequently gene–level *E*, we calculated a combined parameter of *cH* and *vd*. The two parameters being inversely related to each other (Fig S21), we calculated *vd*/*cH* (Fig 6A) and used its *C/N* ratios to rank genes (Table S14).

**Fig 6.**
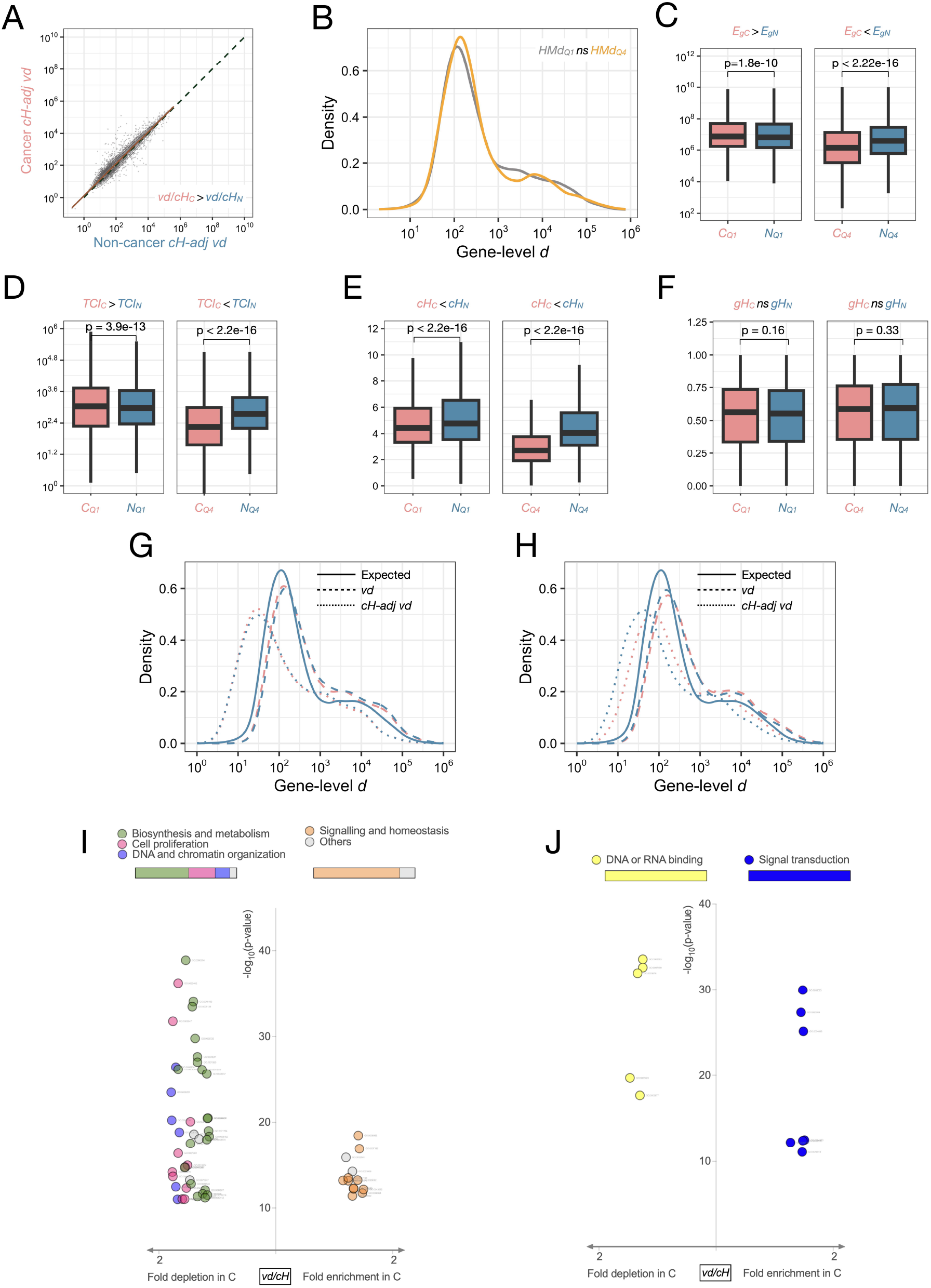
The minimizing effect of *cH* on distal TSS deployment, represented by *d*/*cH*, is selectively impaired in cancers for genes involved in cell proliferation and biosynthesis. A: A combination of *vd* and *cH* (calculated as *vd*/*cH* for each gene) is significantly lower in cancer (Wilcoxon signed–rank test paired by 11928 genes, p–value < 2.2e-16). B: Q1 and Q4 genes sorted by log_2_ ratios of *vd*/*cH* from *C* and *N* shows that, unlike for *cH* (Fig 5D), the compound entropy–factored effective *d* does not affect genes based on their TSS *d* profiles. C–E: The Q1 and Q4 genes have highly significant different *E*_*g*_, *TCI* and *cH* profiles between *C* and *N*; low *cH*–factored *vd* genes have higher *E*_*g*_ (C), *TCI* (D) and *cH* (E) in cancers and vice versa (Wilcoxon signed–rank test paired by 2982 genes in Q1 and Q4, all p–values ≤ 1.8e-10). F: *gH* shows no significant differences between *C* and *N* for either Q1 or Q4 genes. G and H: *HMd* (expected), *vd* and *vd*/*cH* comparisons between *C* and *N* for Q1 (G) and Q4 (H) genes show that both *vd* and *vd*/*cH* are decreased in *C* for Q1 and increased in *C* for Q4 genes (Wilcoxon signed–rank test paired by 2982 genes in Q1 and Q4, p–values < 2.2e-16). I and J: Functional category enrichment comparison between Q1 and Q4 genes shows that genes belonging to the biological processes cell proliferation and biosynthesis (I) and molecular function DNA or RNA binding (J) are enriched in *C* as compared to *N* (details in table S14).

Ranked by *vd*/*cH* log_2_ *C*/*N* ratios, there was no significant difference in the *HMd* profile of the Q1 and Q4 genes (Fig 6B). The *E*_*g*_ of Q1 genes were significantly greater than those of the Q4 genes in *N* as well as *C* (Fig 6C and not shown). The *E*_*g*_ excess in *C* was limited to Q1 (Fig 6C; Wilcoxon signed–rank test for Q1 and Q4 genes each paired by 2982 genes, p– values ≤ 1.8e-10). This *E*_*g*_ excess in *C* for Q1 genes was driven by a combination of *TCI* and *d* resembling the overall gene pool (Fig S22A; *r*_*c*_ < *r*_*N*_ and *r*_*c*_ < *r*_*N*_ for Q1 and Q4 respectively using Pearson and Filon’s z test, p–value = 0). The deficit of *E*_*g*_ in *C* for the Q4 genes was driven by a leaky deployment of TSSs across all *d*; For the Q4 genes even if the effective *d* was increased in *C* (*vd*_*C*_ > *vd*_*N*_ ; Wilcoxon signed–rank test paired by 2982 genes, p–value < 2.2e-16), the low *TCI* in *C* (Fig 6D) offset its effect resulting in a net lower *E*_*g*_ (Fig S22B).

For Q1 as well as Q4 genes *cH* was lower in *C* than *N* with no significant difference in *gH* (Fig 6, E and F). The Q1 genes ranked by *vd*/*cH* between *C* and *N* were expected to have higher *cH* in *C* as compared to *N*. Intriguingly however, the observed *cH* was strongly lower in *C* than *N* for the Q1 genes. The only reason why *vd*/*cH* would be lower in *C* even when *cH* is extremely low is an over–compensatory decrease in *vd* for the Q1 genes. This extremely low *vd* in *C* was not inferable from *gH* as Q1 and Q4 genes showed no significant differences in their *gH* between *C* and *N* (Fig 6F). Thus, the combination of *vd* and *cH* led us to gene sets with collaborating changes in *vd* and *cH*, two otherwise independent parameters. These compensatory changes in *vd* were evident through comparisons of *vd* or *vd*/*cH* between *C* and *N* for Q1 (Fig 6G) and Q4 (Fig 6H) genes. These results showed that while *vd* is higher than *HMd* for Q1 as well as Q4 genes, the *vd*/*cH* is lower than *HMd* (Wilcoxon signed–rank test paired by 11928 genes for *N* and *C*, p–value < 2.2e-16; not shown). *vd*/*cH* was increased over a wide range of *d* for the Q4 genes in *C* as compared to *N* (Fig 6H; Wilcoxon signed–rank test paired by 2982 genes, p–value < 2.2e-16). We next evaluated if *cH* and *vd* are functionally relevant and if their association with genes explains cancer hallmarks. *vd*/*cH* turned out to be the most efficient parameter for ranking the genes leading to functional categorization relevant to cancer hallmarks. Biosynthesis, cell proliferation and nucleic acid metabolism were highly enriched in Q1 genes ranked by *vd*/*cH* log_2_ *C*/ *N* ratios (Fig 6I and Table S14). Most emphatically, these Q1 genes were almost singularly enriched in the GO MF terms corresponding to nucleotide metabolism (Fig 6J and Table S14). The Q4 genes remained generic with an enrichment of biological processes and molecular functions relevant to signalling. Overall, *vd*/*cH* retained the functional categories (Fig 6, I and J) that were enriched or depleted in *C* as observed by the *vd* quartile genes (Fig 2, F and G). However, *vd*/*cH* was a much stronger classifier of *C* from *N* through the cancer hallmark relevant genes. *vd*/*cH*, a combination of *vd* and *cH*, is thus, a useful parameter to reveal how *TCI*–weighted *d* is exerted in a regulated and non–stochastic manner.

These functional categories enriched in genes ranked by *vd* and *vd*/*cH* differences between *C* and *N* led us to something more fundamental. A pan–GeneOntology analysis showed that the changes in *vd, vd*/*cH* and *E* were not restricted to just the genes which we identified in our analyses. Rather all the genes mapped to these GO terms showed *vd, vd*/*cH* and *E* differences between *C* and *N* recapitulating the expected directions of the differences (Fig S23 and 24, A-R) suggesting that the effect we have observed is widespread throughout the gene functional category and tightly linked to the gene function. It seemed that there is feedback from gene function on the *d*–linked TSS deployment for the entire gene sets. We argued that if this is true then we shall observe an effect on gene function feedback on TSS *d* evolution itself. A comparison of genome–wide TSS and gene–level *d* profiles showed that indeed chromatin regulatory and cell proliferation related genes have higher *d* profiles at the TSS and gene–level (Fig S24, A-D) whereas the genes related to biosynthesis and metabolism have a lower TSS *d* profile with no significant gene–level *d* differences (Fig S24, E and F). Thus, genes have primarily evolved TSSs at varying *d* under feedback from their function. The secondary evolution of TSS deployment preferences in cancers further accentuate the *d*–dependent preferences of TSS to adapt to their proliferative and biosynthetic needs. The most significantly different TSS and gene–level *d* profiles from the genomic profiles was that of the genes involved in chromatin regulation (Fig S25, A and B) suggesting that the adaptive TSS *d* switches operate at these genes capable of rewiring the cancer epigenomes for a better proliferative and biosynthetic outcome for incipient cancer cells.

## 4. Discussion

Quantitative transcriptional deregulation in cancers has been widely reported. Qualitative changes in the transcriptome are however less well studied. A major qualitative feature of transcription as a process is that although it is tightly regulated, it is fundamentally stochastic (43,44,63–66) due to the nature of TSS deployment (67–70). Since transcription regulation is significantly tied up with *cis* regulatory sequences, its stochasticity inherently puts limits on controls that can be exercised for directed mRNA synthesis (44,71–77). Studies show that TSS usage is non–adaptive (78), and has strong site– and sequence–dependent features (29,79–81), lending support to the importance of some deterministic effects of local sequence context in TSS selection. This feature of TSS usage stands in contrast to the stochasticity in determining its selection and usage. Deregulated systems such as incipient cancer cells could benefit from escaping the regulations on TSS deployment to favor their transformative evolution in a developing tumor mass.

The FANTOM CAGE data is a great resource to study transcription initiation and discover its new properties. Here we have made use of this publicly available data to understand how TSS deployment is qualitatively affected in cancers universally. The following hypotheses form the premise of our analyses: (i) That transcriptional deregulation in cancer is not just quantitative in terms of mRNA abundance but also qualitative in terms of commitment to transcribe from a particular start site, (ii) that transcription is an energy and resource– intensive process and normal cells would maintain a more energy–conserving transcriptome, (iii) that deregulated transcription in cancers would be driven by energy addiction and associated wasteful transcription, and (iv) that transcriptional resource optimization in normal cells would not only minimize unnecessary transcription initiation rather also preferentially avoid transcription from distal start sites, and finally (v) the flexibility of proximal versus distal TSS deployment would be sensitive to the gene functions with consequences on the adaptive needs of the cell.

In our analyses we have used curated subsets of CAGE samples based on commonly known sample annotation attributes of “cancer” which distinguish them from the “non–cancer”. The two sample sets include a diverse set of cancer and non–cancer samples. The heterogeneity of the sample sets ensures that our observations are not limited to a specific cancer type/cell type/tissue type; a property ascribed to cancer hallmarks. Also, the number of samples in each set gives our analyses a sound statistical confidence that the differences which survive the heterogeneities of over 1200 non–cancer and 400 cancer samples are most likely pan–cancer differences.

A more stochastic TSS deployment allows for random combinations of gene expressions, out of which, those favouring cell proliferation and survival would be selected in an evolving tumor cell population. The advantages of such a strategy for competing tumor cell subpopulations are multiple. Mutations which could alter gene expression or gene dosage to the selection advantage of cancer cells take long to manifest whereas stochastic TSS deployment for cancer phenotype–supporting gene expression allows a bypass to this dependence of cancer cells fitness on mutations. This function–relevant increased stochasticity of TSS deployment in cancer cells is likely to be affected by local epigenetic landscapes, however. Since some basal activity of TSSs, especially the distal ones, are DNA sequence encoded, any stochastic and ectopic TSS deployment will offer combinations of gene expressions without being limited by mutations and epimutations affecting them. Stochastic TSS deployment thus provides a molecular basis for multifaceted phenotypic fluidity to evolving tumor cell populations. We show that for genes belonging to functional groups biosynthesis and cell proliferation a lower stochasticity of TSS deployment favours an energy–conserving proximal TSS in accordance with the evolved TSS *d* profiles in the human genome even if the net *E*_*g*_ remains higher due to a disproportionately high *TCI*. The biosynthesis and cell proliferation gene enrichment in the relatively less stochastically deployed TSS set is accompanied by an exceedingly stochastic deployment of TSSs from high *d* TSSs; the latter having no strong functional category enrichment other than processes such as signal transduction, which do not directly confer a proliferative advantage to cancer cells. Such a high *d* TSS deployment is a gene function agnostic feature of stochastic TSS deployment that seems cancer type independent and highly energy intensive. Alongside genes involved in biosynthesis and cell proliferation, an increase in the regulated deployment of chromatin–regulatory genes with molecular functions involving DNA or RNA–binding suggests is also observed. This could support the chromatin modulation required for a more regulated proximal TSS deployment of pro–cancer phenotype genes. Metabolic anomalies which drive energy sequestering by cancer cells are universal to cancers. The absence of these metabolic anomalies in non–cancer cells automatically creates an energy resource bottleneck against stochastic wasteful transcription. The steady state gene expression profile, as represented by the transcriptomes, can hence be viewed as an outcome of a tug–of–war between high *d* TSS deployment and an energy–resource curb on it exerted through stochastic deployment of TSSs following the energy conserving TSS design of the genome. A gradual shift of this balance towards stochastic TSS deployment supported by loss of energy resource curbs is a pervasive feature of cancers. We observed it as a grand difference between heterogenous cancer or non–cancer sample sets for three independently calculated parameters B*C*I, *E*_*g*_ and *vd*, each of which represents the *d* and TSS deployment frequencies in different functional perspectives. It is no coincidence that the very genes involved in biosynthesis and core features of cancer cell fitness, growth and proliferation, exhibit a selection for energy conserving TSS deployment. Our report is the first evidence that transcriptional cost mitigation provides a feedback mechanism for genes involved in rapid biosynthetic and proliferative requirements of cancer cells.

A net higher energy cost coefficient of transcription in cancers ensues from the largely stochastic deployment of high *d* TSSs and only a selective loss of stochastic TSS deployment for cancer hallmark genes. Our results establish that this is achieved through a combination of transcriptional activity and *d* of the TSSs which can be defined as a consistent preference for more distal TSSs in cancers. A combination of preferential TSS deployment within genes and their consistent deployment in the cancer sample set, would prefer a larger effective *d*. To reconcile these observations and establish a *d*–dependence of the stochasticity in the two dimensions, an explanation for how non–cancers conduct their transcriptomes is needed. The cancer transcriptomes could then be interpreted in the light of the properties of the non– cancers transcriptomes. Our results establish that non–cancer cell and tissue type identities depend on distinct TSS deployment. The cell type–specific TSS deployment is a highly regulated process achieved through differentiation and yields low sample–space stochasticity for *frequently deployed TSSs*. The high sample–space stochasticity in cancers for the *frequently deployed TSSs*, which also have low *d* TSSs, can be understood as a disregard for the functional identity of the normal precursor cells for any cancer type as long as the cancer type independent hallmarks of biosynthesis and cell proliferation are supported by energy conserving transcription. The low *TCI* deployment of high *d* TSSs with low *sH* in cancers occurs in cancers is neither concentrated targeted to genes as per their functions nor seems to be of any adaptive advantage. It just appears to be a leaky deregulated transcription from normally silent TSSs. Thus the TSS deployment patterns in cancers display a convergent TSS deployment which is poised against the divergent TSS deployment evolved to attain and maintain cellular diversity, as observed in the non–cancers.

The TSS localization with respect to the start codons has evolved to minimize the futile transcription. This is evident in the proximal concentration of TSSs with the strongest mode at *d* <1 kb; apparently *d* is limited by the 5′ UTR length constraints. In the absence of such an evolutionary design a simple stochastic deployment of TSSs would incur a higher futile transcription and consequently a higher energy and resource cost of transcriptomes. The gene–space entropies are higher in non–cancer cells indicating that the energy saving mechanism due to stochastic deployment of proximally concentrated TSSs is sufficient to overcome the regulated tissue–type deployment of >1 kb TSSs. This higher stochasticity in the gene–space as an energy–saving mechanism for non–cancer transcriptomes significantly limits the *E*. Cancers maintain a higher gene–sapace entropy of TSS deployment which is inherently more expensive.

The CAGE data using which we have worked out the entropies give information only about the end process of mRNA capping as a marker of the successful TSS deployment; the former could not materialize without the latter. Events like aborted transcription, RNA Polymerase falling off are likely to entail a higher futile transcription cost if the pre–abortion transcribed *d* is large. The possibility that such initiated TSSs not leading to capping and CAGE capture would only add to the *E* from the more distal TSSs and would further accentuate the *E* excess due to high *d* TSS deployments in cancers. The regulatory layers which can impede ectopically started transcription can be contained in the intervening region between the TSSs and the start codon. For instance, the TSS and the start codon could be segregated into distinct TADs or sub–TADs. The fate of transcripts generated from such long–distance TSSs remains unrepresented in the CAGE data. Our observations are based on the assumption that all capped mRNA reported in the CAGE data are translation–competent. If this were to be not true we would not have observed a *d*–dependence wherein *d* is the distance between the TSSs and respective start codons. Similarly, our assumptions are supported by our gene–level analyses where TSSs are pooled by the ORF they encode and we observe a *d*–dependent TSS deployment. There are additional complexities of transcript processing which could be exceptional to our observations. These include events, although infrequent, of template– switching which would simply bypass the long regions between TSSs and start codons. RNA–seq data can be queried to assess transcriptional activities in the intervening regions between TSSs and start codons. However, an unequivocal proof requires a contiguous readout from the TSS to the start codon. For TSSs located tens of kilobases away from the start codons, such information is unavailable. Even if the CAGE data do not allow us to address these possibilities, our observations are justifiable from the energy conservation feature of natural systems. Long distance transcription entails a higher resource usage and non–cancer cells exercise a low stochasticity mechanism to deploy them. Deregulated TSS deployment in cancers is most prominent at long distance TSSs. This energy and resource intensive stochastic deployment of TSSs in cancers is likely to derive support from addiction of cancer cells to glucose, glutamine and an overt preference for biosynthetic pathways, including nucleotide biosynthesis.

The futile transcription distance affected by the choice of TSS is not functionally irrelevant. The stochastic deployment of distal TSSs in cancers selectively spares genes which code for some of the most fundamental features of cancer cells. These include cell proliferation, DNA and chromatin organisation, and RNA processing (including transcription and splicing). This discriminative transcription of cancer phenotype–supporting genes preferentially from proximal TSSs cannot be achieved through a stochastic deployment. On the contrary, these genes present a case for a selective and deterministic disallowance for stochastic distal TSS deployment. Apparently, two different spheres of stochasticity affect TSS deployments in a representative cancer cell; a gene function–agnostic very high stochasticity for distal TSSs and a gene function–sensitive low stochasticity TSS deployment for proximal TSSs. Such a duality of stochastic TSS deployment would require spatiotemporal segregation of TSSs in the nuclear space and co–localizing TSSs would share similar transcriptional resources with similar stochastic or regulated deployment outcomes. A *d*–dependent TSS colocalization is not established in cancers. However, if similar transcriptional activity outcomes depend on TSS colocalization, then it follows from our analyses that non–cancers and cancers have fundamentally different TSS colocalization patterns. The non–cancers would maintain a chromatin topology and hence TSS colocalization necessary for the functional outcome of the specific cell type. The cancers on the other hand abandon the cell type–specific regulated TSS colocalization patterns and render the chromatin topology stochastic. In evolving tumor cell masses, the cells which maintain and enrich chromatin topologies compatible with regulated proximal TSS deployments for cell proliferation and biosynthesis genes save on the energy cost coefficients of transcription and get selected in an energy–constrained tumor environment. Such a selection of cancer phenotype–supporting TSS colocalization would follow as a secondary epigenetic outcome due to a primary change in the TSS deployment landscapes of chromatin regulatory genes. This argument derives support the observations that the GO BP term chromatin regulation (highly enriched in the cancer–specific changes in *vd, E*_*g*_, *gH, sH, cH* and *vd*/*cH*) co–occur with DNA/RNA–binding as the only GO MF terms.

A combined interpretation of these findings suggest that there is a feedback between the transcriptional resource cost for a gene and the fitness advantage for a cancer cell accorded by the function of that gene upon its expression. This feedback selects against a chancy and high cost transcription from distal TSSs for those genes which are fundamental drivers of cancer cell fitness and selection in an evolving tumor. Additional support for this argument comes from the observation that the functional categories of genes for which there is a stochastic distal TSS preference in cancers do not support cancer phenotype unlike those with a proximal TSS preference. The parameter *vd* allows us a gene–specific estimation of *d* preference. Its difference between non–cancer and cancer samples represents the increase in the effective *d* for each gene. That most cancer samples cluster differently from most non– cancer samples just based on genome–wide *vd* profiles show that *vd* is a useful parameter. It combines quantitative differences in gene expression with qualitative features of TSS choice and associated resource costs and differentiates cancers from non–cancer samples. The *vd* differences between non–cancer and cancer also allow us to understand how TSS deployment is interlaced with its resource cost and gene function. The parameter *vd*/*cH* combines two critical components: the effective *d* for a gene regardless of the stochasticity or regulation underlying its manifestation, and compound entropy which represents the entire measurable stochasticity of TSS deployment for each gene across a sample set. *d* is a fixed parameter whereas *TCI* reduces the sample–space to 1. As such, by themselves, the parameters *d* and *TCI* represent components of the overall energy cost coefficients (*vd* and *E*_*g*_) without taking into account the stochasticity associated with TSS deployments. *d* being identical to all samples for any given genome can only be deployed differently in different samples with different *E* outcomes. *cH* combined the stochasticity in two independent dimensions (samples and genes) using which the stochasticity underlying any *vd* could be estimated. These results established that *cH* is a gene function–relevant estimator of the stochasticity with which TSSs at different *d* are deployed relative to each other in heterogeneous pools of cancers or non– cancers. By themselves neither of these two parameters (*vd* or *cH*) are expected to be concentrated at genes with specific cancer–relevant functions. It follows then that a combination of *vd* and *cH* is also unlikely to be a gene–function–relevant parameter. *vd*/*cH* combines these two parameters in a way that it allows cancer versus non–cancer distinction. *vd*/*cH* differences between *C* and *N* shows that effective *d* of transcription is achieved through a limited stochastic TSS deployment that is calibrated by the functional outcome of the gene in question. Of all the parameters tested (*gH, sH, cH, vd* and *vd*/*cH*) the most significant enrichment of biosynthesis and cell proliferation genes was obtained by this compound parameter of *vd*/*cH*. Thus, at these cancer hallmark phenotype driving genes the effective *d* minimization and overall stochasticity of TSS deployment are not independent. These two parameters cooperate such that high *cH* drives low *vd* for cancer phenotype driving genes at a higher energy cost coefficient. As a corollary, the specific deployment of high *d* TSSs spares cancer hallmark driving genes at a high *E* with apparently a much attenuated functional feedback on TSS deployment.

Thus, the energetics of transcription contributes to cancer–supporting gene expression patterns through a feedback between transcribed futile distance and the corresponding gene functions. The TSS deployment in non–cancer cells follows an energy–conserving design and cancer cells universally subvert it to the extent it benefits their selection in the process of tumor evolution.

## 5. Author contribution

ALS: Conceptualization, data curation, formal analysis, investigation, methodology, software, validation, funding acquisition, visualization, writing – original draft, review and editing.

PK: Data curation, resources and validation.

IM: Data curation, resources and validation.

US: Conceptualization, formal analysis, investigation, project administration, supervision, validation, funding acquisition, visualization, writing – original draft, review and editing.

## 6. Acknowledgement

The authors acknowledge ParamAnanta Supercomputing facility at IIT Gandhinagar and inputs from Ruthwick Meduri.

## 9. Data availability

This manuscript utilises only publicly available data as mentioned in the methods section. There is no other data availability statement applicable to this study.

## Appendix

Custom codes used in this work are available over GitHub (Available upon request).

## Supplementary Figure Legends

**Fig S1. Incremental *d* profiles calculated gene–wise as ratio of *d* difference between two TSSs and the *d* of the proximal TSS in the pair shows that there is a selection for proximal concentration of TSSs**. The frequency distribution shows that the majority of TSSs are at a *d* which is up to three orders of magnitude (region between the two red dotted lines) lower than the *d* expected in the absence of a selection pressure favoring reduction of *d*.

**Fig S2. Density distribution of 5^′^ UTR lengths (grey) obtained from GENCODE and TSS *d*–profiles (red) calculated *de novo***. Expectedly, the TSS *d* distribution is larger than or equal to the 5′ UTR lengths.

**Fig S3. A. TSS representation profiles in cancer (pink) and non–cancer (blue) sample sets highlighting extremes of TSS detection patterns at the two ends of the X–axis**. The sample–type restricted TSSs are present in a smaller proportion of the sample set (the left end of the X–axis) whereas the commonly detected TSSs in cancers as well as non–cancers are present in a larger proportion of the sample set (the right end of the X–axis). The two datasets are plotted without any pairing along the X–axis. B. A correlation between the sample counts (non–cancer on the X–axis and cancer on the Y–axis) in which a TSS occurs shows that the majority of TSSs are detected in the same number of samples. The X and Y axes values are multipliers of the counts depicted in the color bar to the right. C. TSSs that are detected in a sample–type exclusive manner (cancer exclusives in pink and non–cancer exclusives in blue) have a significantly different *d*. The *d* of cancer exclusive TSSs (in <5% of non–cancers) are significantly larger than those of the non–cancer exclusive TSSs (in <5% of cancers); (Wilcoxon rank–sum test, p–value = 8.75e-4).

**Fig S4. *TCI* in cancers (*TCI*_*C*_, pink) is significantly higher than *TCI* in non–cancers (*TCI*_*N*_, blue)**. Wilcoxon signed–rank test paired by 55048 TSSs, p-value ≤ 7.846e-3.

**Fig S5. Distribution of TSSs with maximum *TCI* (*TCI*_*max*_) (A) or minimum *TCI* (*TCI*_*max*_) (B) shows *d*–dependent differences between *C* (pink) and *N* (blue)**. As *d* exceeds 1kb *TCI*_*max*_ and *TCI*_*min*_ distributions attain a larger range (between 10^1^ and 10^5^ on the Y–axes in both the plots) with diminishing differences. Remarkably, even if overall*TCI*_*C*_ >*TCI*_*N*_, the *TCI*_*max*,_ was not significantly higher than *TCI*_*max,N*_ (A) even if *TCI*_*min,C*_ was significantly lower than *TCI*_*min,N*_ (Wilcoxon signed–rank test paired by 16458 *d*s, p–value = 2.85e-82. Pairings of TSSs are only by *d*, not by gene). For *TCI*_*max*_ and *TCI*_*min*_ both a low level of “leaky” transcriptional activity was observed for *C* (below the dashed horizontal line a Y–axes value of 10^1^).

**Fig S6. TSS–level *E* (*E*_*t*_; *E*_*tN*_ for non–cancers and *E*_*t*_ for cancers) and gene–level *E* (*E*_*g*_; *E*_*gN*_ for non–cancers and *E*_*gC*_ for cancers) comparisons between cancers and non– cancers reveals that *E* minimization is a gene function–dependent feature which is compromised in cancers**. *E*_*tC*_ < *E*_*tN*_ without gene–level bundling of TSSs (A, Wilcoxon signed–rank test paired by 55048 TSSs, p–value = 5.9e-17). This reverses to *E*_*g*_ > *E*_*g*_ after gene–level bundling of TSSs (B, Wilcoxon signed–rank test paired by 15861 genes, p-value = 1.65e-27).

**Fig S7. A *d*–independent relative preference for distal TSS deployment for a gene, calculated as *TCI* ratios of a TSS and its preceding proximal TSS (referred to as distal/proximal ratios), show a bias for distal TSS deployment in cancers**. A: A frequency distribution of the log_2_ *C*/*N* of the distal/proximal ratios shows a clear *C*–heavy skew. B: For each pair of consecutive TSSs involved in the calculation of the ratios presented in A, a Wilcoxon signed–rank test comparing *C>N* was performed and the p–values are presented as -log_10_ on the Y–axis. The X–axis depicts the number of TSSs per gene. Total number of genes analyzed = 11928. All genes with single TSSs were excluded. No ratios were calculated for the first most proximal TSS for all the genes.

**Fig S8. *BCI* in cancers (*BCI*_*C*_, pink) is significantly higher than *BCI* in non–cancers (*BCI*_*N*_, blue);** Wilcoxon signed–rank test paired by 55048 TSSs, p–value = 3.67e-37.

**Fig S9. A: *vd* in cancers (*vd*_*C*_, pink) is significantly higher than *vd* in non-cancers (*vd*_*N*_, blue);** Wilcoxon signed–rank test paired by 15861 genes, p–value = 2.21e-35. B: An M–A plot of the terminal quartiles of genes sorted by *vd* ratios calculated as *vd*_*C*_ /*vd*_*N*_ (grey dataset is Q1 with a reduction of *vd* in cancers and yellow dataset is Q4 with an increase of *vd* in cancers). All single TSS genes were excluded. These Q1 and Q4 genes were used for functional category enrichment analyses.

**Fig S10. Comparison of *HMd* of the terminal quartiles of genes sorted by *gH* ratios calculated as *gH*_*C*_/*gH*_*N*_ (grey dataset is Q1 with a reduction of *gH* in cancers and yellow dataset is Q4 with an increase of *gH* in cancers) shows that TSSs with a less stochastic deployment in cancers (lower gene–space entropy of TSS deployment; Q1) have a significantly higher gene–level** *d*; Wilcoxon signed–rank test paired by 11928 genes, p–value = 9.19e-15.

**Fig S11**. A: A subjective classification of genes based on the *gH* threshold of 0.4 (dashed line) classifies genes into two groups (*gH*>0.4 and *gH*<0.4). B: TSSs belonging to the genes with *gH*<0.4 also have lower *d* as compared to the TSSs of genes with *gH*>0.4 (Wilcoxon rank–sum test, p–value for *C* = 7.99e-25, and p–value for *N* = 1.73e-42). Genes classify into the *gH*>0.4 or *gH*<0.4 independently in *C* and *N*. C: For the *gH*<0.4 genes (different sets of genes in *C* and *N*) the TSS *d* profiles are significantly different (*d*_*C*_ > *d*_*N*_; Wilcoxon rank– sum test, p–value = 0.016).

**Fig S12**. A: Induced *gH* maximization by merging of TSS scores from the most distal towards the most proximal TSS shows that *gH*_*C*_ remains lower than *gH*_*N*_ when merging the terminal 10 TSSs (six iterations highlighted for comparison with the proximal to distal merging data in Fig 3E) with high significance. B: The same iterations as described for A above were used to calculate *E*_*g*_ and *E*_*gN*_ . *E*_*gC*_ remained higher than *E*_*gN*_ over most iterations even if the maximal *E*_*g*_ excess in *C* is linked only to the difference in the deployment of the two most distal TSSs. Y–axes show -log_10_ p–value and X–axes show TSS score–merging iterations (p–values reported from Wilcoxon signed–rank tests paired by 11928 genes in every iteration).

**Fig S13. Controlled *sH* increment of artificially generated promoteromes (see Methods, 2.13) shows that a change in *sH* affects *E*_*t*_ in a *d*–dependent manner:** A. TSSs with *d*<1kb do not show an increase in *E*_*t*_ as *sH* is forced to increase. B–E: TSSs with d 1– 5kb (B), 5–10 kb (C), 10–50 kb (D) and >50 kb (E) all show an *E*_*t*_ increase as *sH* is forced to increase. The pink and blue dotted lines show *E*_*t*_ interpolations using actual *sH* from *C* (pink) or *N* (blue). Notably, only for TSSs with *d*<1kb the *sH* and *E*_*t*_ remain in the range showing that the *sH* for the *d*<1kb TSSs is high enough to resist further forced increment.

**Fig S14**. A: *sH* and *E*_*t*_ cooperate differently in cancers and non–cancers. *Non–frequently deployed* TSSs have *sH* * *E*_*t*_ significantly lower in cancers whereas the same parameter is significantly higher in cancers for *frequently deployed* TSSs (Wilcoxon signed–rank test paired by 21816 and 33232 TSSs respectively, p–values < 2.2e-16). B and C (*non–frequently deployed* TSSs): A density distribution of log_2_ *c*/*N* ratios of *sH* * *E*_*t*_ from shows a decrease in *C* (X–axis values <0, blue AUC). A scatter plot of log_2_ *C*/*N* ratios of *sH* (X–axis) and *E*_*t*_ (Y–axis) shows that the majority of low *sH* and *E*_*t*_ (bottom left quadrant) are observed in *C*. D and E (*frequently deployed* TSSs): The distribution of log_2_ *c*/*N* of *sH* * *E*_*t*_ ratios (D), and *sH* versus *E*_*t*_ scatter plot (E) for the *frequently deployed* TSSs shows that the *frequently deployed* TSSs have a pattern of cooperation between *sH* and *E*_*t*_ which is opposite to that observed for *non–frequently deployed* TSSs.

**Fig S15. *Frequently* or *non–frequently deployed* TSSs exhibit different *d*–dependent patterns of *E*_*t*_ as a function of *sH* (calculated as *E*_*t*_/*sH*) in *C* and *N***. A: *Non–frequently deployed* TSSs (CAGE scores available from <100 samples) show a decline in the significance of *E* _*t*_/*sH* difference (*C<N*) with increasing *d* till 10kb but increase at TSSs with *d* >10kb albeit at all *d* values the difference remains highly significant (Wilcoxon signed– rank test paired by TSSs in the *d*-bins, p–value < 0.05). B: *Frequently deployed* TSSs (CAGE scores available from ≥100 samples) show a declining significance of difference (*C>N*), albeit significant, till ≤50kb. Extremely distal TSSs (>50kb) show no significant increase in *C* as compared to *N* (Wilcoxon signed–rank test paired by TSSs in the *d*-bins, p–value < 0.05 except for the last bin). Parameters are color-coded: *E*_*t*_ (orange), *sH* (grey). *E*_*t*_ /*sH* (green). X–axes show the discrete TSS *d*–bins and Y–axes show -log_10_ P of the test of significance. p–value of 0.05 indicated by a horizontal dashed line.

**Fig S16. *sH* differences between *C* and *N* are revealed by iterative and incremental substitution of CAGE scores of *C* with those from *N* (see Methods, 2.14)**. The different scatter plots represent data split by *d*–bins; 0–1 kb (A), 1–5 kb (B), 5–10 kb (C), 10–50 kb (D) and >50 kb (E). The entire substitution spanning 50 independently performed iterations on randomized inputs is plotted from 0% *N* and 100% *C* (left ends of the X–axes) to 100% *N* and 0% *C* (right ends of the X–axes). Left Y–axes show *sH* and right Y–axes show corresponding *E*_*t*_. A complete disconnect between *E*_*t*_ (increasing with iterations) and *sH* (decreasing upon restoration of *C* to *N*) is visible only for the TSSs with *d*<1kb (A) with other bins showing varying degrees of direct correlation between *E*_*t*_ and *sH* (B–E). The findings (panel A compared with the rest) demonstrate that higher sample–space entropy in *C* conserves *E*_*t*_ only for the most proximal TSSs.

**Fig S17. Artificial promoterome reconstructed using actual CAGE data with equal contributions from *C* and *N* randomly shows that *E*_*g*_ increases rapidly with externally forced increase in *cH*.** The actual *cH* and *E*_*g*_ observed in CAGE samples range similar in *cH* but much lower in *E*_*g*_ . The gene identity–blind increase in TSS scores increases *E*_*g*_ tremendously. *cH* and associated low *E*_*g*_ observed in actual samples is thus a relationship governed by gene–level bundling of TSSs implying a gene function feedback governing *cH* and related *E*_*g*_ .

**Fig S18. *E*_*g*_ as a function of *cH*, calculated as *E*_g_/*cH* comparison between *C* and *N* shows a high similarity between *C* and *N* with a heavy *C*–biased distribution**. For every unit increase in *cH, E*_*g*_ increases in *C* more than that in *N*. The correlation scatter plot shows that this trend holds true for most genes.

**Fig S19. *cH* is overall lower in *C* than *N***. The Q1 as well as Q4 (terminal quartiles) of log_2_ *cH*_*C*_/*cH*_*N*_ have *cH* significantly lower in cancers but the magnitude of difference is stronger in Q1 genes.

**Fig S20. *cH* difference between *C* and *N* is linked to gene functions**. *cH*, although overall lower in *C* compared to *N*, shows a strong enrichment in *C* for genes involved in the same cancer hallmark functions as observed earlier for *vd, gH* and *sH*, including biological processes biosynthesis, chromatin organization and cell proliferation (A), and molecular functions DNA or RNA–binding are noteworthy (B). Y–axes show p–value of Fisher exact test for significant enrichment using GOrilla Gene Ontology term enrichment and X–axes show the fold change of *cH* in *C*, over *N*. Specific Gene Ontology terms (marked in light grey fonts for every data point and presented in table S14) are merged into color–coded assigned terms presented as legends on the top of the scatter plots.

**Fig S21. Overall distribution of *cH* (X–axis) and *vd* (Y–axis) for 11928 genes in *C* (pink) and *N* (blue)**. The *vd* differences between *C* and *N* arise at the lowest and the highest extremes of *cH. vd* in *C* is lower than that in *N* at *cH* values lower than 5. It shows that the enhanced preference for extremely low *d* TSSs is less stochastic, and by corollary more deterministic, in nature. The scatter does not present the data as paired between *C* and *N* by gene identities. The pairing is only by the plotted parameters *cH* and *vd*.

**Fig S22. *vd* as a function of *cH*, calculated as *vd*/*cH* varies between *C* and *N* with distinct patterns of *d*–dependent TSS deployments**. Q1 (A) or Q4 (B) genes obtained from genes ranked by log_2_ *C*/*N* ratios of *vd*/*cH* show that leaky low level TSS deployment in *C* (pink) is less frequent for Q1 genes (A) than the Q4 genes (B). This enhanced leaky TSS deployment in cancers is linked with high genes having *vd*/*cH* ratios in cancers. The insets show Spearman coefficients for the *d*–*TCI* correlations as per the sample color code. In A as well as B *r*_*N*_ and *r*_*C*_ are significantly different (Pearson and Filon’s z test, *r*_*C*_ < *r*_*N*_ for Q1 genes and *r*_*C*_ > *r*_*N*_ for Q4 genes with p–values < 2.2e-16).

**Fig S23. Differences between *C* and *N* for the parameters (identified through 1og_2_ *vd*_*C*_/ *d*_*N*_) *TCI, E*_*g*_, *BCI, vd, cH* and *vd*/*cH* are properties related to gene function and not specific gene identities or locations**. Significant *C* versus *N* differences exist for all the genes contained in the GO terms differentially enriched in cancers (as shown in Fig 2, F and G). DNA and chromatin organization (A–F), Biosynthesis and metabolism (G–L), Cell proliferation (M–R), *TCI* (A, G and M), *E*_*g*_ (B, H and N), B*C*I (C, I and O), *vd* (D, J and P), *cH* (E, K and Q), *vd*/*cH* (F, L and R). Actual GO terms for which the genes were fetched for the analyses from GeneOntology database are mentioned in table S7. Samples are indicated along the X–axes and the Y–axes depict the parameters described above and indicated by the axes. Values on the Y–axes are median ± IQR and 95% CI. Differences between *C* and *N* are indicated at the top of each plot with an accompanying p–value (Wilcoxon signed–rank test paired by 947 genes (A–F), 5031 genes (G–L) and, 781 genes (M–R), p–values ≤ 4.64e-11).

**Fig S24. Genes belonging to the differentially enriched functional categories between *C* and *N* (identified through log_2_ *vd*_*C*_/ *d*_*N*_) have significantly different *d* profiles as compared to the rest of the genes** (A–F; Wilcoxon rank–sum test, p–value ≤ 1.72e-6). Genes belonging to the functional categories are indicated along the X–axes by the assigned terms and the term Control gene–set (referring to all genes except the genes belonging to the functional category plotted to the right in the same graph. Y–axes represent the indicated parameters gene–level *d* (*HMd*) or TSS–level *d*. Values on the Y–axes are median ± IQR and 95% CI. DNA and chromatin organization (A and B), Cell proliferation (C and D) and Biosynthesis and metabolism (E and F), *HMd* (A, C and E), *d* (B, D and F). Actual GO terms for which the genes were fetched for the analyses from GeneOntology database are mentioned in table S7.

**Fig S25. Differences between *C* and *N* for the parameters (identified through log_2_ *C*/*N* of *vd*/*cH* ratios) *TCI, E*_*g*_, *BCI, vd, cH* and *d*/*cH* are properties related to gene function and not specific gene identities or locations**. Significant *C* versus *N* differences exist for all the genes contained in the GO terms differentially enriched in cancers (as shown in figure 2, F and G). DNA and chromatin organization (A–F), Biosynthesis and metabolism (G–L), Cell proliferation (M–R), *TCI* (A, G and M), *E*_*g*_ (B, H and N), B*C*I (C, I and O), *vd* (D, J and P), *cH* (E, K and Q), *vd*/*cH* (F, L and R). Actual GO terms for which the genes were fetched for the analyses from GeneOntology database are mentioned in table S14. Samples are indicated along the X–axes and the Y–axes depict the parameters described above and indicated by the axes. Values on the Y–axes are median ± IQR and 95% CI. Differences between *C* and *N* are indicated at the top of each plot with an accompanying p– value (Wilcoxon signed–rank test paired by 1158 genes (A–F), 6506 genes (G–L) and, 1237 genes (M–R), p–values ≤ 9.46e-14).

**Fig S26. Genes belonging to the differentially enriched functional categories between *C* and *N* (identified through log_2_ *C*/*N* of *d*/*cH* ratios) have significantly different TSS profiles as compared to the rest of the genes** (A–F; Wilcoxon rank–sum test, p–value ≤ 1.03e-4). Genes belonging to the functional categories are indicated along the X–axes by the assigned terms and the term Control gene–set (referring to all genes except the genes belonging to the functional category plotted to the right in the same graph. Y–axes represent the indicated parameters gene–level *d* (*HMd*) or TSS–level *d*. Values on the Y–axes are median ± IQR and 95% CI. DNA and chromatin organization (A and B), Cell proliferation (C and D) and Biosynthesis and metabolism (E and F), *HMd* (A, C and E), *d* (B, D and F). Actual GO terms for which the genes were fetched for the analyses from GeneOntology database are mentioned in table S14. Note that the genes belonging to the assigned term Biosynthesis and metabolism did not have significantly different *vd* compared to the rest of the genes when identified through log_2_ *vd* _*C*_/*vd*_*N*_ (please refer to Fig S24E). However, *vd* is significantly different between genes mapping to Biosynthesis and metabolism when *vd* is factored by *cH* and the genes are ranked through log_2_ *C*/*N* of *vd*/*cH* ratios (compare Fig S26E with Fig S24E).

## Supplementary Table Legends

**Table S1. The table contains the GRCh38–hg38 genomic coordinates of the region between the first base of a TSS of a gene (CAGE–derived) and the first base of the most upstream start codon for that gene (GENCODE–derived):** The length of this region is marked as *d* for that corresponding TSS in all subsequent tables denoting the distance (in bp) of the TSS from its start codon. TSSs with the same ID and overlapping coordinates were merged. TSS IDs follow the nomenclature as available in FANTOM CAGE data.

**Table S2. The table mentions the TSS–level distance parameters *d*, incremental *d* and fractional *d*:** *d* – the distance of the TSS from its corresponding start codon (as described in table S1, incremental *d* – the ratio of *d* increment for a (distal) TSS from its penultimate (proximal) TSS and *d* of the penultimate TSS, and, fractional *d* – the ratio between the *d* of a TSS and the highest *d* for a TSS for the same gene.

**Table S3. The table contains the gene–level distance parameters *HMd, vd* in *N* and *C* and the log_2_ /*N* of *vd* for 15861 genes that have curated TSSs common in *N* as well as *C***. *HMd* (harmonic mean *d* of all TSSs for a gene), *vd*_*N*_ and *vd*_*C*_ (*TCI*–weighted mean *d* of a gene in non–cancers (*vd*_*N*_ ) and (*vd*_*C*_ ) cancers).

**Table S4**. The table contains CAGE samples from FANTOM5 with sample identities as mentioned in FANTOM, curated sample number appendix with a binary *N* (non–cancer) or *C* (cancer) sample ID.

**Table S5. The table is the CAGE raw score matrix for all the samples:** The first column header is TSS name (in the format *pn@GENE*), followed by 1255 columns with non–cancer CAGE scores and 476 columns with cancer CAGE scores and the last column contains the TSS *d* value. Sample names correspond to sample IDs as mentioned in table S4. CAGE raw scores include only those TSSs which were common to all *N* and *C* samples.

**Table S6**. The table contains transcript count index (*TCI*) and base count index (B*C*I) for 55048 TSSs in *N* (*TCI*_*N*_ and B*C*I_*N*_) and *C* (*TCI*_*C*_ and B*C*I_*C*_ ) sample sets.

**Table S7. Tabulation of the GOrilla gene ontology for terminal quartile comparisons of *vd*.** Column headers *GO Term, Description, p–value, FDR q–value* and *Enrichment* were obtained directly from the results generated by the software (PMID 19192299 and 17381235 and see methods for details). The header *Assigned term* corresponds to a group of related functional categories. The column *Comparison* indicates the target versus background. Q1 and Q4 correspond to the quartiles 1 and 4 respectively when sorted by log_2_ *vd* _*C*_/*vd*_*N*_ . *Ontology* indicates the biological process (BP) or molecular function (MF) categories.

**Table S8**. The table contains gene–space entropies of 11928 genes for *N* (*gH*_*N*_ ) and *C* (*gH*_*C*_ ) and the log_2_ *C*/*N* ratio of the entropies.

**Table S9. Tabulation of the GOrilla gene ontology for terminal quartile comparisons of *gH*.** Column headers *GO Term, Description, p–value, FDR q–value* and *Enrichment* were obtained directly from the results generated by the software (PMID 19192299 and 17381235 and see methods for details). The header *Assigned term* corresponds to a group of related functional categories. The column *Comparison* indicates the target versus background. Q1 and Q4 correspond to the quartiles 1 and 4 respectively when sorted by log_2_ *gH* _*C*_/*gH*_*N*_. *Ontology* indicates the biological process (BP) or molecular function (MF) categories.

**Table S10. The table contains sample–space entropy (***sH***) for each TSS calculated two ways. *sH* from non–zero scores**: *sH* for a TSS from randomly selected 100 samples such that the 100 samples have a score greater than 0 in *N* as well as *C* and their log_2_ *sH*_*C*_ /*sH* _*N*_. TSSs that did not satisfy these criteria could not have their *sH* calculated and are marked NA. *sH* from all scores: *sH* for a TSS from randomly selected 100 samples in *N* and *C* and their log_2_ ratios. TSSs that could be used to calculate *sH* in both ways were considered *frequently deployed* while the other TSSs were labelled *non–frequently deployed*.

**Table S11. The table contains the product of parameters *E*_*t*_, *TCI* or *d* and *sH* calculated two ways for 55048 TSSs categorized as *frequently* or *non–frequently deployed***. Columns labelled “*sH* from non–zero scores” indicate *sH* calculation from samples that have scores greater than 0. Columns labelled “*sH* from all scores” indicate no filtering of scores when calculating *sH*.

**Table S12. Tabulation of the GOrilla gene ontology for terminal quartile comparisons of *sH*.** Since *sH* is a TSS–level parameter, the TSSs segregating into the two quartiles were grouped by their specific genes. Column headers *GO Term, Description, p–value, FDR q– value* and *Enrichment* were obtained directly from the results generated by the software (PMID 19192299 and 17381235 and see methods for details). The header *Assigned term* corresponds to a group of related functional categories. The column *Comparison* indicates the target versus background. Q1 and Q4 correspond to the quartiles 1 and 4 respectively when sorted by log_2_ *sH*_*C*_ /*sH*_*N*_ . *Ontology* indicates the biological process (BP) or molecular function (MF) categories.

**Table S13**. The table contains compound entropy, its *vd*–based derivative *vd*/*cH* and, log_2_ *C*/*N* ratios of the two for 11928 genes.

**Table S14. Tabulation of the GOrilla gene ontology for terminal quartile comparisons of *vd*/*cH*.** Column headers *GO Term, Description, p–value, FDR q–value* and *Enrichment* were obtained directly from the results generated by the software (PMID 19192299 and 17381235 and see methods for details). The header *Assigned term* corresponds to a group of related functional categories. The column *Comparison* indicates the target versus background. Q1 and Q4 correspond to the quartiles 1 and 4 when sorted by log_2_ ratio of *vd* _*C*_/*cH*_*C*_ and *vd*_*N*_ /*cH*_*N*_. *Ontology* indicates the biological process (BP) or molecular function (MF) analysis.

**Table S15**. Coefficients of the exponential growth function with rate constant parameter for each of the assigned *d* bins. *y*0 is the offset, *A* is the initial value and, *R*0 is the rate constant.

