## Supplementary material for "Gene functions determine stochastic or adaptive futile transcription in cancers through deregulated start site deployments": Fig S1 to S26

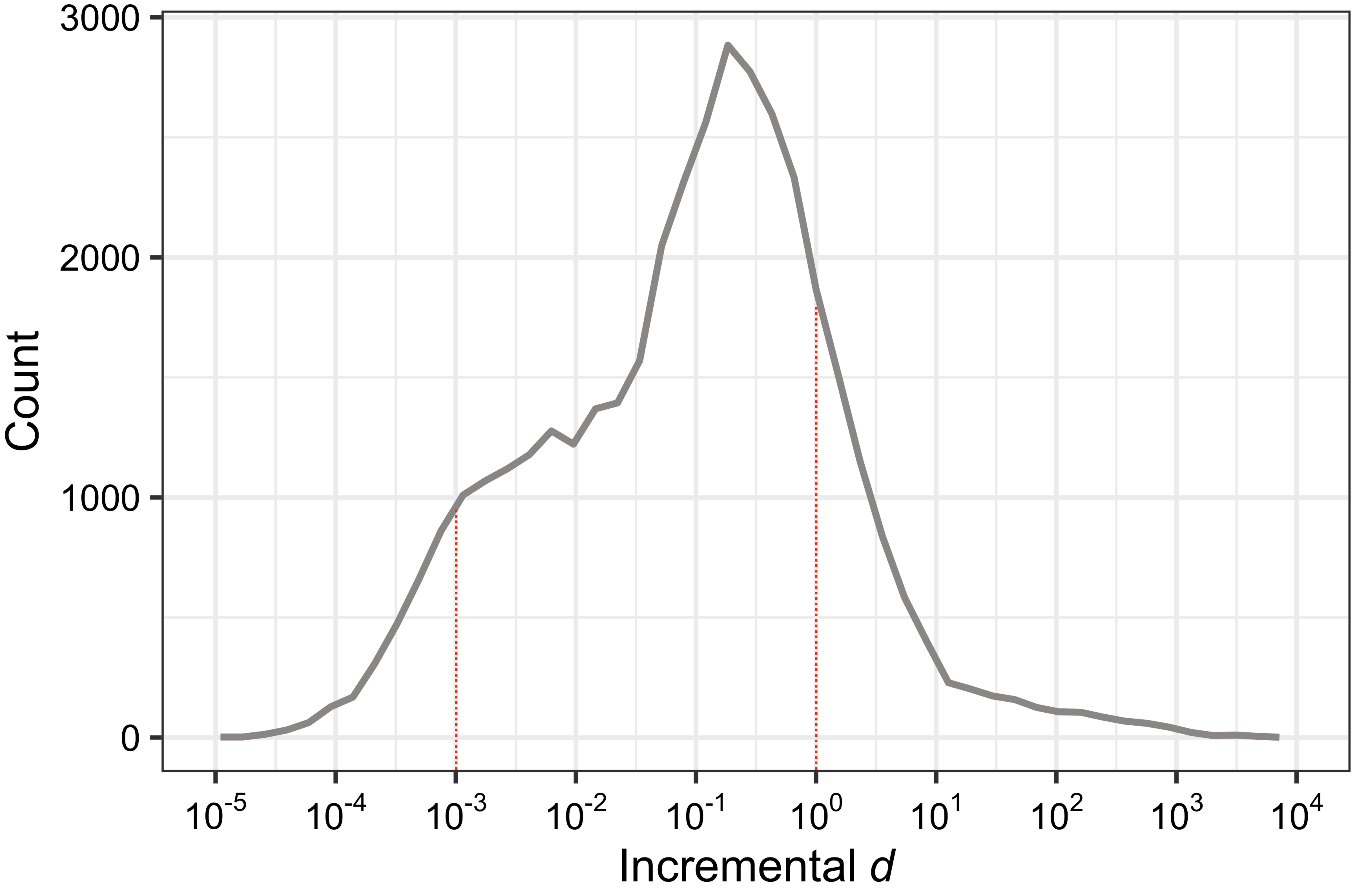

3000
2000
1000
0
Count
10-5
10-4
10-3
10-2
10-1
100
101
102
103
104
Incremental d

Fig S2

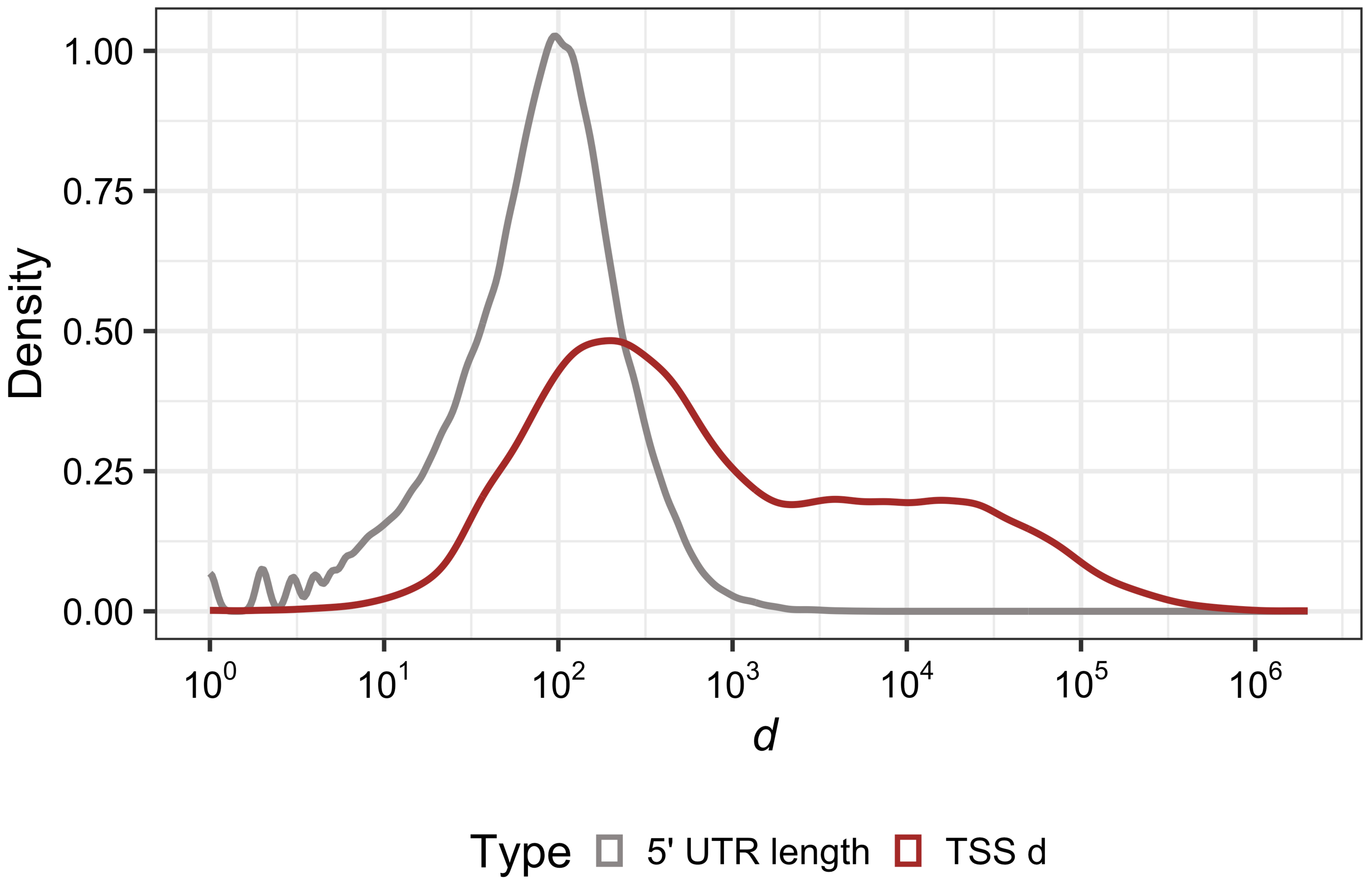

Density
1.00
0.75
0.50
0.25
0.00
$10^0$
$10^1$
$10^2$
$10^3$
$10^4$
$10^5$
$10^6$
*d*
Type
5' UTR length
TSS d

Fig S3

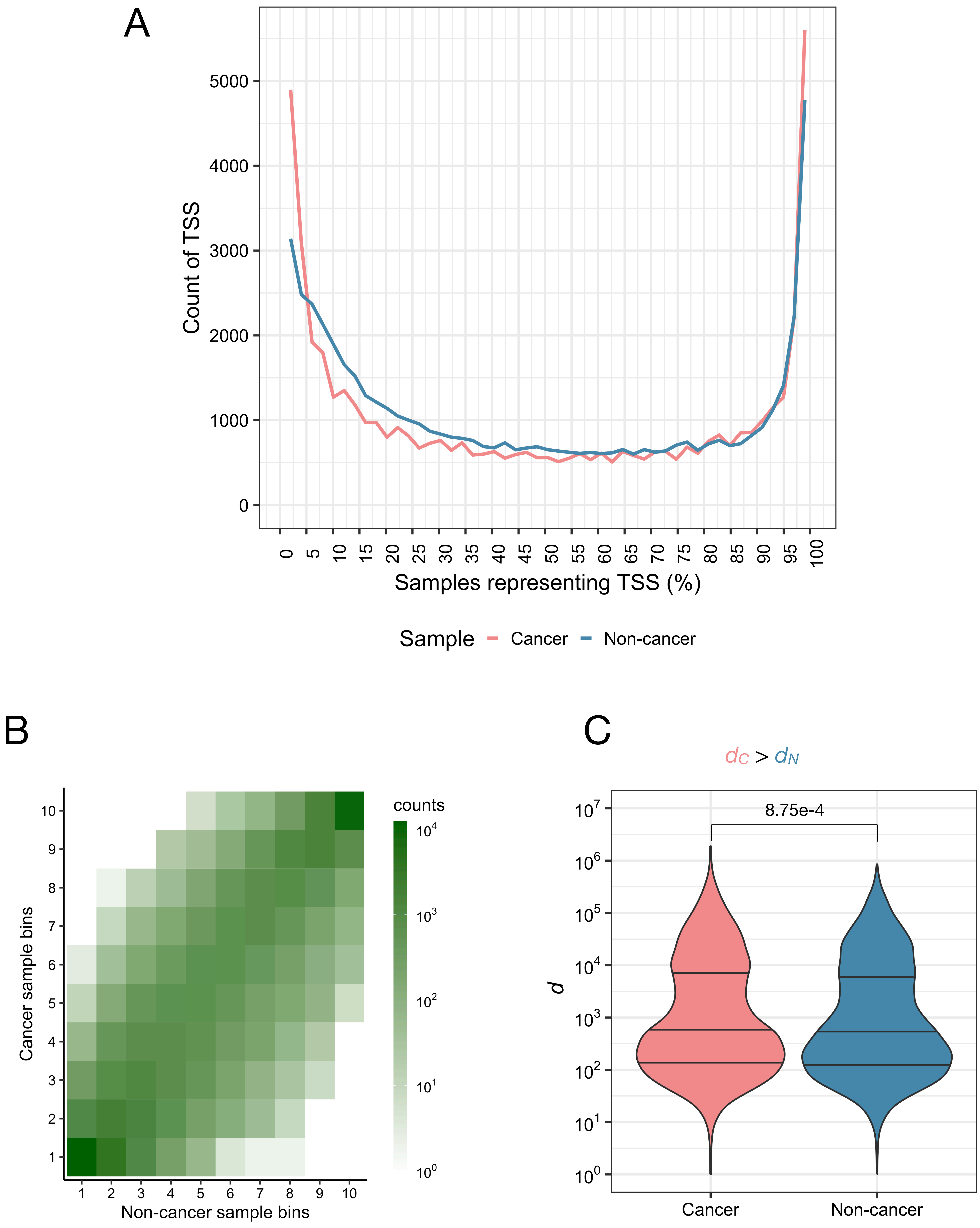

A
Count of TSS
Samples representing TSS (%)
Sample
Cancer
Non-cancer
B
Cancer sample bins
Non-cancer sample bins
counts
C
$d_C > d_N$
8.75e-4
d
Cancer
Non-cancer

Fig S4

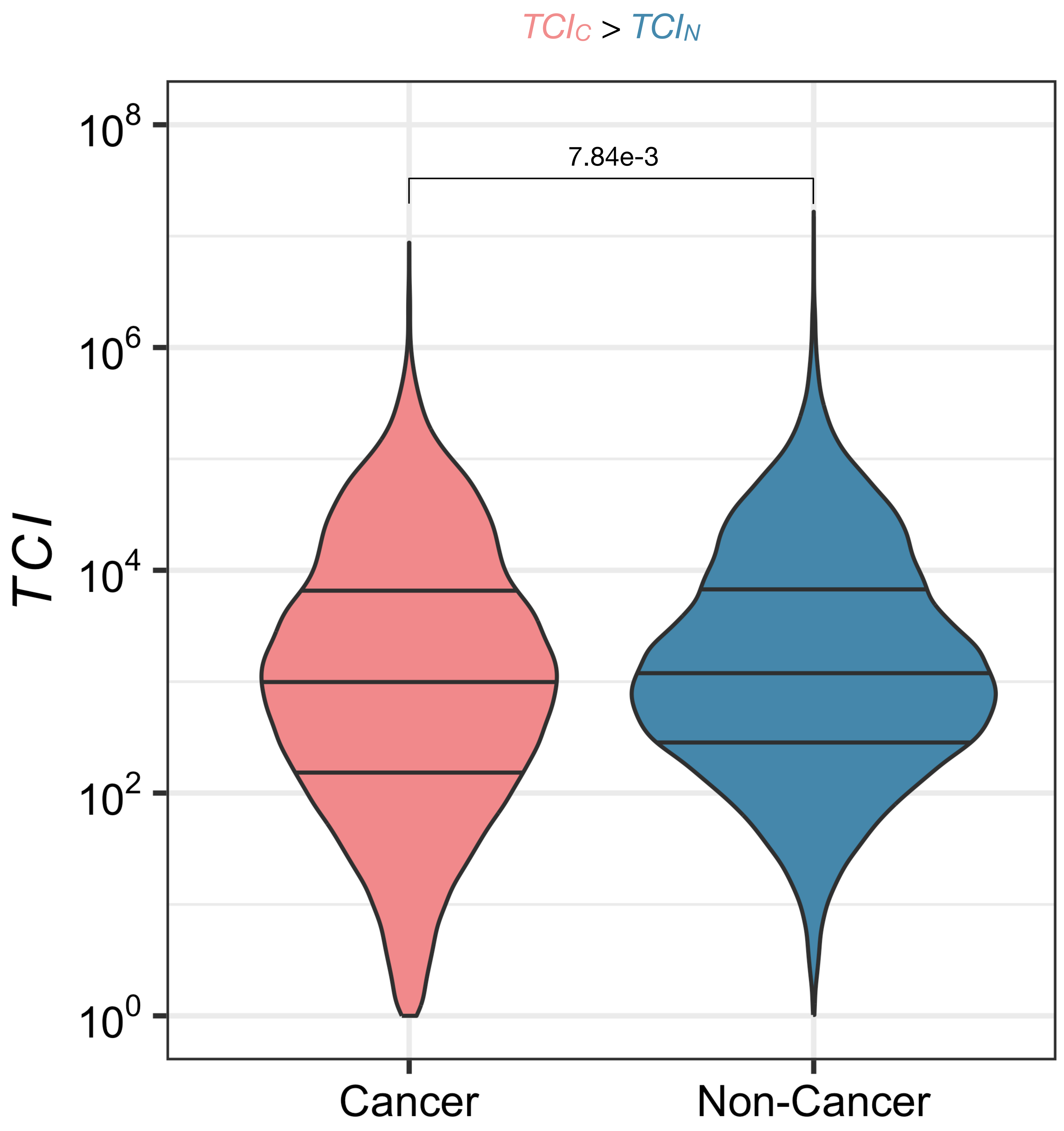

### Fig S5

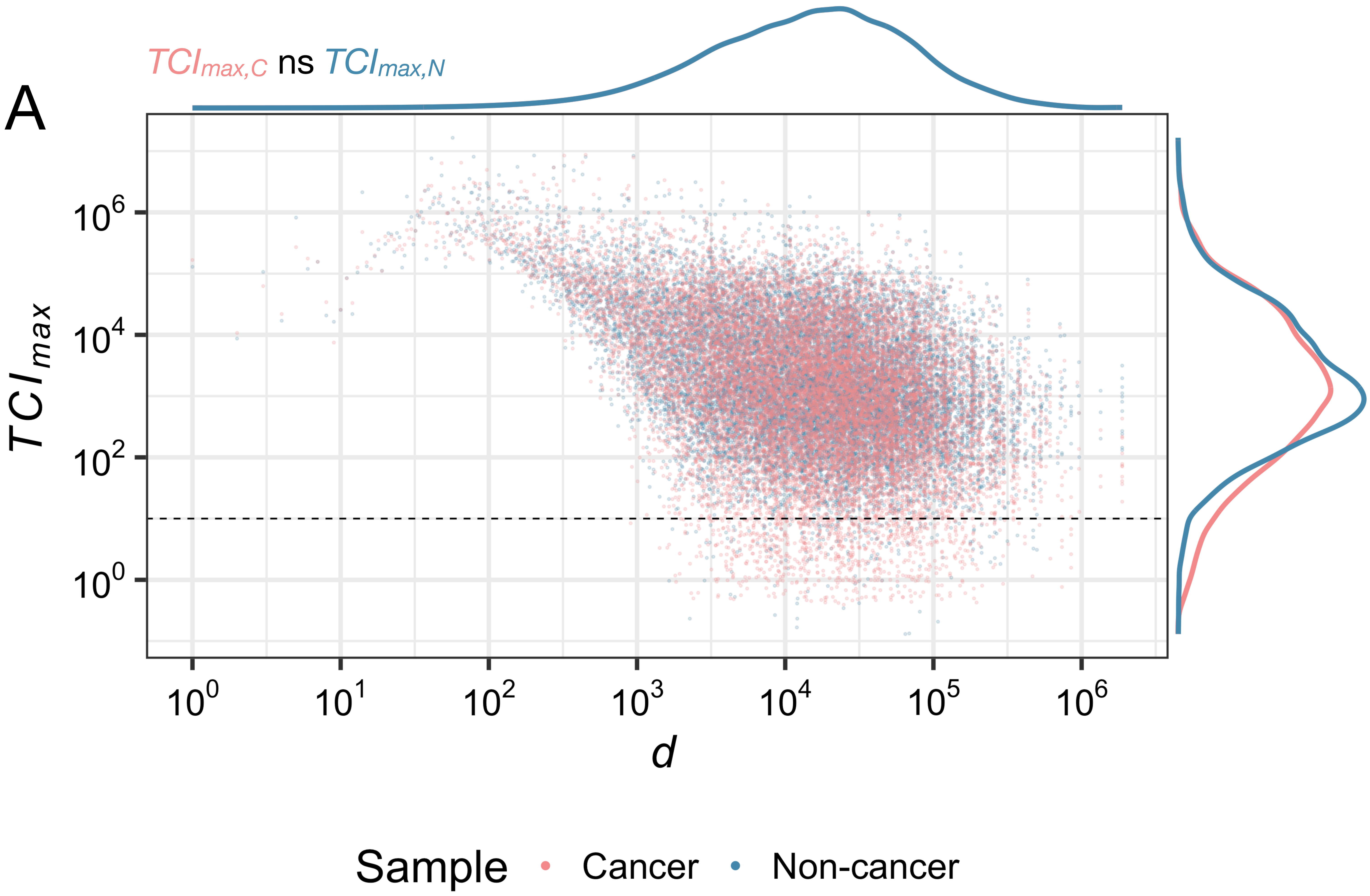

A
TCImax,C ns TCImax,N
TCImax
10^6
10^4
10^2
10^0
10^0 10^1 10^2 10^3 10^4 10^5 10^6
d
Sample Cancer Non-cancer


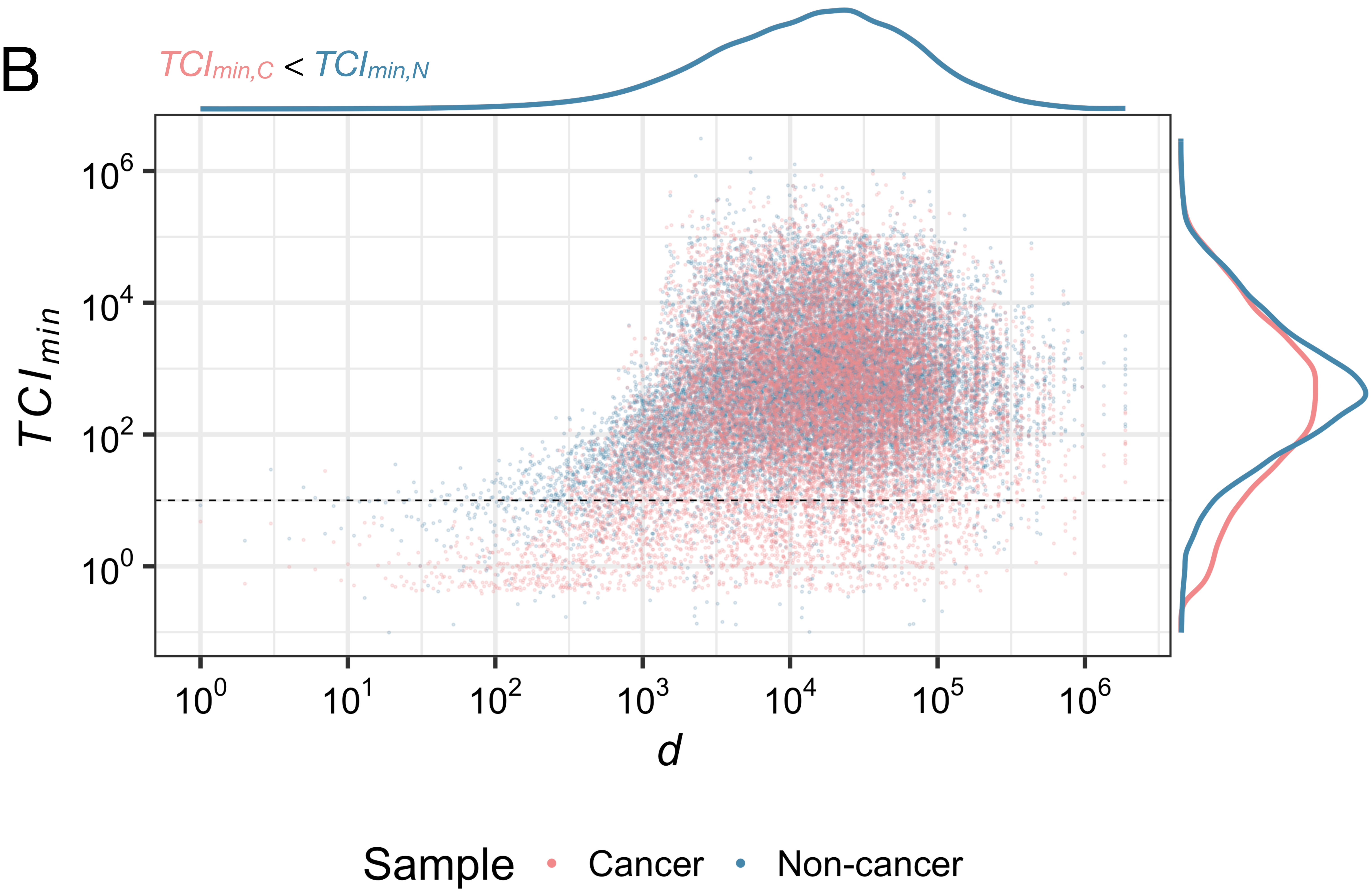

B
TCImin,C < TCImin,N
TCImin
10^6
10^4
10^2
10^0
10^0 10^1 10^2 10^3 10^4 10^5 10^6
d
Sample Cancer Non-cancer

### Fig S6

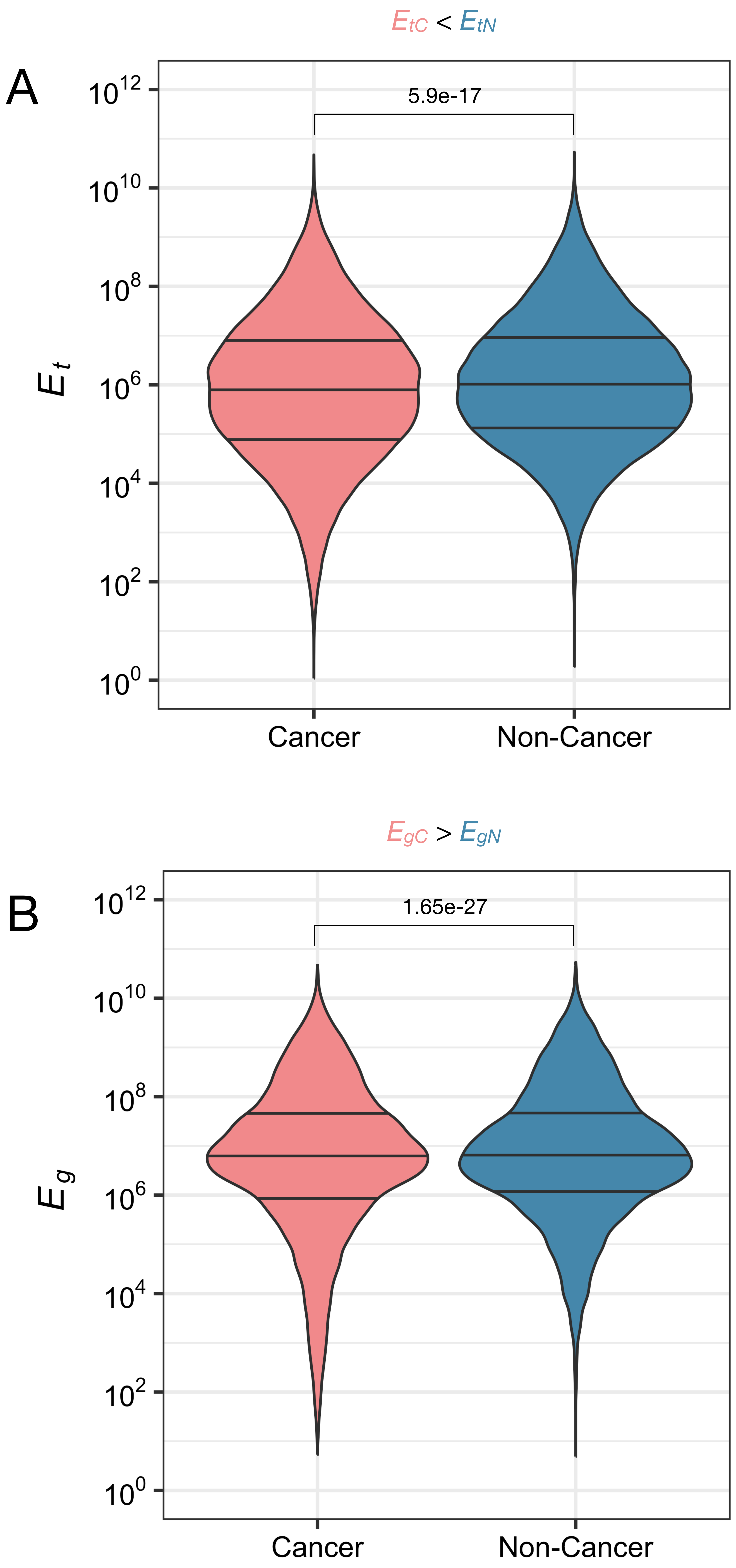

$E_{tC} < E_{tN}$
A
5.9e-17
$E_t$
$10^{12}$
$10^{10}$
$10^8$
$10^6$
$10^4$
$10^2$
$10^0$
Cancer
Non-Cancer
$E_{gC} > E_{gN}$
B
1.65e-27
$E_g$
$10^{12}$
$10^{10}$
$10^8$
$10^6$
$10^4$
$10^2$
$10^0$
Cancer
Non-Cancer

Fig S7

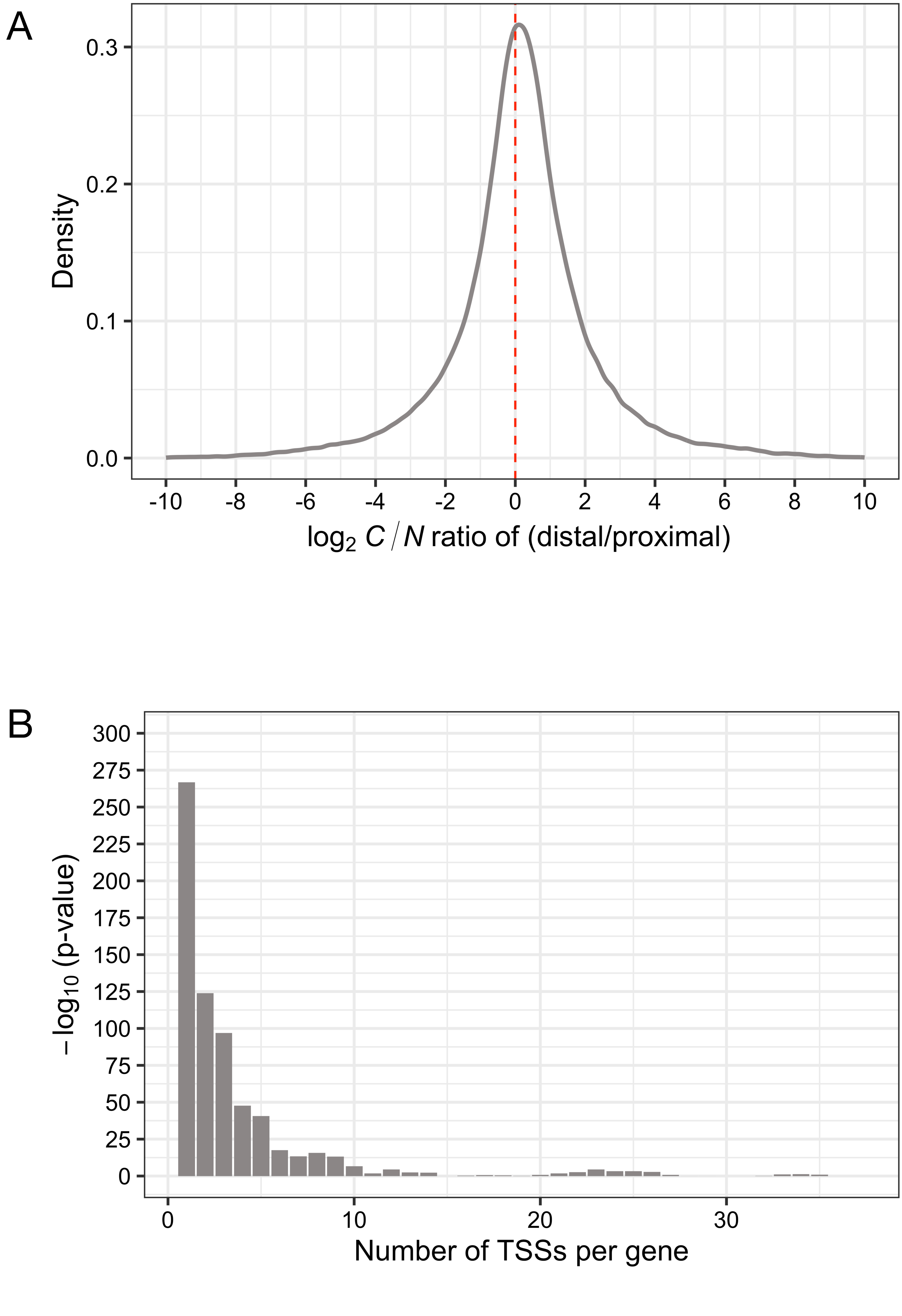
A
0.3
0.2
0.1
0.0
Density
-10
-8
-6
-4
-2
0
2
4
6
8
10
log2 C / N ratio of (distal/proximal)
B
300
275
250
225
200
175
150
125
100
75
50
25
0
−log10 (p-value)
0
10
20
30
Number of TSSs per gene

Fig S8

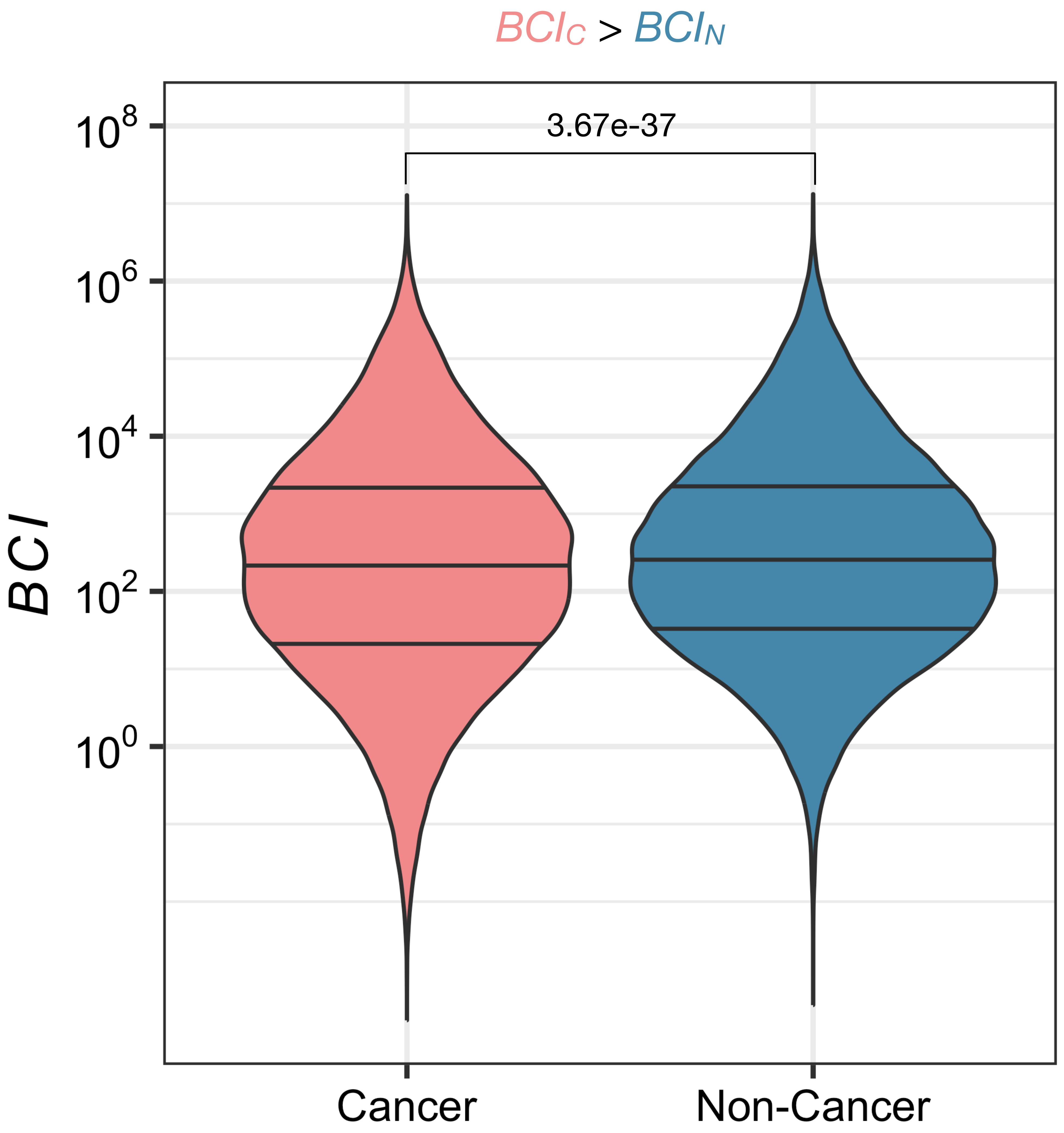

Fig S9

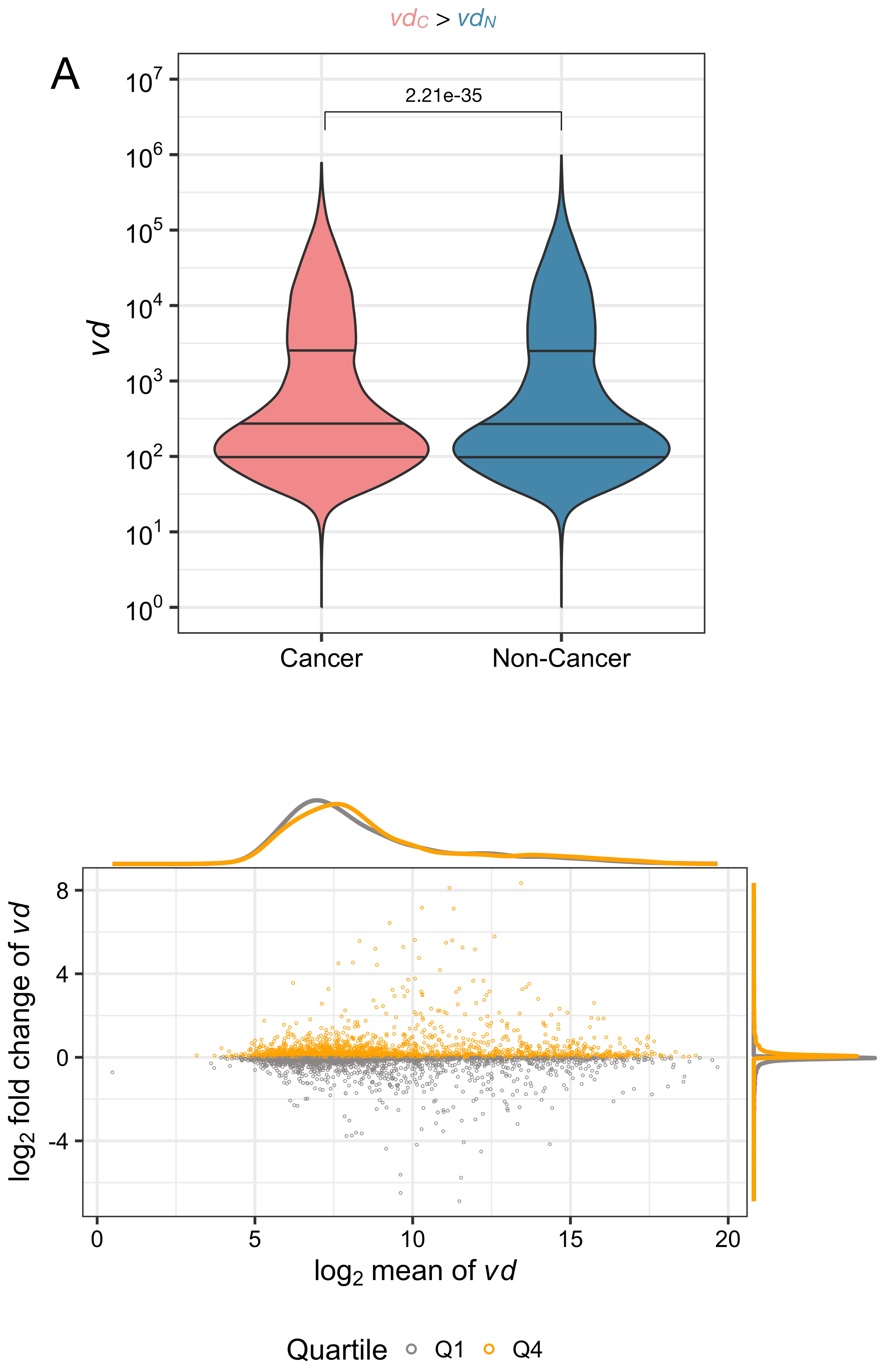

$vd_C$ > $vd_N$
A
2.21e-35
$10^7$
$10^6$
$10^5$
$10^4$
$10^3$
$10^2$
$10^1$
$10^0$
vd
Cancer
Non-Cancer
B
8
4
0
-4
log2 fold change of vd
0
5
10
15
20
log2 mean of vd
Quartile
Q1
Q4

Fig S10

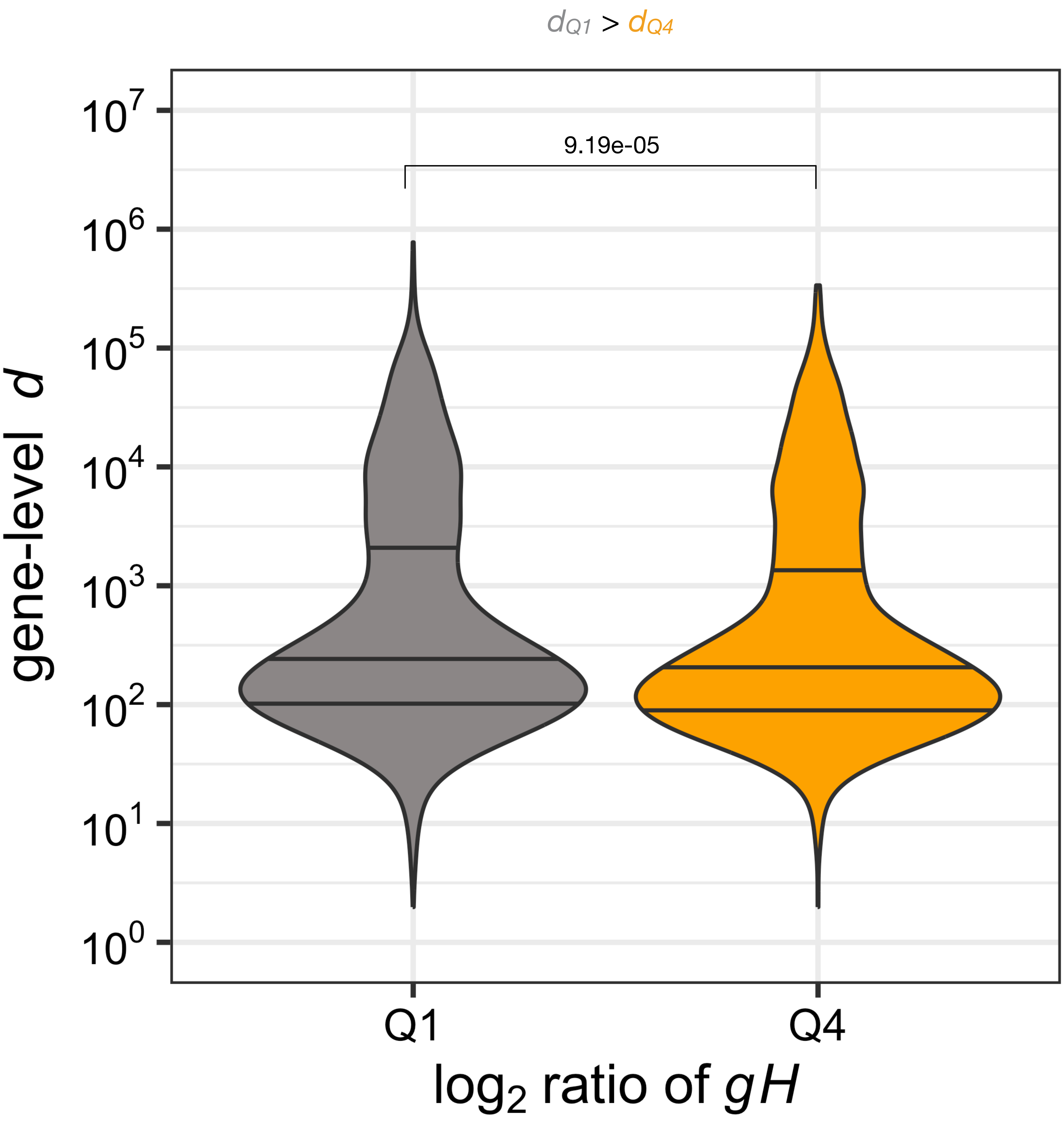

$d_{Q1} > d_{Q4}$
9.19e-05
gene-level d
$10^7$
$10^6$
$10^5$
$10^4$
$10^3$
$10^2$
$10^1$
$10^0$
Q1
Q4
$\log_2$ ratio of $gH$

### Fig S11

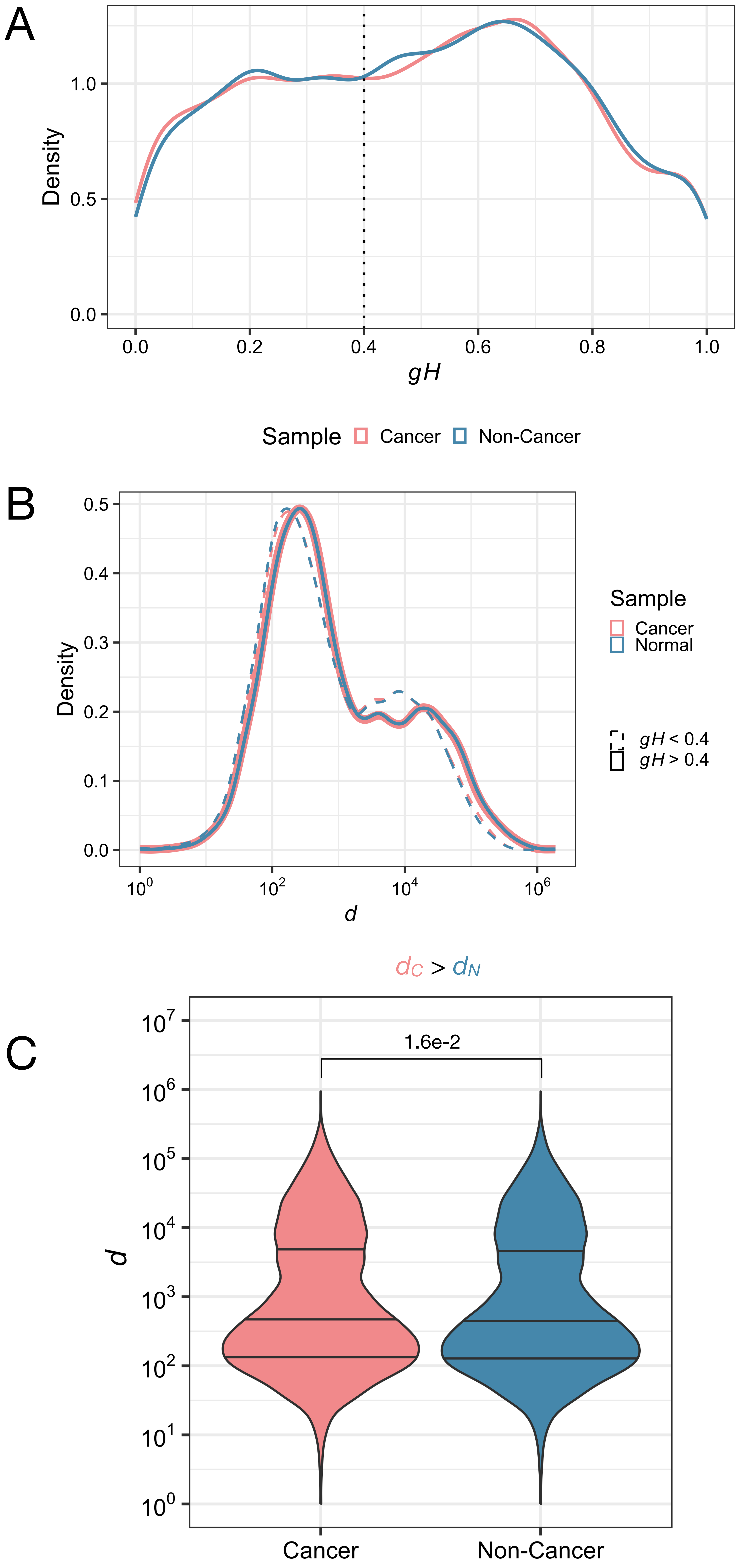
A
Density
1.0
0.5
0.0
0.0
0.2
0.4
0.6
0.8
1.0
gH
Sample
Cancer
Non-Cancer
B
0.5
0.4
0.3
0.2
0.1
0.0
Density
10^0
10^2
10^4
10^6
d
Sample
Cancer
Normal
gH < 0.4
gH > 0.4
dC > dN
C
1.6e-2
10^7
10^6
10^5
10^4
10^3
10^2
10^1
10^0
d
Cancer
Non-Cancer

Fig S12

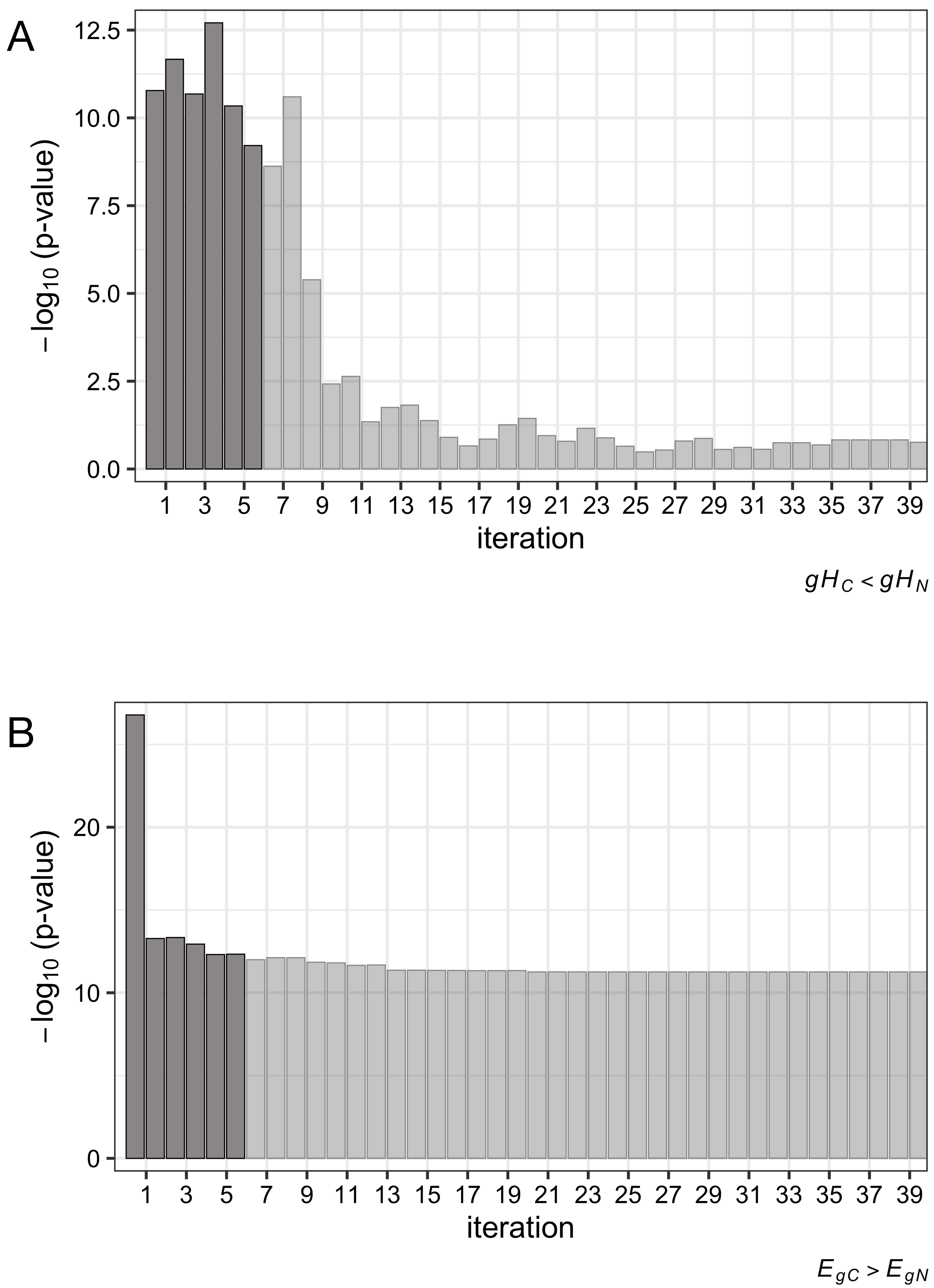
A
12.5
10.0
7.5
5.0
2.5
0.0
$-\log_{10}$ (p-value)
1 3 5 7 9 11 13 15 17 19 21 23 25 27 29 31 33 35 37 39
iteration
$gH_C < gH_N$
B
20
10
0
$-\log_{10}$ (p-value)
1 3 5 7 9 11 13 15 17 19 21 23 25 27 29 31 33 35 37 39
iteration
$E_{gC} > E_{gN}$

Fig S13

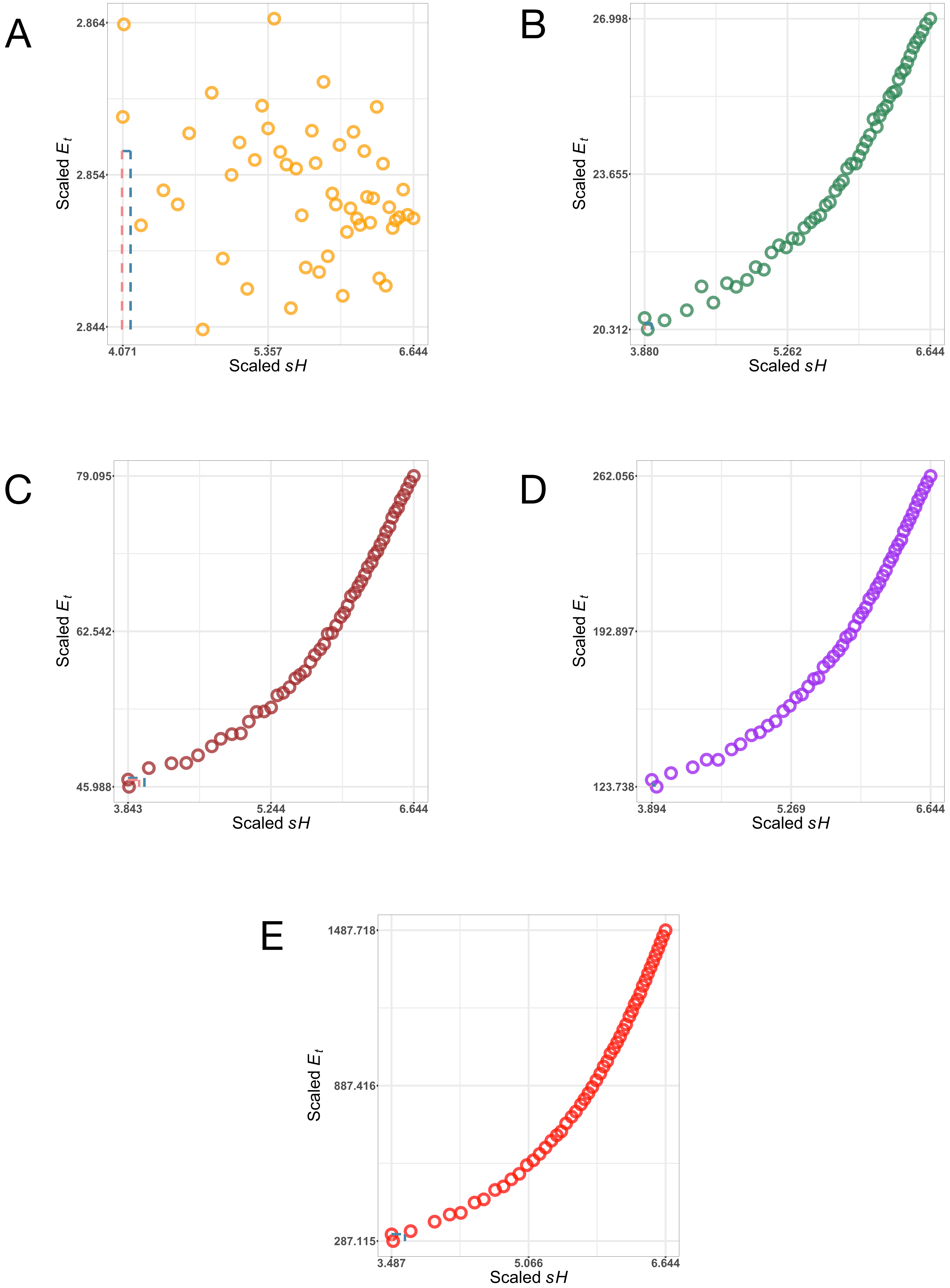

A
2.864
2.854
2.844
Scaled $E_t$
4.071
5.357
6.644
Scaled $sH$
B
26.998
23.655
20.312
Scaled $E_t$
3.880
5.262
6.644
Scaled $sH$
C
79.095
62.542
45.988
Scaled $E_t$
3.843
5.244
6.644
Scaled $sH$
D
262.056
192.897
123.738
Scaled $E_t$
3.894
5.269
6.644
Scaled $sH$
E
1487.718
887.416
287.115
Scaled $E_t$
3.487
5.066
6.644
Scaled $sH$

### Fig S14

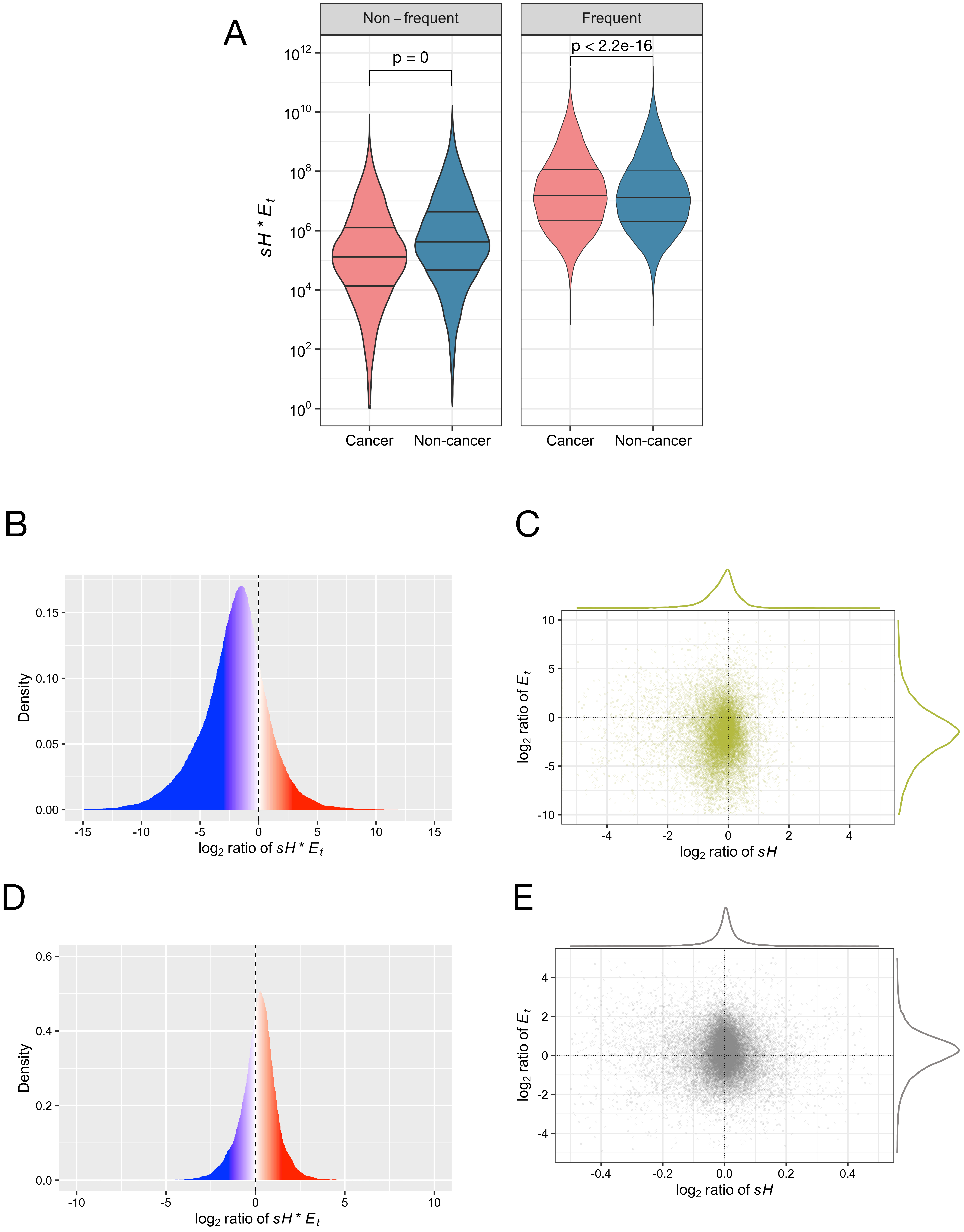
A
Non – frequent
Frequent
p = 0
p < 2.2e-16
sH * E_t
Cancer
Non-cancer
B
Density
log2 ratio of sH * E_t
C
log2 ratio of E_t
log2 ratio of sH
D
Density
log2 ratio of sH * E_t
E
log2 ratio of E_t
log2 ratio of sH

Fig S15

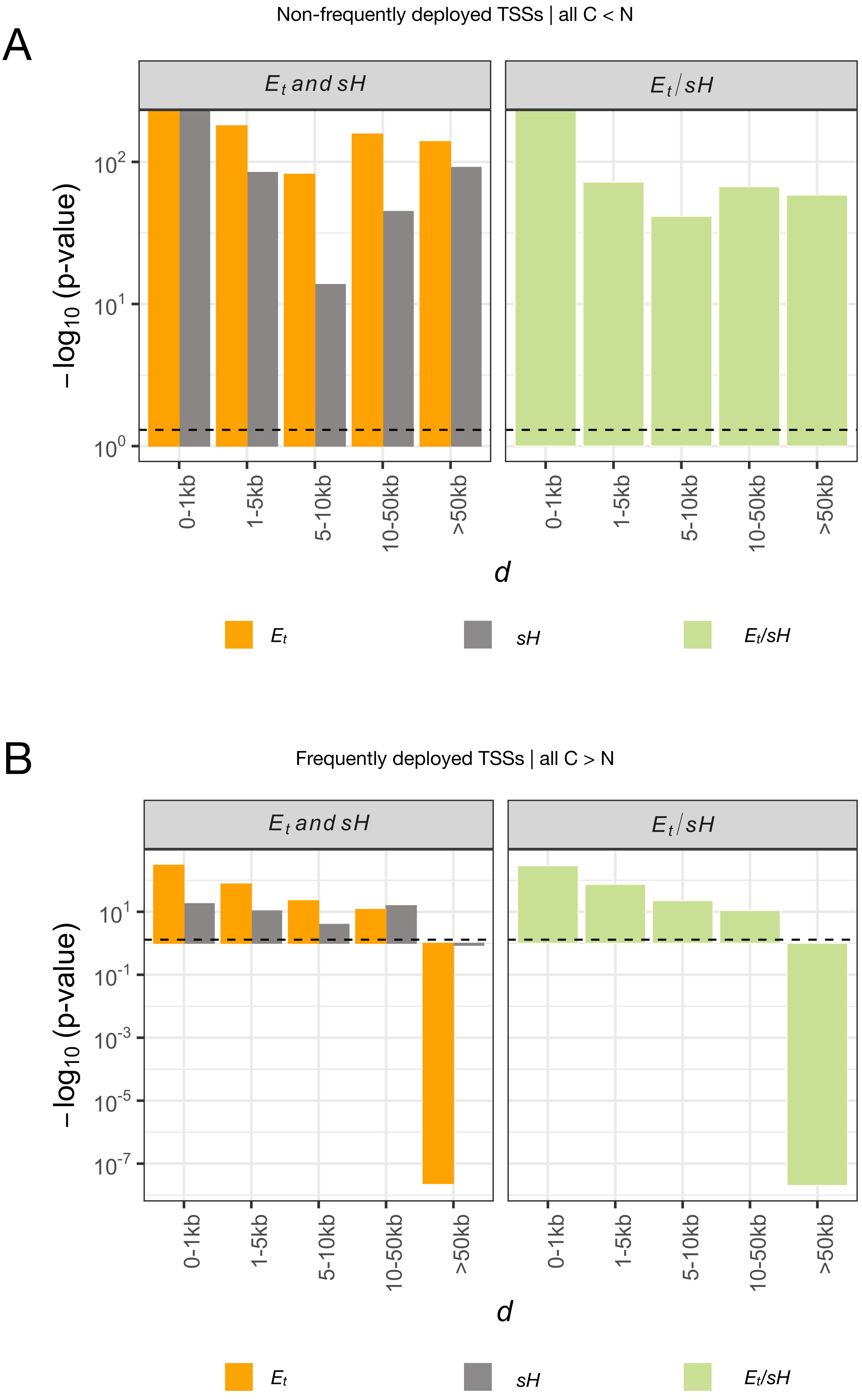
A
Non-frequently deployed TSSs | all C < N
Et and sH
Et / sH
−log10 (p-value)
10^2
10^1
10^0
0-1kb
1-5kb
5-10kb
10-50kb
>50kb
d
Et
sH
Et/sH
B
Frequently deployed TSSs | all C > N
Et and sH
Et / sH
−log10 (p-value)
10^1
10^-1
10^-3
10^-5
10^-7
0-1kb
1-5kb
5-10kb
10-50kb
>50kb
d
Et
sH
Et/sH

### Fig S16

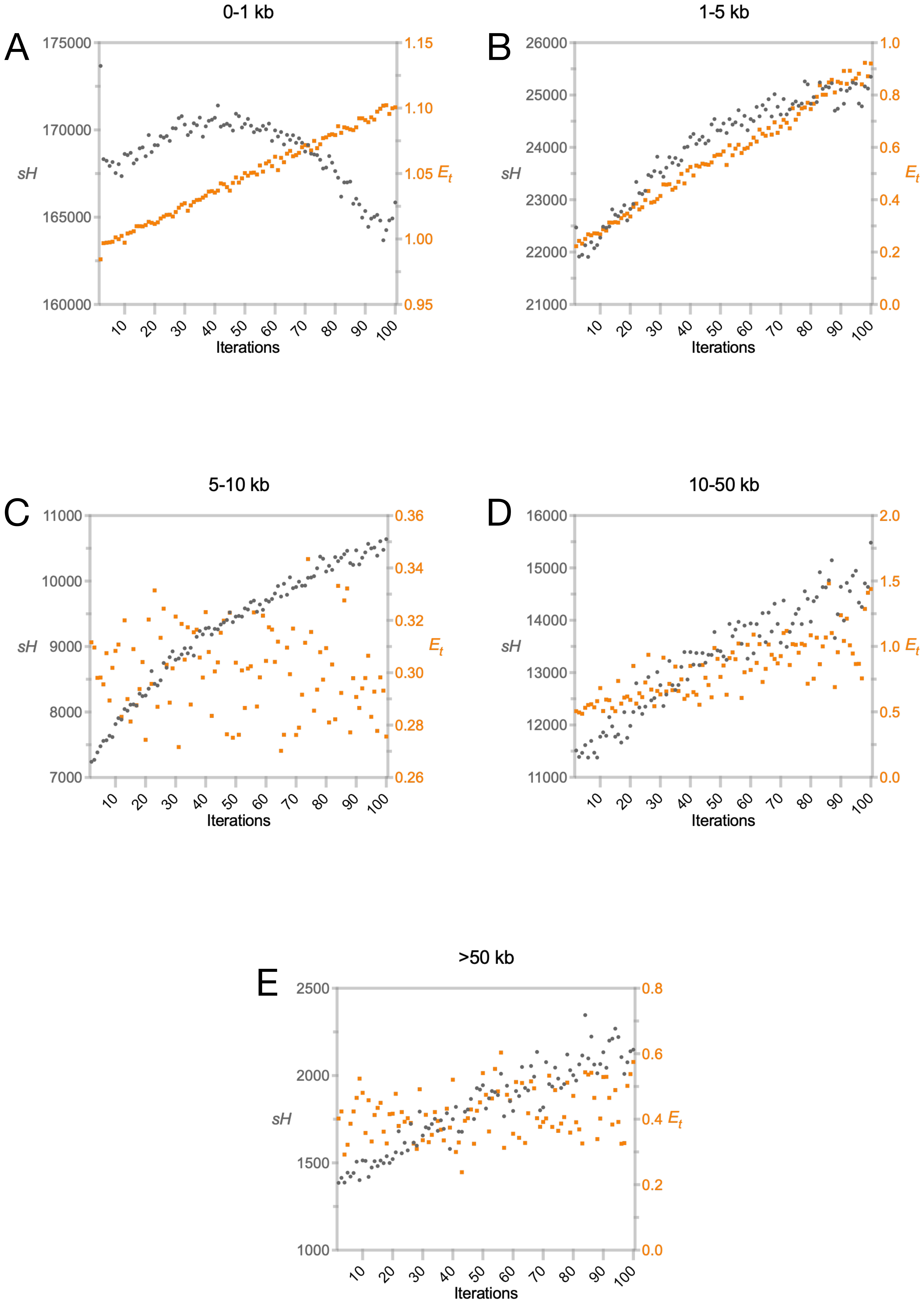

0-1 kb
A
175000
170000
165000
160000
sH
1.15
1.10
1.05 $E_t$
1.00
0.95
10
20
30
40
50
60
70
80
90
100
Iterations
1-5 kb
B
26000
25000
24000
23000
22000
21000
sH
1.0
0.8
0.6
$E_t$
0.4
0.2
0.0
Iterations
5-10 kb
C
11000
10000
9000
8000
7000
sH
0.36
0.34
0.32
$E_t$
0.30
0.28
0.26
Iterations
10-50 kb
D
16000
15000
14000
13000
12000
11000
sH
2.0
1.5
1.0 $E_t$
0.5
0.0
Iterations
>50 kb
E
2500
2000
1500
1000
sH
0.8
0.6
0.4 $E_t$
0.2
0.0
Iterations

Fig S17

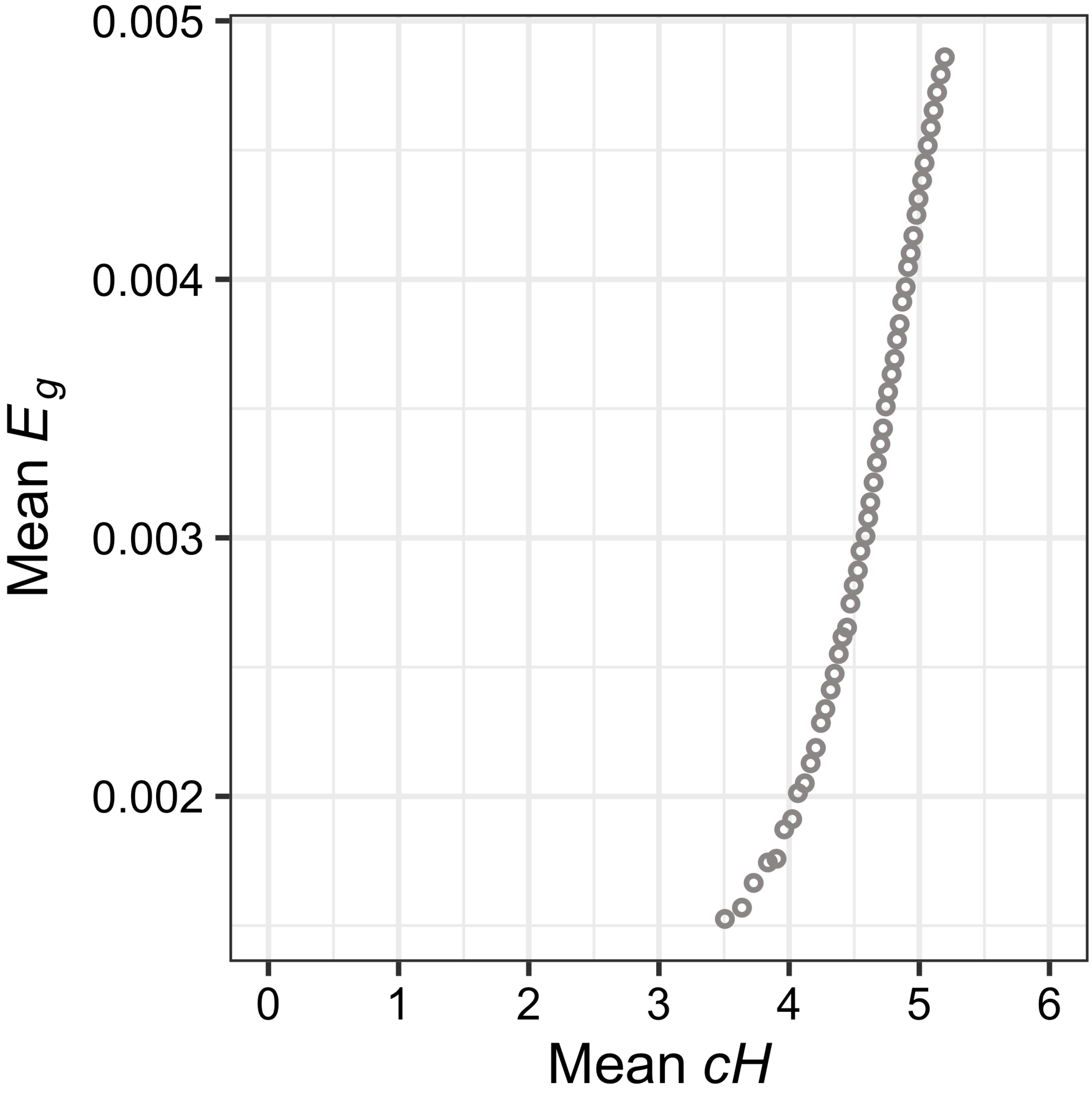

Fig S18

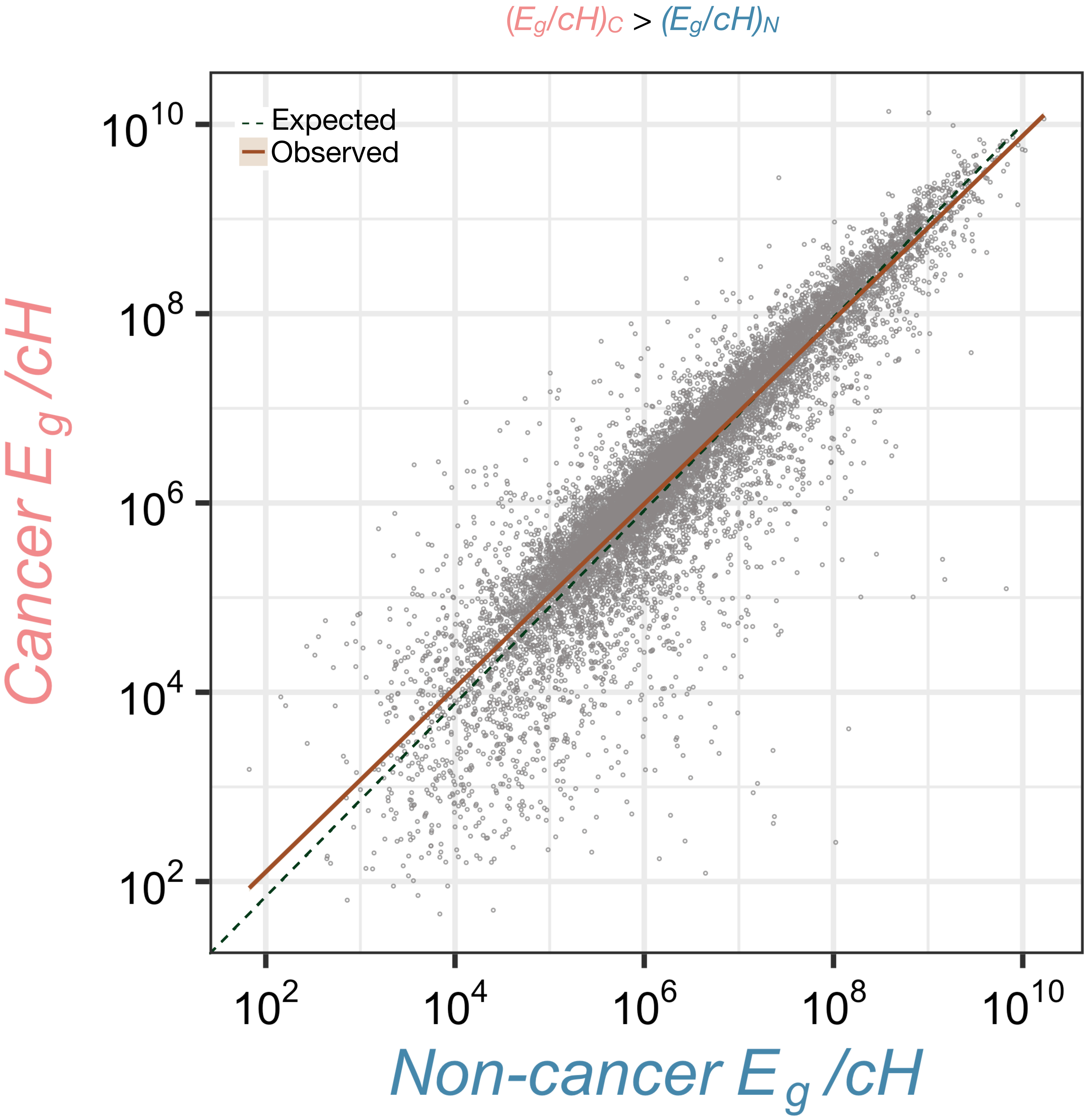

$(E_g/cH)_C > (E_g/cH)_N$
Expected
Observed
$10^{10}$
$10^{8}$
$10^{6}$
$10^{4}$
$10^{2}$
Cancer $E_g$ /cH
Non-cancer $E_g$ /cH

Fig S19

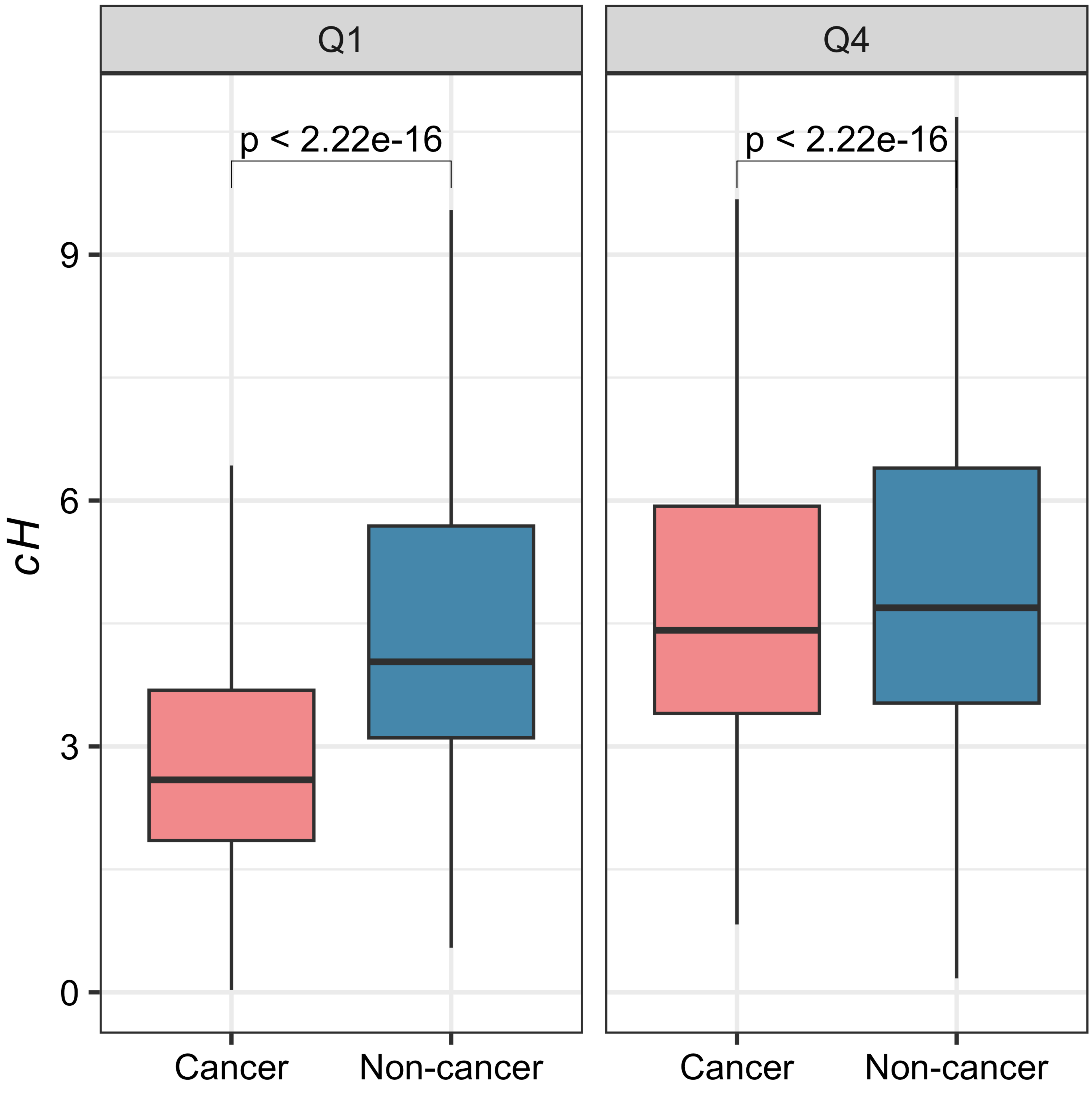

Q1
Q4
p < 2.22e-16
p < 2.22e-16
cH
9
6
3
0
Cancer
Non-cancer
Cancer
Non-cancer

Fig S20

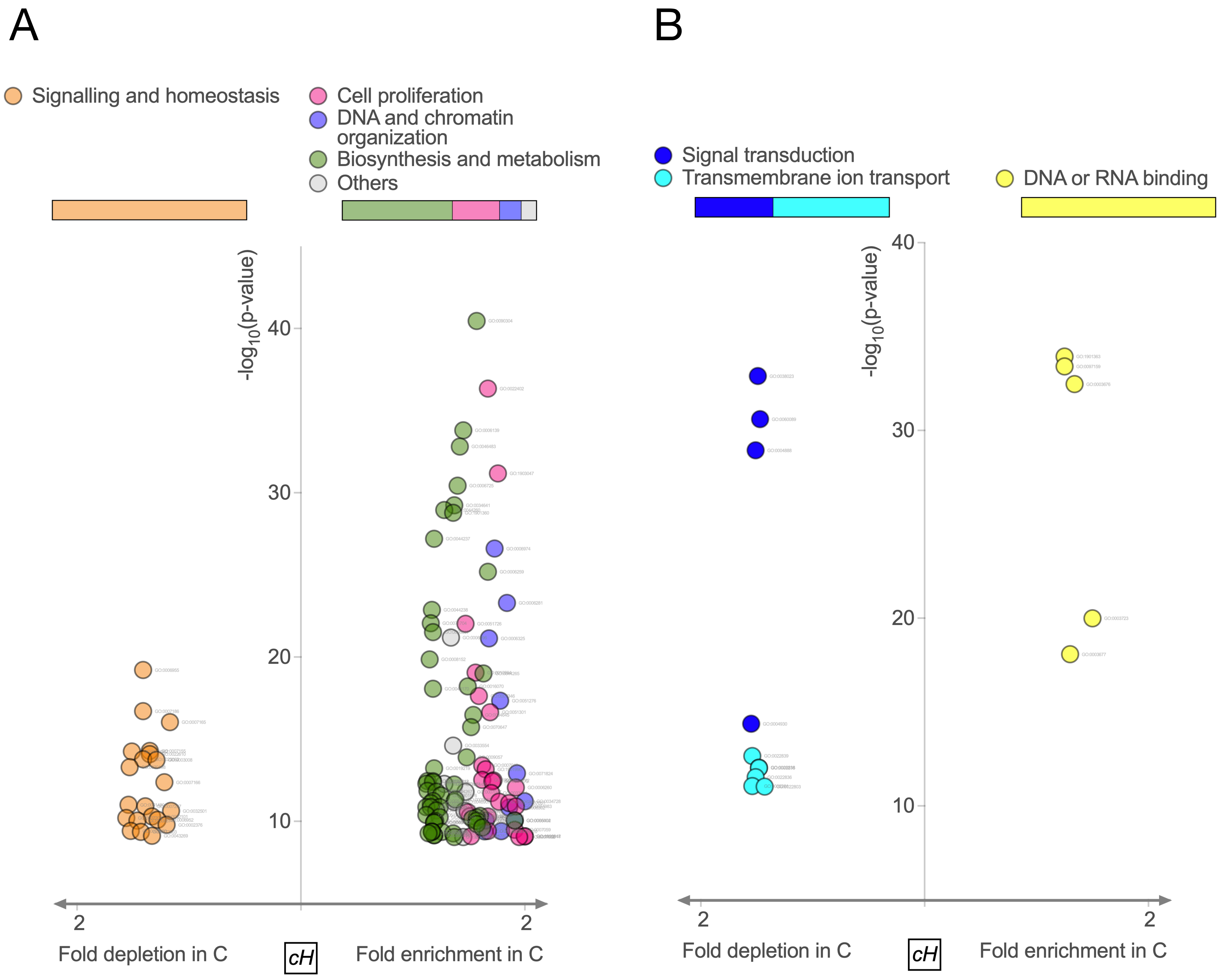
A
Signalling and homeostasis
Cell proliferation
DNA and chromatin organization
Biosynthesis and metabolism
Others
$-\log_{10}$(p-value)
40
30
20
10
2
2
Fold depletion in C
cH
Fold enrichment in C
B
Signal transduction
Transmembrane ion transport
DNA or RNA binding
$-\log_{10}$(p-value)
40
30
20
10
2
2
Fold depletion in C
cH
Fold enrichment in C

Fig S21

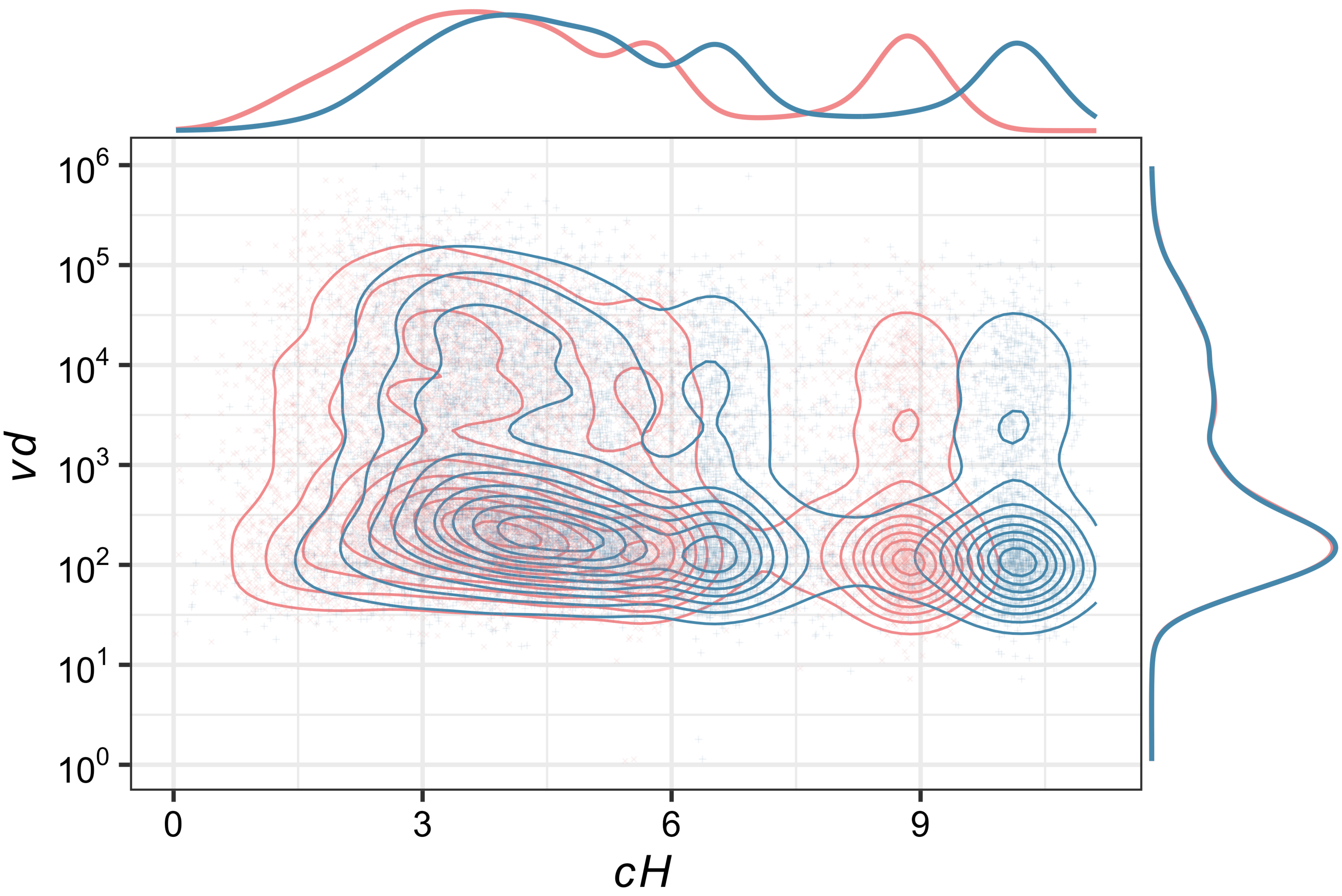

Fig S22

A

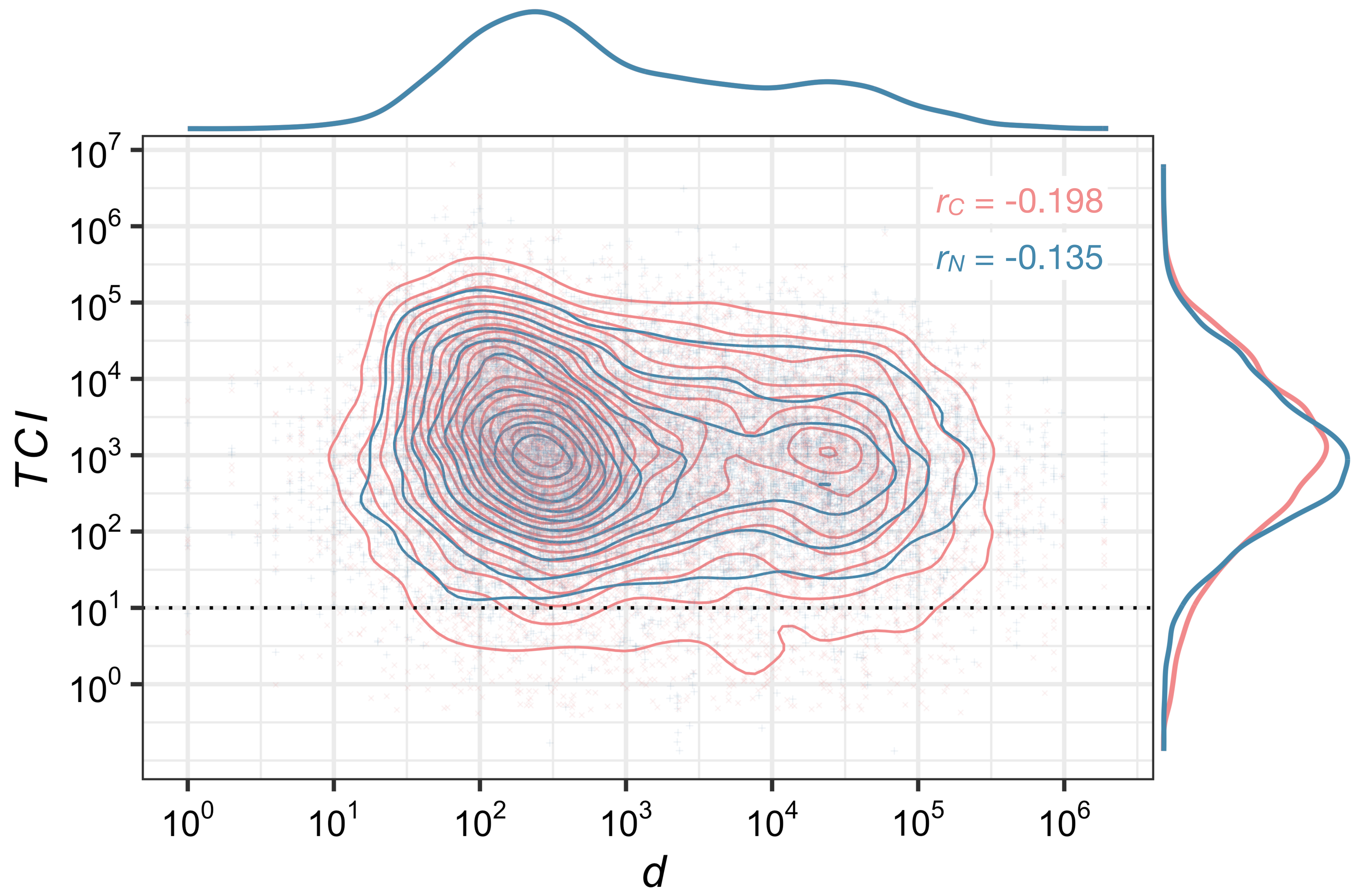


B

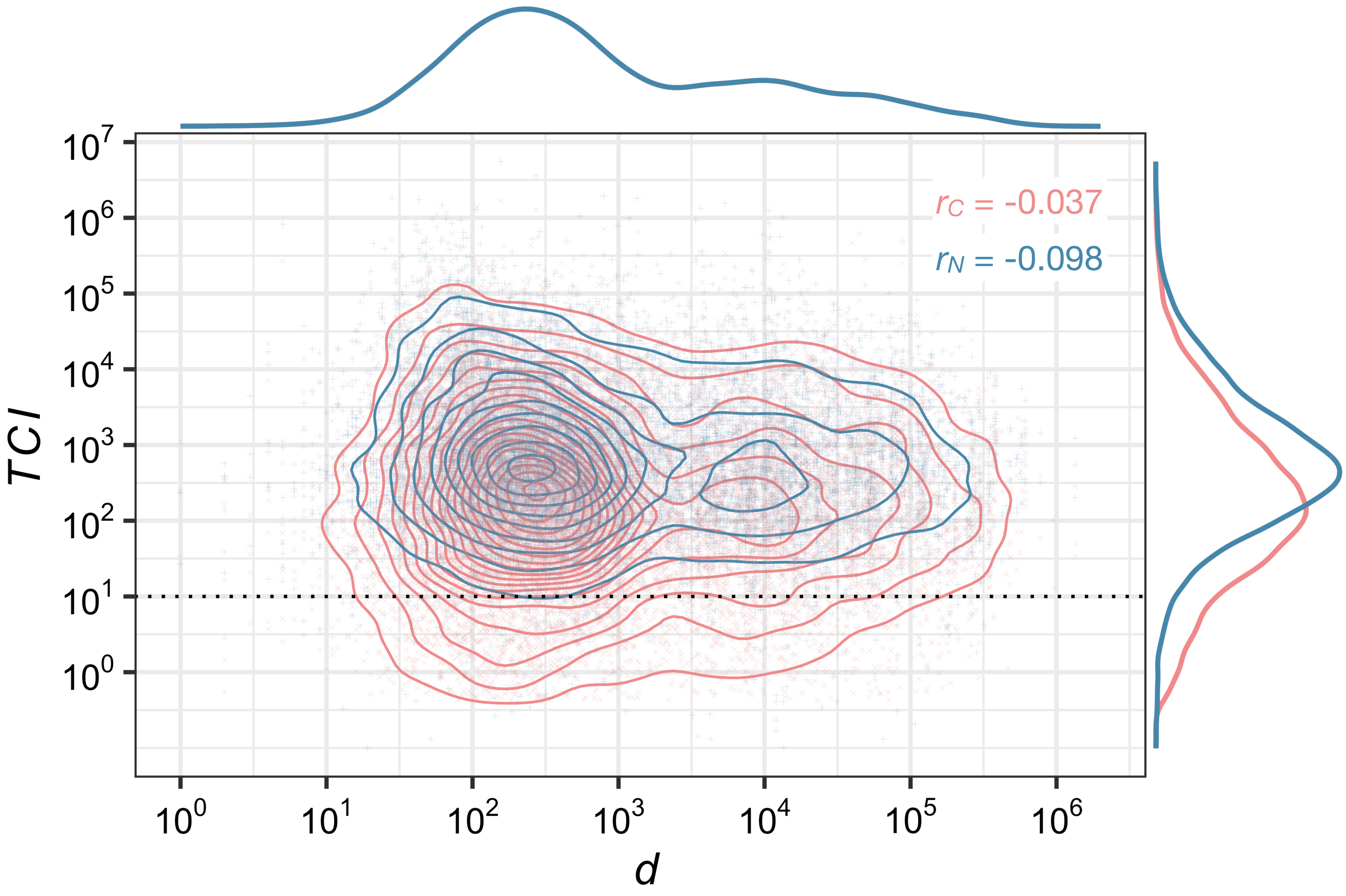

### Fig S23

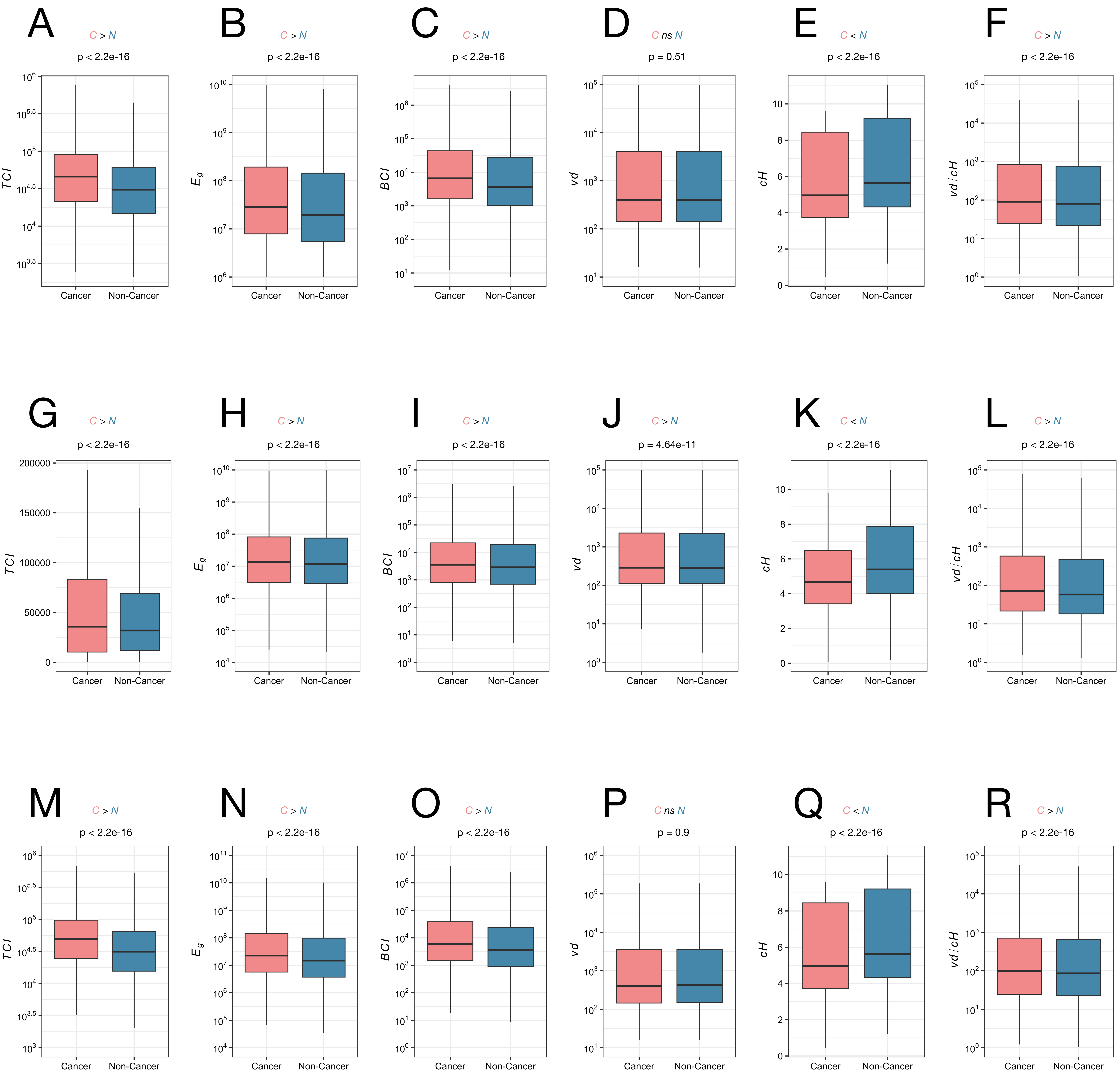
A
C > N
p < 2.2e-16
TCI
Cancer
Non-Cancer
B
C > N
p < 2.2e-16
Eg
C
C > N
p < 2.2e-16
BCI
D
C ns N
p = 0.51
vd
E
C < N
p < 2.2e-16
cH
F
C > N
p < 2.2e-16
vd / cH
G
C > N
p < 2.2e-16
TCI
H
C > N
p < 2.2e-16
Eg
I
C > N
p < 2.2e-16
BCI
J
C > N
p = 4.64e-11
vd
K
C < N
p < 2.2e-16
cH
L
C > N
p < 2.2e-16
vd / cH
M
C > N
p < 2.2e-16
TCI
N
C > N
p < 2.2e-16
Eg
O
C > N
p < 2.2e-16
BCI
P
C ns N
p = 0.9
vd
Q
C < N
p < 2.2e-16
cH
R
C > N
p < 2.2e-16
vd / cH

Fig S24

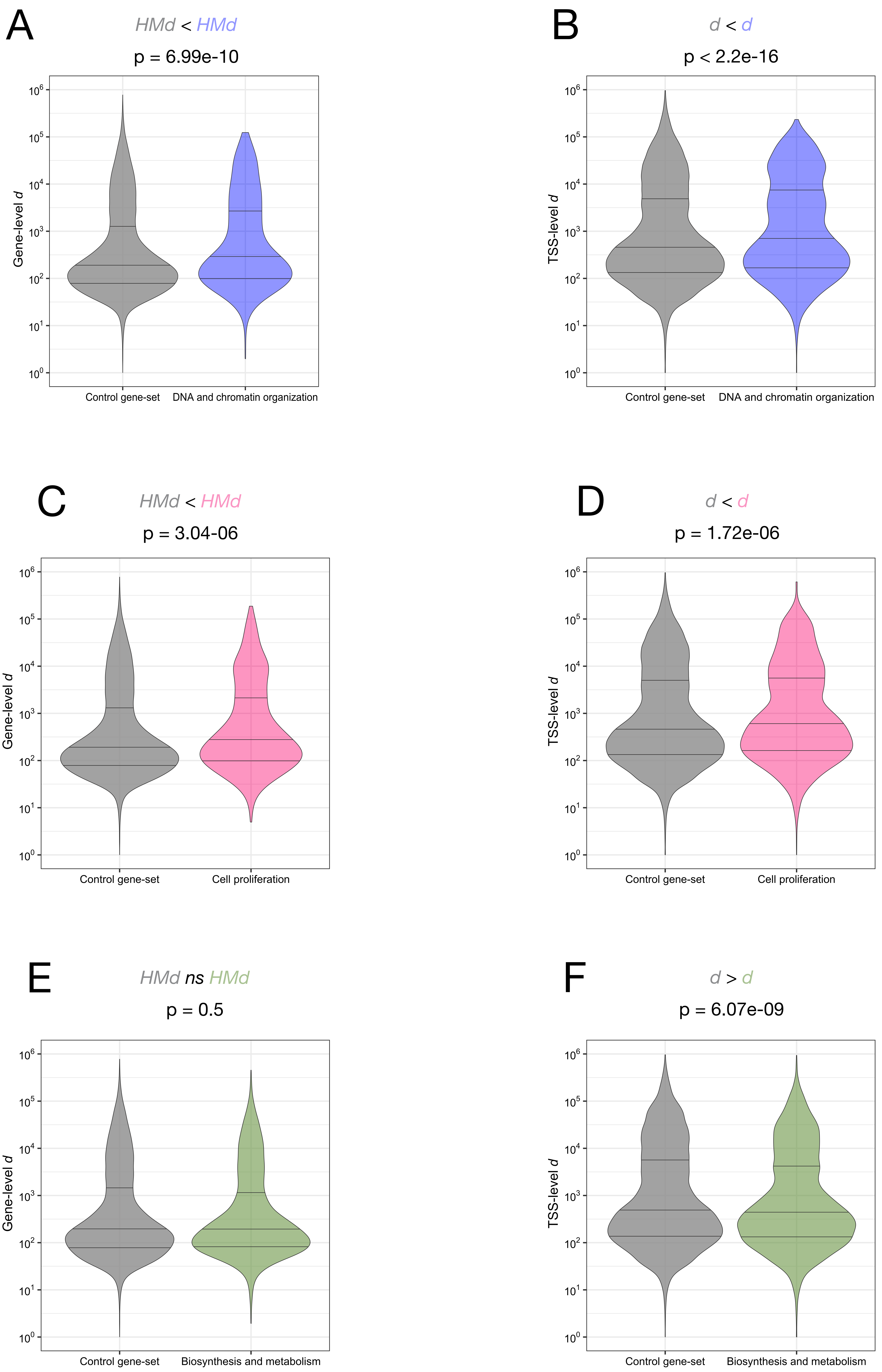
A
HMd < HMd
p = 6.99e-10
Gene-level d
Control gene-set
DNA and chromatin organization
B
d < d
p < 2.2e-16
TSS-level d
Control gene-set
DNA and chromatin organization
C
HMd < HMd
p = 3.04-06
Gene-level d
Control gene-set
Cell proliferation
D
d < d
p = 1.72e-06
TSS-level d
Control gene-set
Cell proliferation
E
HMd ns HMd
p = 0.5
Gene-level d
Control gene-set
Biosynthesis and metabolism
F
d > d
p = 6.07e-09
TSS-level d
Control gene-set
Biosynthesis and metabolism

### Fig S25

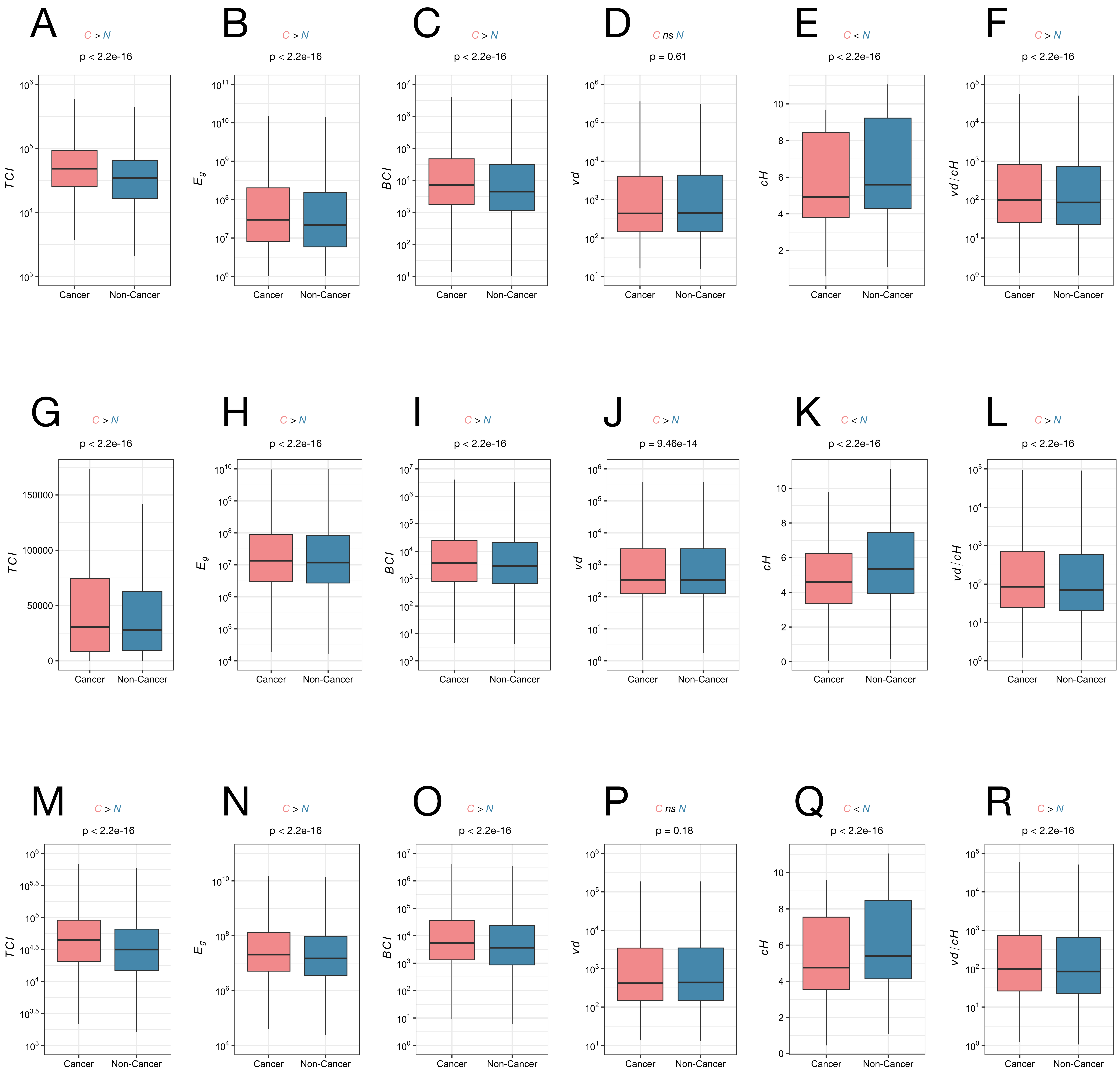
A
C > N
p < 2.2e-16
TCI
B
C > N
p < 2.2e-16
C
C > N
p < 2.2e-16
BCI
D
C ns N
p = 0.61
vd
E
C < N
p < 2.2e-16
cH
F
C > N
p < 2.2e-16
vd / cH
G
C > N
p < 2.2e-16
H
C > N
p < 2.2e-16
I
C > N
p < 2.2e-16
J
C > N
p = 9.46e-14
K
C < N
p < 2.2e-16
L
C > N
p < 2.2e-16
M
C > N
p < 2.2e-16
N
C > N
p < 2.2e-16
O
C > N
p < 2.2e-16
P
C ns N
p = 0.18
Q
C < N
p < 2.2e-16
R
C > N
p < 2.2e-16
Cancer
Non-Cancer

Fig S26

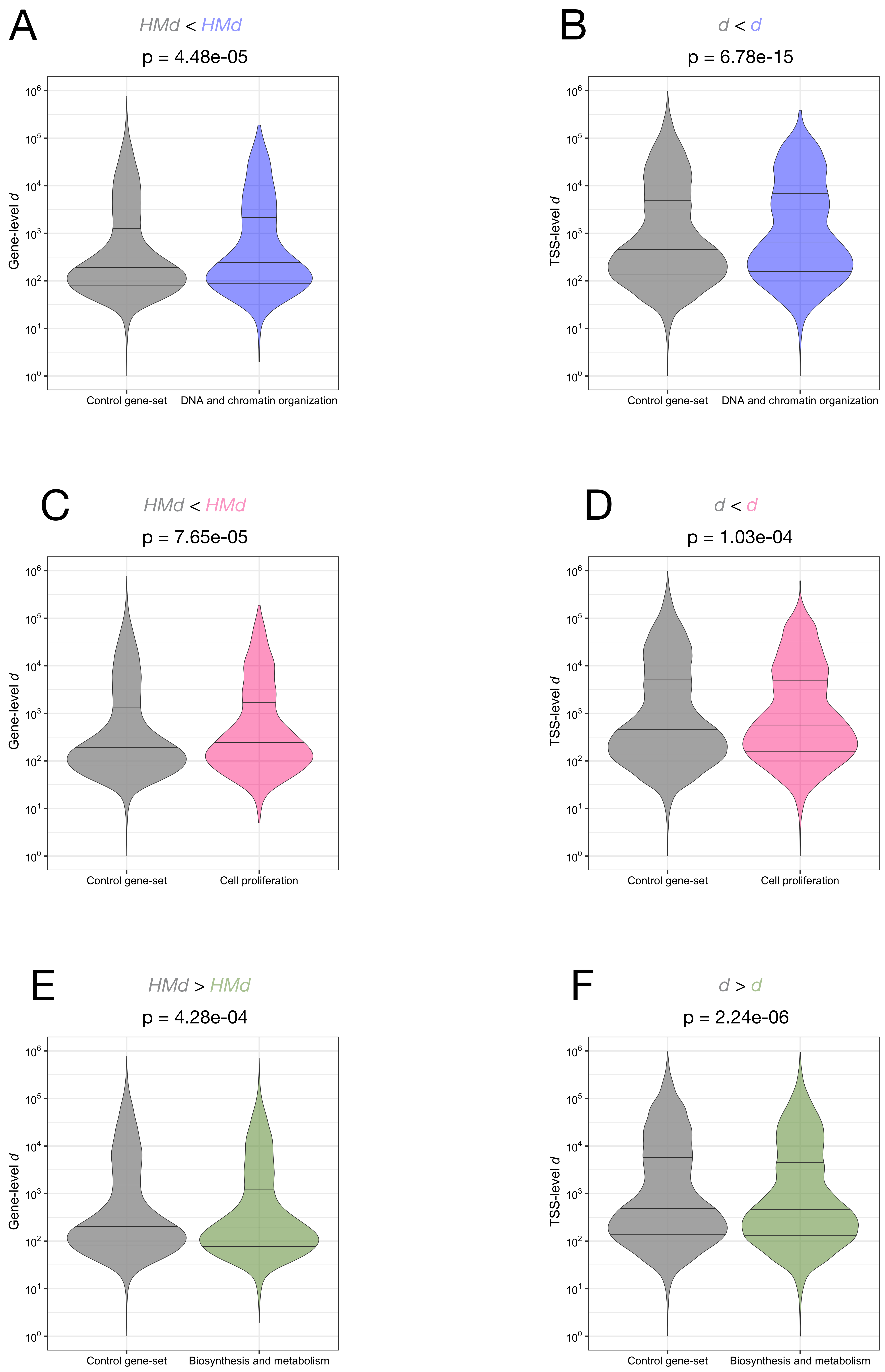
A
HMd < HMd
p = 4.48e-05
Gene-level d
Control gene-set
DNA and chromatin organization
B
d < d
p = 6.78e-15
TSS-level d
Control gene-set
DNA and chromatin organization
C
HMd < HMd
p = 7.65e-05
Gene-level d
Control gene-set
Cell proliferation
D
d < d
p = 1.03e-04
TSS-level d
Control gene-set
Cell proliferation
E
HMd > HMd
p = 4.28e-04
Gene-level d
Control gene-set
Biosynthesis and metabolism
F
d > d
p = 2.24e-06
TSS-level d
Control gene-set
Biosynthesis and metabolism
